# A Mechanistically Integrated Model of Exploitative and Interference Competition over a Single Resource Produces Widespread Coexistence

**DOI:** 10.1101/2023.09.27.559600

**Authors:** Daniel J.B. Smith, Joanna Masel

**Affiliations:** Department of Integrative Biology, University of Texas at Austin, Austin, TX, 78701, USA; Department of Ecology and Evolutionary Biology, University of Arizona, Tucson, Arizona 85721, USA

## Abstract

Many ecological models treat exploitative competition in isolation from interference competition. Corresponding theory centers around the *R*^∗^ rule, according to which consumers who share a single limiting resource cannot coexist. Here we model motile consumers that directly interfere while handling resources, mechanistically capturing both exploitative and interference competition. Our analytical coexistence conditions show that interference competition readily promotes coexistence. In contrast to previous theory, coexistence does not require intra-specific interference propensities to exceed inter-specific interference propensities, nor for interference behaviors to carry a direct (rather than merely an opportunity) cost. The underlying mechanism of coexistence can resemble the hawk-dove game, the dominance-discovery trade-off (akin to the competition-colonization trade-off), or a novel trade-off we call the “dove-discovery trade-off”, depending on parameter values. Competitive exclusion via the *R*^∗^ rule occurs only when differences in exploitative abilities swamp other differences between species, and more easily when differences in *R*^∗^ reflect different search speeds than when they reflect different handling times. Our model provides a mathematically tractable framework that integrates exploitative and interference competition, and synthesizes previous disparate models.

## INTRODUCTION

The *R*^∗^ rule (Armstrong and McGehee 1980; Tilman 1981; Tilman 1982) is a fundamental building block of theoretical ecology. Briefly, *R*^∗^ theory equates the competitive ability of a focal species or genotype with its ability to deplete a limiting resource. Consider one species consuming a single limiting resource. The consumer’s population grows until it depletes the resource concentration (*R*) down to the level (*R*^∗^) at which its growth stops. A species or genotype is predicted to exclude any competitors with higher *R*^∗^, because it drives the resource level too low for others to persist – this is referred to as the *R*^∗^ rule.

*R*^∗^ theory and the *R*^∗^ rule thus focus on *exploitative competition*, in which consumers compete indirectly through shared use of a depletable resource (either biotic or abiotic). This emphasis on exploitation has often neglected *interference competition*, i.e. direct interactions between individuals at the same trophic level, via behavioral contests or chemical inhibition. Interference competition is common in natural communities (Boynton 2019; DeLong and Vasseur 2011; Drury et al. 2020; García-Bayona and Comstock 2018; Grether et al. 2017; Inderjit et al. 2011; Leighton et al. 2024; Schoener 1983; Ziv et al. 1993; Ziv and Kotler 2003) but is rarely emphasized in coexistence theory. For example, otherwise wide-ranging reviews of coexistence theory, replete with discussions of resource exploitation/partitioning and predation, lack a single direct mention of interference competition (e.g. Adler et al. 2013; Barabás et al. 2018; Chesson 2018; Letten et al. 2017). While informal, this reflects a lack of emphasis on interference.

Several previous models that highlight interference competition use phenomenological terms to summarize the strengths of intra-specific (or intra-type) and inter-specific (or inter-type) interference (Case and Gilpin 1974; Hsu 1982; Kishi and Nakazawa 2013; Schoener 1976; Schoener 1978; Vance 1984). These studies broadly conclude that interference can promote coexistence if intra-specific interference exceeds inter-specific interference, but further insight is limited by the phenomenological nature of both terms. Moreover, these studies generally model resources implicitly or non-dynamically, making them incommensurable with *R*^∗^ theory.

Amarasekare (2002) incorporates both resource dynamics and interference competition to produce an *R*^∗^-related analysis of species coexistence. Amarasekare (2002) concludes that interference competition can only maintain coexistence when one consumer species engages in a interference mechanism that brings direct benefits to themselves, such as intraguild predation or parasitism, while the second consumer is superior at exploitation. Amarasekare (2002) explicitly argues that contests over resources that instead bear direct costs (for which she explicitly lists food items or territories as examples) cannot promote coexistence. Interpreting what is in its most direct effects a “beneficial” vs. a “costly” interference mechanism is made more difficult by the fact that Amarasekare (2002) continues the phenomenological tradition for modeling interference competition via Lotka-Volterra-like competition coefficients, on top of a typical consumer-resource model. We do, however, note that she models only inter-specific interference.

In contrast, in the hawk-dove game (Maynard Smith 1974; Maynard Smith 1982; Maynard Smith and Price 1973), interference does promote coexistence. “Doves” abstain from, while “hawks” participate in, contests (i.e., interference competition) for access to a shared, implicitly modeled resource. Obtaining a resource yields benefit *b*, and losing a contest yields cost *c*. When a hawk within a hawk-only population incurs more cost than benefit from interference (*c*/*b* > 1), a dove that forgoes interference can invade, enabling coexistence. This hawk-hawk interference does not occur in the model of Amarasekare (2002), which corresponds to modeling hawk-dove interactions but not hawk-hawk nor dove-dove interactions. We also note that hawk-dove coexistence does not require co-evolution as in Grether and Okamoto (2022) (i.e., hawks and doves can coexist when each of the two strategies is unchanging over time).

Surprisingly, the mechanistic insights from the hawk-dove game into how resource interference shapes coexistence have not been fully incorporated into resource competition theory. Several models incorporate hawk-dove-like dynamics into ecological models but, because they continue the hawk-dove tradition of treating resources implicitly (e.g., Auger et al. 1998; Moussaoui et al. 2014) cannot provide insights into resource competition. Other models have incorporated explicit consumer-resource dynamics into the hawk-dove game by having searching consumers interfere with resource handlers (Auger et al. 2006; Galanthay et al. 2023; Kooi 2015; Křivan et al. 2018). However, these models have so far neglected differences in exploitative abilities (e.g., search rates, handling times), and results were not contextualized within *R*^∗^ theory. Kuang (2003) and McPeek (2012) show that incorporating intra-specific density-dependent terms (akin to hawk-hawk interference) into a consumer-resource model can permit coexistence of species with different *R*^∗^s. However, the phenomenological nature of these terms decouples interference from resource use (unlike the hawk-dove game).

Here we consider mechanisms by which exploitation and interference interact. A consumer’s *R*^∗^ is affected by its search rate (how quickly it encounters resources) and handling time (how long it takes to utilize resources). Search and handling also affect interference over resources. Specifically, long handling provides more vulnerability to disruption by interference, and a faster searching consumer can initiate more contests.

Within ecological theory, the dominance-discovery trade-off offers mechanistic insights into how interference can facilitate coexistence. The theory, describing a trade-off between colonizing resource-rich patches (exploitation), and displacing heterospecifics from patches (interference), was developed for ants, and seems to be used only in that context (Adler et al. 2007; Davidson 1998; Fellers 1987; Holway 1999; Van Oudenhove et al. 2018). Unlike consumer-resource models with phenomenological interference interaction terms (e.g., Amarasekare 2002; Kuang et al. 2003) dominance-discovery models incorporate a mechanism of interference (resource use disruption). The better known competition-colonization trade-off (Hastings 1980; Horn and MacArthur 1972; Skellam 1951; Tilman 1994) is formally similar to the dominance-discovery trade-off, but inferior competitors are interpreted as being displaced via locally superior exploitation (i.e. a per-patch version of the *R*^∗^ rule), rather than by interference that disrupts resource use. Key results on the competition-colonization trade-off also apply to the dominance-discovery trade-off, because of the formal similarity between the models.

While both the dominance-discovery trade-off and the hawk-dove game describe interference/contests over a shared resource, they achieve coexistence in qualitatively different ways. In the hawk-dove game, interference bears a direct fitness cost (*c*) that trades off with the benefits of interference (*b*). Hawks and doves have identical exploitative ability (e.g. Galanthay et al. 2023; Maynard Smith 1982). In contrast, dominance-discovery has interference (contest-winning) ability trade off with exploitative (resource-discovery) ability (Adler et al. 2007; Van Oudenhove et al. 2018). Interference is not costly in these models beyond the trade-off in abilities. Rather, when one consumer encounters another handling a resource, a contest occurs, and the winner reaps the benefit of stealing the resource without incurring a potential cost. No existing theory unites the alternative interference-mediated coexistence mechanisms outlined by hawk-dove and dominance-discovery models.

We develop a model of motile consumers that mechanistically treats both exploitative and interference competition for an explicitly modeled resource. This unifies a continuum of mechanisms that, depending on parameter values, may resemble the *R*^∗^ rule, the hawk-dove game, the dominance-discovery trade-off, or a novel coexistence mechanism that we call the “dove-discovery” trade-off. We explore which parameter values lead to which regime, when interference-exploitation trade-offs enable coexistence, and whether the *R*^∗^ ratio for competing species (or genotypes) is a reliable metric of competitive success given the possibility of interference.

## METHODS

### Model

We extend a typical consumer-resource model to incorporate consumer-consumer interference competition over a single shared resource. We consider the following set of differential equations for a two consumer system:

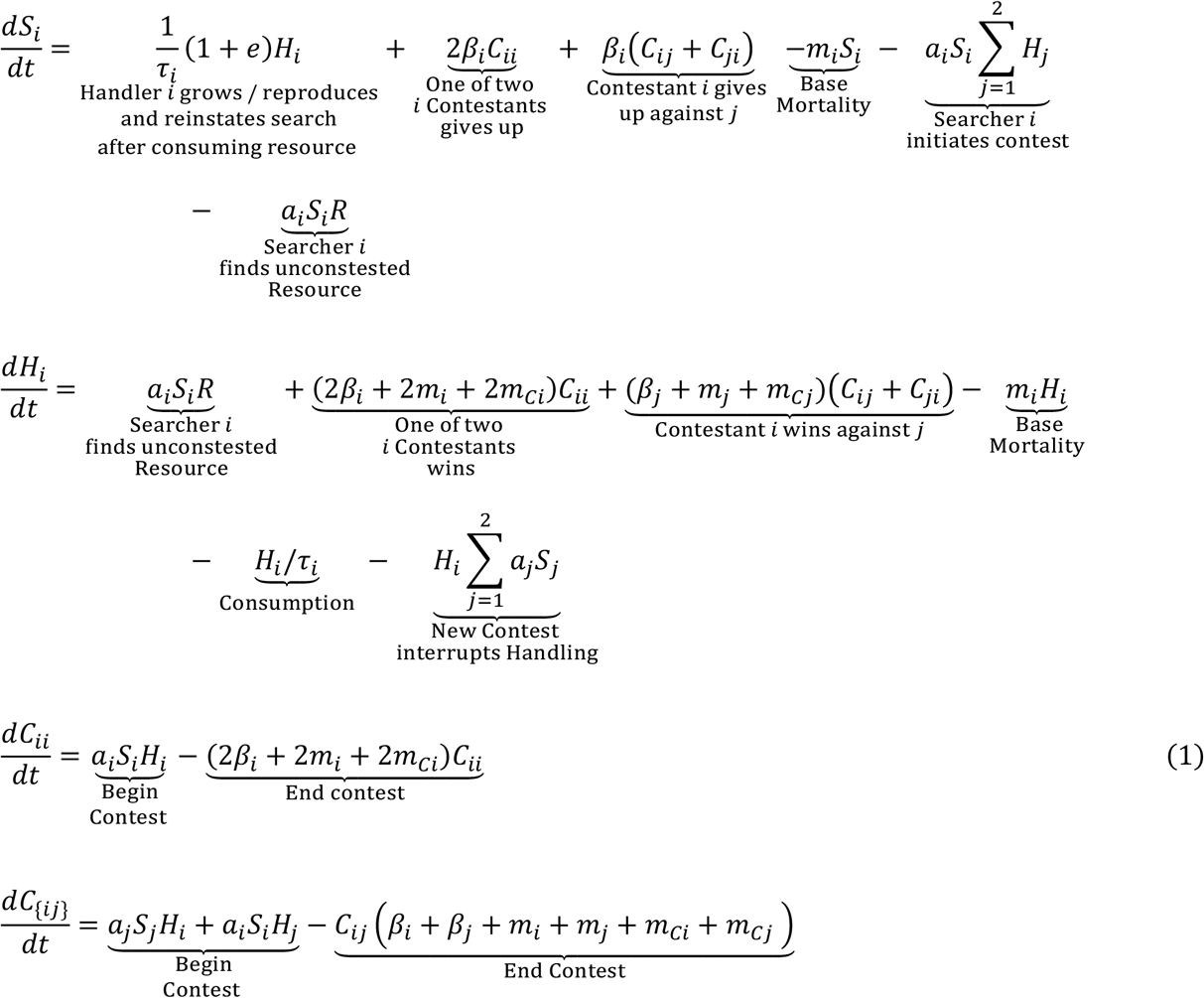

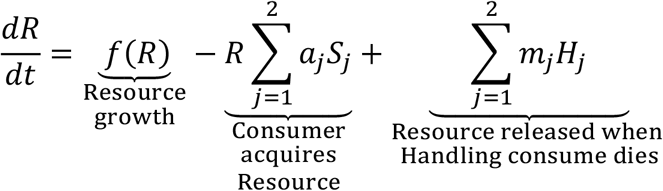

where *i* = 1,2, *j* = 1,2, *i* ≠ *j*. While seemingly complex, the system of equations in (1) straightforwardly describes mass action kinetics (i.e. approximating exponential distributions of waiting times by using continuous, additive rates) that are most easily understood through inspection of Fig. 1. Consumer types *i* and *j* could either be different species, or different variants of a single polymorphic species.

**Fig. 1:**
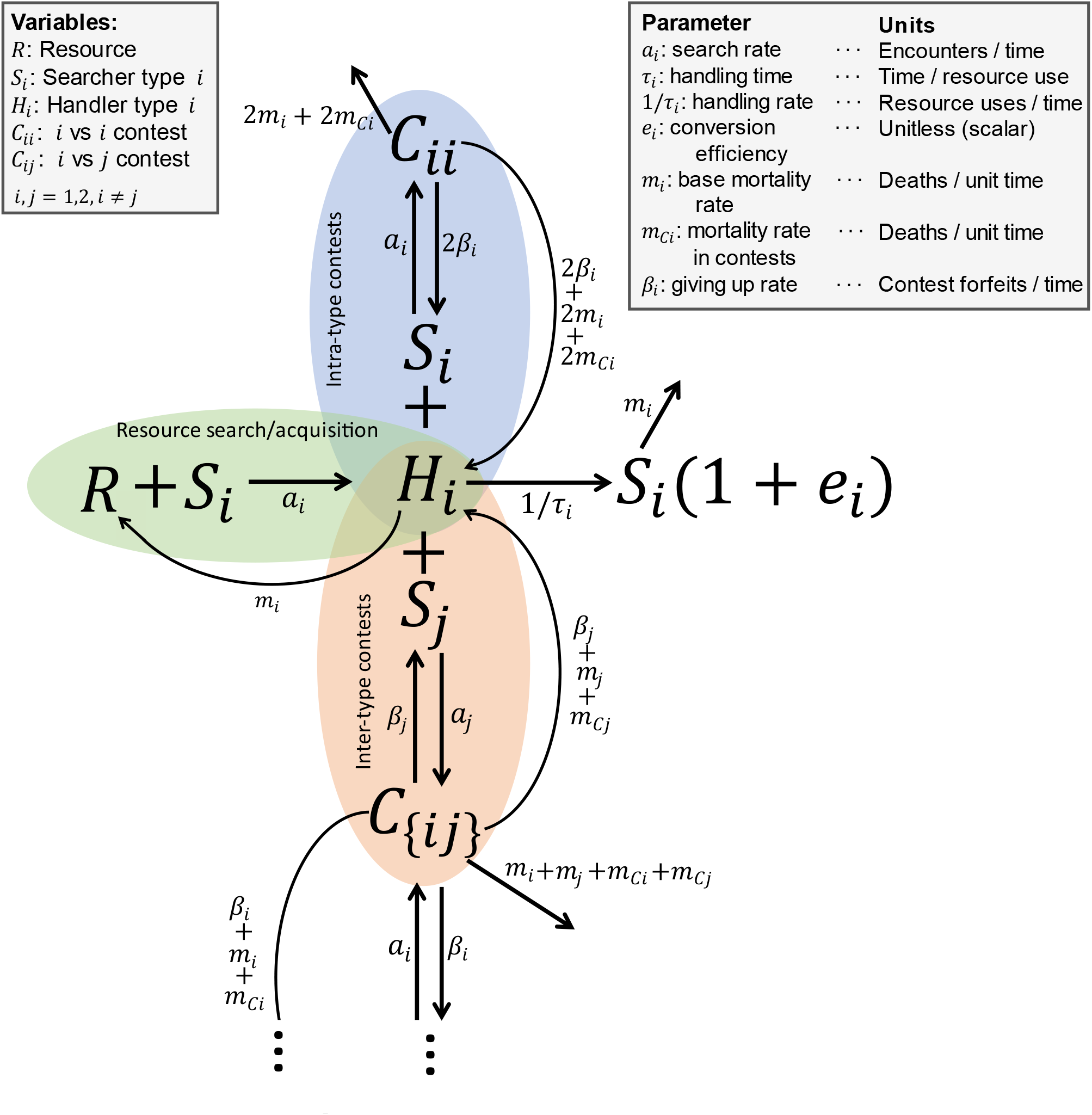
Model schema. Variables are in upper case, parameters in lower. Full dynamics of consumer *i* are shown; those of consumer *j* are symmetric and truncated from the lower portion of the figure for simplicity; Supplementary Figure I1 is not truncated. In green, <u>R</u>esources and type *i* <u>S</u>earchers (*R* + *S*_*i*_) combine to form a <u>H</u>andler (*H*_*i*_). <u>H</u>andlers of type *i* combine with <u>S</u>earchers (*S*_*i*_ or *S*_*j*_) to form *C*_*ii*_ (blue) or *C*_{*ij*}_ = *C*_*ij*_ +*C*_*ji*_ (peach) <u>C</u>ontesting pairs, respectively. Rates are denoted as search *a*, base mortality *m*, additional mortality from contests *m*_*C*_, giving up on contests *β*, and handling 1/*τ*.

Exploitation is shown in the green/left oval of Figure 1. Searching consumers *S*_*i*_ of type *i* find Resources *R* at rate *a*_*i*_ and begin to handle them once encountered. Handlers *H*_*i*_ consume resources at rate 1/*τ*_*i*_ (Figure 1, right). We refer to *τ* as “handling time,” although we note that it differs from the transformed handling time (*h*) traditionally used in ecological models with a type II functional response (*aR*/(1 + *ahR*); Holling 1959) where *h* is a composite parameter formally derived by splitting consumers into searching and handling state variables and performing a separation of timescales (Huisman and De Boer 1997; Real 1977). Reproduction is proportional to resource consumption *e*_*i*_*H*_*i*_/*τ*_*i*_, with *e*_*i*_ as a conversion factor.

Consumers die at base rate *m*_*i*_, elevated by *m*_*Ci*_ during contests (see below). We treat resources as discrete, such that *τ* specifies the probability that a resource is 0% vs 100% consumed, rather than the degree to which a resource is consumed. When a consumer dies while handling a resource, the resource returns to the resource pool (i.e., at rate *m*_*i*_*H*_*i*_; Fig. 1, curved arrow below green/left oval).

Interference is shown in the blue and peach ovals of Figure 1. Contests begin when a Searcher encounters a Handler. *C*_*ii*_ represents the number of within-type contests of type *i* (Fig. 1, blue), and *C*_{*ij*}_ = *C*_*ij*_ +*C*_*ji*_ (*j* ≠ *i*) represents the number of between-type contests (Fig. 1, peach). Future extensions might distinguish *C*_*ij*_, in which type *i* found the resource first, from *C*_*ji*_, but we assume they are equivalent. Conflicts resolve either when a consumer gives up at rate *β*_*i*_, or dies at rate *m*_*i*_ + *m*_*Ci*_. Because each unit of *C*_*ii*_ contains two individuals of type *i*, victors re-enter the handling state at rate (2*β*_*i*_ + 2*m*_*i*_ + 2*m*_*Ci*_)*C*_*ii*_. The losers similarly re-enter the searching state at rate 2*β*_*i*_*C*_*ii*_. *C*_{*ij*}_ resolves into a type *i* victory at rate (*β*_*j*_ + *m*_*j*_ + *m*_*Cj*_)*C*_{*ij*}_, with losers of type *i* re-entering the searching state at rate *β*_*i*_*C*_{*ij*}_.

Intra-type interference propensity would exceed inter-type interference if type *i* gave up more slowly against *i* than against *j*. We avoid this assumption, which is known to easily produce coexistence (Vance 1984), by making giving up rates *β*_*i*_ depend only on the focal type *i* and not on the opponent type *j*. Similarly, contest initiation occurs whenever a Searcher encounters a Handler, regardless of their type.

Resource *R* grows according to a saturating function *f*(*R*). We assume *f*(*R*) = *r*(*K* − *R*), representing a spontaneously regenerating consumable resource. We consider this functional form because it avoids limit cycles in the 1-consumer-1-resource case (Supplementary Materials A), which is not the case for a biological resource exhibiting logistic growth (i.e. *f*(*R*) = *rR*(1 − *R*/*K*)) (Johnson and Amarasekare 2015; Rosenzweig 1971). The lack of oscillations simplifies our coexistence analyses, because oscillations driven by nonlinear functional responses (i.e., handling time) can affect species coexistence (Armstrong and McGehee 1980). However, numerical simulations suggest that results are not qualitatively different with logistic resource growth (Supplement Materials B).

Our model (1) has similarities to several previous works. Kooi (2015) and Galanthay et al. (2023) use a similar set of equations to model hawk-dove-like dynamics, in the special case where types *i* and *j* have identical exploitative ability (*a, τ*, etc.). Adler et al. (2007)’s model of the dominance-discovery trade-off corresponds to the limit of fast contests in our model, which are not explicitly modeled in theirs, with outcomes instead resolved as soon as one consumer encounters another.

### Analysis

In the Supplementary Materials Sections D, F, and G, we solve for invasion criteria (Turelli 1978) for the two-consumer one-resource system described by equations in (1). To do so, we first solve the non-trivial equilibrium for each one-consumer one-resource system, yielding 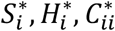 and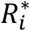. We derive the 4 × 4 Jacobian matrix for the variables *S*_*j*_, *H*_*j*_, *C*_*jj*_, and *C*_{*ij*}_ where *j* ≠ *i*. We then compute the eigenvalues of the Jacobian matrix at consumer *i*’s non-trivial equilibrium when consumer *j* is a rare invader: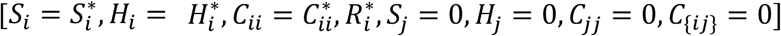. We derive the conditions for invasion and coexistence both analytically and numerically using Mathematica version 13.2. Consumer *j* will increase in abundance (invade) when rare if and only if at least one eigenvalue is positive. Coexistence is inferred from mutual invasibility, which we validate using numerical simulations of the system of differential equations in (1) (Supplementary Materials Section C).

## RESULTS

### R^∗^ rule in the absence of interference competition

The *R*^∗^ rule is a limiting case of our model, which occurs when we remove interference entirely. When a Searcher who encounters a Handler does not contest the resource, the system of equations reduces to

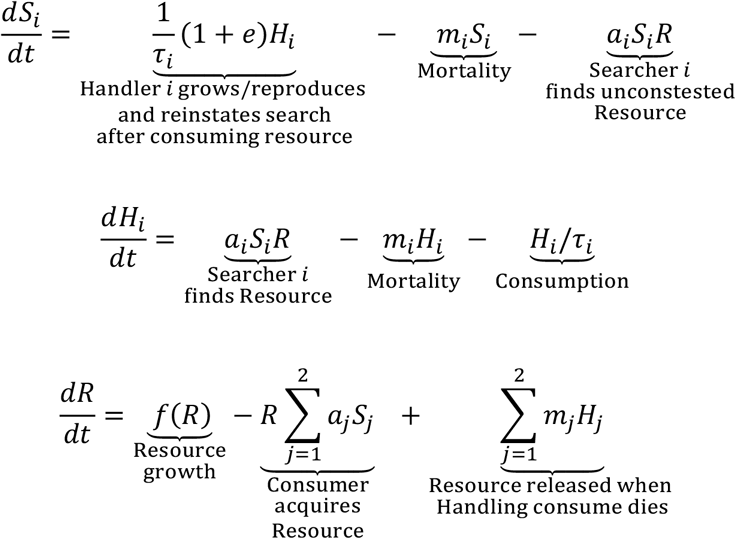

Let 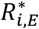 denote the corresponding equilibrium resource abundance set by type *i* in isolation, in this scenario when only *<u>E</u>xploitative* competition occurs. Assuming population viability 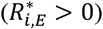:

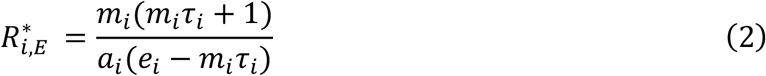

(Supplementary Materials Section D.3) which is biologically feasible when *e*_*i*_ > *τ*_*i*_*m*_*i*_ and 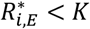. We refer to 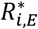 as the *exploitative ability* of consumer *i*, and to 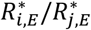 as the *relative exploitative ability* of *i* and *j*. 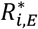 also represents the minimum resource requirement for consumer *i*.

Type *i* can invade a population of type *j* if and only if 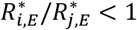 (Supplementary Materials Section D.4), and so mutual invasibility requires the unfulfillable condition:

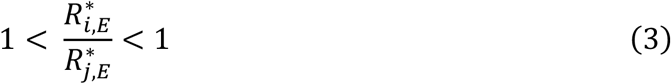

This is the *R*^∗^ rule of Tilman (1982). Neutral coexistence is possible in the knife edge case of 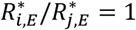. Equation (3) can be compared to our more general Equation (11).

### Interference increases equilibrium resource density

Interference competition increases the equilibrium resource density set by a consumer, setting the stage for deviations from the *R*^∗^ rule. We denote the equilibrium resource density set by consumer *i* in the 1-consumer-1-resource system when *<u>I</u>nterference* occurs as (Supplementary Materials Section E):

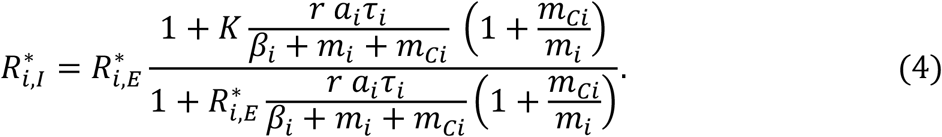

Because 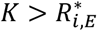, Equation (4) implies 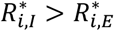, with 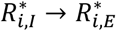 as *β*_*i*_ → ∞ (contests end immediately, so they do not affect dynamics) or as *τ*_*i*_ → 0 (handling time is arbitrarily short, so consumers have no opportunity to begin contests). 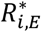 continues to describe the minimum resource requirement for *i*: interference thus decouples a consumer’s resource requirement from the resource abundance it sets at equilibrium.

There are two reasons why interference increases equilibrium resource density. First, conflict-induced mortality *m*_*Ci*_ directly removes consumers, allowing the resource to grow. The direct cost of contests *c* in the classic hawk-dove game can readily be interpreted in terms of a probability of death per contest 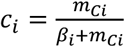 as *m* _*Ci*_, *β*_*i*_ → ∞ (contests are very fast and lethal). In this limit, equation (4) becomes:

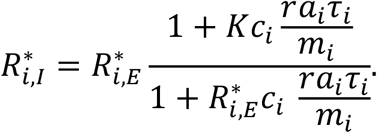

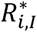 increases as the probability of contest mortality *c*_*i*_ increases.

Second, engaging in conflicts incurs an opportunity cost, because time spent in a conflict is not available for searching or handling resources, allowing the resource to grow. Consequently, 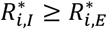 even if contests do not increase death rates (substitute *m*_*Ci*_ = 0 into Equation (4)). The extent to which opportunity costs increase resource density depends on how the time spent in contests compares to the rate at which new resources appear. Opportunity costs are large when contests begin frequently (large *a*_*i*_*τ*_*i*_), when contests are long (small *β*_*i*_) and when resources renew rapidly (large *r*; see Equation (4)). Below, we emphasize opportunity costs only (by assuming *m*_*Ci*_ = 0), but the same qualitative results apply for direct costs.

### Classifying outcomes when only one type interferes

Our main goal is to unify previously disparate mechanisms of interference and exploitative competition over resources. To this end, we first consider a simplified version of the set of equations in (1), with limiting cases that resemble the hawk-dove game and the dominance-discovery trade-off. We consider two types of consumers: (1) a “hawk” *h* that contests resources (*β*_*h*_ < ∞) and (2) a “dove” *d* that does not (*β*_*d*_ = ∞), replacing our previous, more general notation of *i* and *j*. In later sections, we will consider the more general asymmetric case of *β*_*h*_ < *β*_*d*_ ≤ ∞. Note that the *h* indicating a “<u>h</u>awk” should not be confused with the *H* indicating “Handlers”.

Doves can invade a community of hawks if:

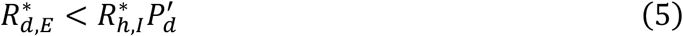

where

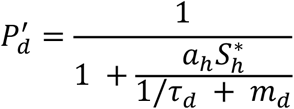

is the probability that a dove does not experience hawk interference during handling (Supplementary Materials Sections F.4.1 and F.5). Equation (5) represents the fact that doves must be able to persist at a resource density 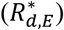 not just below the equilibrium set by the hawks 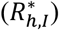, but below a lower level that takes into account doves’ inability to benefit from resources that they relinquish to hawks 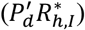. Note that from *β*_*d*_ = ∞, we have 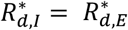 (doves give up immediately, so dove-dove interference does not increase the equilibrium resource density), in contrast to 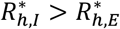 (because *β*_*h*_ < ∞, hawk-hawk interference increases the equilibrium resource density). Hawk interference generates both a benefit 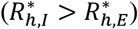 and a cost 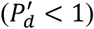 to dove invasion.

Hawks can invade a resident dove population if:

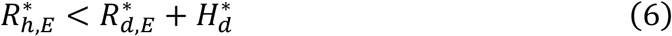

(Supplementary Materials Section F.4.2). Equation (6) represents the fact that hawks must be able to persist on resources that include both those currently unclaimed by doves 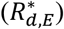, and those currently held by doves that hawks can readily steal 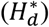.

The two invasion criteria can be rearranged to form the mutual invasion criterion

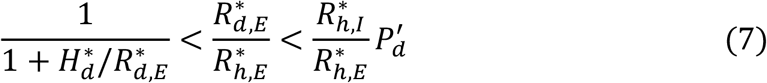

(Supplementary Materials Section F.4.3), which implies coexistence when fulfilled. Equation (7) can be compared to Equation (3) describing exploitation alone.

#### Connection to the Classic Hawk-Dove Game

The traditional hawk-dove game is akin to a special case of our model, when hawks and doves have identical exploitative ability 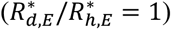. In this scenario, the hawk can always invade a dove population (from Equations (6) and (7)). The dove can also invade a pure hawk population, implying coexistence, when:

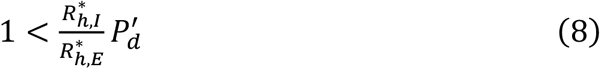

Akin to the classic hawk-dove game, the dove invades if the cost associated with hawk-hawk interference 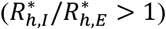 more than compensates for hawk-dove resource theft 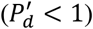. Hawk-dove coexistence, if it occurs, thus stems from an “intra-type interference – inter-type interference trade-off”. Hawks benefit from stealing from doves (inter-type interference). Doves invade due to the increase in resource density caused by hawk-hawk (intra-type) interference.

#### Connection to the Dominance-Discovery Trade-off

In dominance-discovery models, contests resolve instantaneously and are not directly costly. Our model can capture this by letting hawks finish contests arbitrarily fast (*β*_*h*_ → ∞), while always defeating doves (*β*_*h*_/*β*_*d*_ = 0). This yields 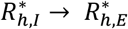. The coexistence criteria then become

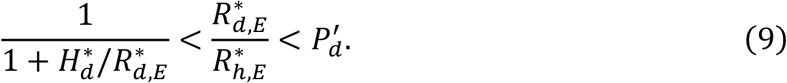

The dove then needs to be superior at exploitation 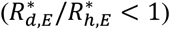 to invade (because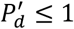), offsetting the disadvantage of losing contests. The resemblance of this model limit to the dominance-discovery trade-off is most intuitive when the dove’s superior exploitation comes from a higher search rate (*a*). The condition for dove invasion is then 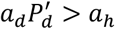, i.e. faster resource discovery of doves must more than compensate for theft by hawks. However, dove invasion can occur via any exploitative parameter(s) (*a, τ, m*, or *e*). Hawks can invade if there are enough doves handling resources 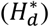 that can be stolen, to compensate for hawks’ inferior exploitative ability. Coexistence, if it occurs, thus stems from an “exploitation – inter-type-interference trade-off”. For simplicity, and given complications when multiple parameters differ between types, we refer to all such cases as dominance-discovery, although we note that dove invasion driven by *τ* may be reasonably thought of as a dominance-handling trade-off.

The dominance-discovery trade-off and hawk-dove coexistence differ with respect to how doves invade. With hawk-dove coexistence, doves invade by taking advantage of increased resource density due to hawk-hawk interference 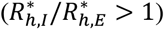. With the dominance-discovery trade-off, doves invade because of superior exploitative ability 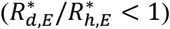. Note each dove invasion criterion depends on the ratio of different *R*^∗^s.

### Classifying outcomes when both types interfere

We generalize our above results by analyzing our model when both consumer types interfere.

#### General Invasion Criteria

When both *β*_*i*_ and *β*_*j*_ < ∞, we refer to the consumer with the smaller *β* as the hawk and that with the larger *β* as the dove based on their propensities to engage in interference contests, although both are on a spectrum of hawkishness and dovishness. Generalizing equations (5) and (6), *j* can invade *i* if and only if:

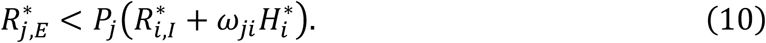

(see Supplementary Materials Section G.3 for derivation). As before, 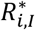 is the resource availability set by consumer *i*, including intra-type-*i* interference effects. The new term *ω*_*ji*_ is the probability that invader *j* wins a contest against resident *i*:

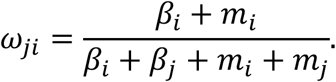

(see Supplementary Section Materials G.5). 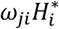 represents resources that invader *j* steals from handling residents of type *i*, i.e. type *j*’s benefit from inter-type interference. 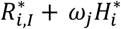 is the total resource available to *j*.

*P*_*j*_ is a generalization of 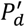 from equation (5). *P*_*j*_ quantifies the negative impact that inter-type interference has on the invasion of type *j* (0 ≤ *P*_*j*_ ≤ 1) where

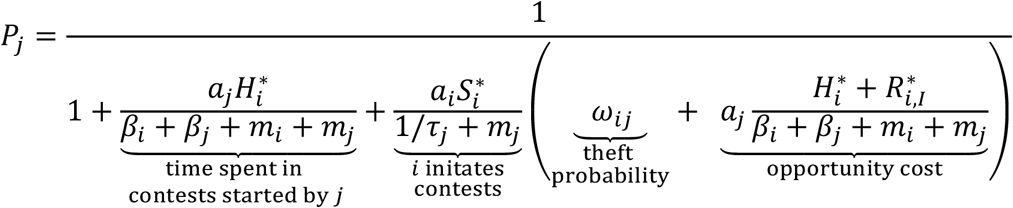

See Supplementary Materials Section G.3 for a derivation.

The first non-unity term in the denominator of *P*_*j*_ captures the opportunity cost from contests initiated by type *j*. The second term reflects costs from contests initiated by type *i* against *j*. Costs are larger when resource theft by *i* is more likely (larger *ω*_*ij*_) and when opportunity costs of participation are greater. Opportunity costs increase with the resource encounter rate, 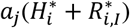. This illustrates the key tension of hawkish behavior: giving up slower (smaller *β*_*j*_) decreases the probability of losing contests but increases opportunity costs. Short handling time reduces the cost of interference; if type *j* rapidly handles resources (*τ*_*j*_ → 0), *i* cannot initiate contests with *j* and the second term in *P*_*j*_ disappears 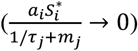.

#### Classifying coexistence and exclusion outcomes

Rearranging Equation (10) for each type (*i* and *j*) in Supplementary Material G.4, we derive criteria for coexistence in terms of how invaders *j* and *i* experience interference competition (*I*_*ji*_ and *I*_*ij*_, respectively), and of relative exploitative competition ability *E*_*ji*_:

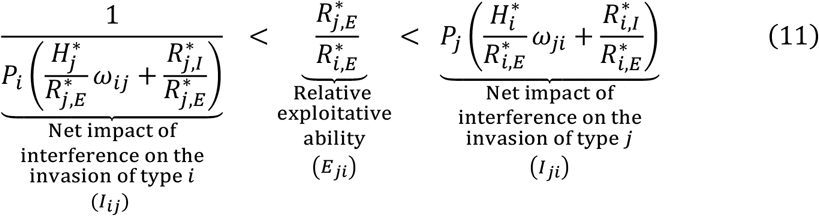

*E*_*ji*_ quantifies the exploitative ability of type *j* relative to type *i*, where *E*_*ji*_ < 1 indicates *j* is the superior exploitative competitor (*j* has a lower 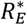). Recall that we refer to “exploitative ability” and “exploitative competition” purely in terms of 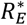 (and, thus *E*_*ji*_). *I*_*ij*_ quantifies the effect interference type *i* experiences while invading *j* (and vice versa for *I*_*ji*_). *i* invades if the left-hand side inequality is fulfilled (1/*I*_*ij*_ < *E*_*ji*_) and *j* invades if the right-hand side inequality is fulfilled (*E*_*ji*_ < *I*_*ji*_). Equation (11) reduces to Equation (7) if *j* does not interfere (*β*_*j*_ → ∞) and further reduces to the *R*^∗^ rule (equation (3)) in the absence of interference (for which *I*_*ij*_ = *I*_*ji*_ = 1). The interference terms (*I*_*ij*_, *I*_*ji*_) permit deviations from the *R*^∗^ rule and quantify the impact interference competition has on invasion.

We quantify the relative importance of interference competition (*I*_*ij*_, *I*_*ji*_) and relative exploitative ability (*E*_*ji*_) in determining outcomes by comparing their proportional deviations from unity (1). For example, consider when type *j* invades with *E*_*ji*_ < *I*_*ji*_. This may occur because *E*_*ji*_ < 1 (*j* is a superior exploitative competitor) and/or because *I*_*ji*_ > 1 (*j* experiences a benefit from interference). When both inequalities are true, this indicates that they both contribute, and we determine which of them predominantly causes invasion by comparing the extent to which *E*_*ji*_ vs. 1/*I*_*ji*_ proportionally deviates from 1 *E*_*ji*_ vs. 1/*I*_*ji*_. If *E*_*ji*_ < 1/*I*_*ji*_, then the benefit *j* experiences from superior exploitative ability predominantly allows invasion. Conversely, if *E*_*ji*_ > 1/*I*_*ji*_, interference contributes more to *j*′*s* invasion.

Similarly, the failure of *j* to invade with *E*_*ji*_ > *I*_*ji*_ may be due to *E*_*ji*_ > 1 (*j* is a poorer exploitative competitor) and/or to *I*_*ji*_ < 1 (*j* experiences a negative effect from interference). When both inequalities are true, indicating that both contribute, we say that exploitative disadvantage predominantly prevents invasion when *E*_*ji*_ > 1/*I*_*ji*_ while interference predominantly prevents invasion when *E*_*ji*_ < 1/*I*_*ji*_.

We again substitute *h* (hawk) and *d* (dove) for *i* and *j*, where *β*_*h*_ < *β*_*d*_. We compare the magnitudes of *I*_*hd*_, *E*_*dh*_, and *I*_*dh*_ to distinguish six possible scenarios summarized in Table 1. For notational convenience, we also refer to *E*_*hd*_ = 1/*E*_*dh*_ below.

**Table 1:**
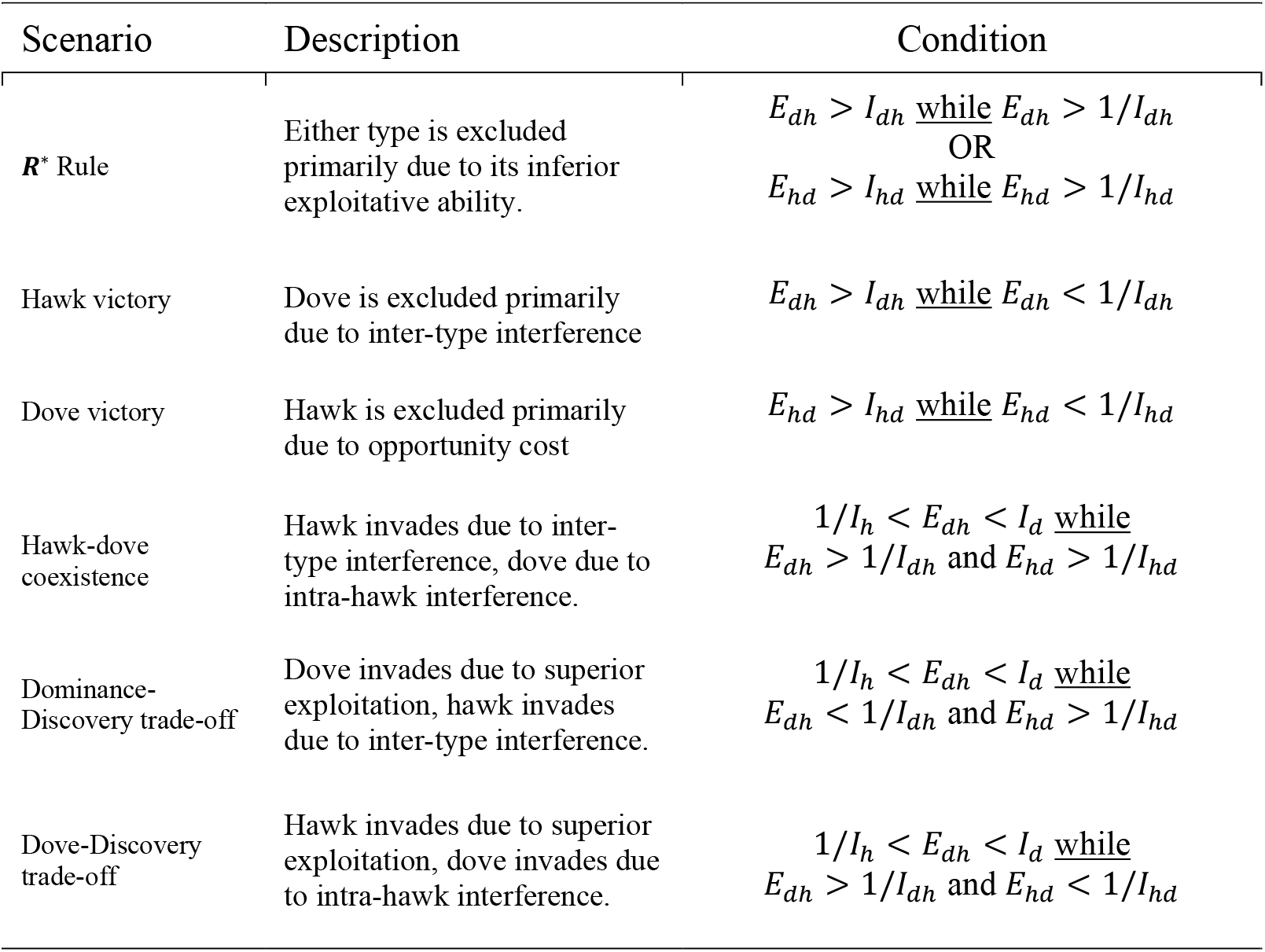
Summary of the different scenarios, associated criteria for invasion or lack of invasion, and the conditions that further distinguish scenarios beyond their invasion criteria. *h* indexes the “hawk” and *d* the “dove” on the basis of *β*_*d*_ > *β*_*h*_. Each scenario refers to the *predominant* mechanism underlying the outcome. Expressions come from Equation (11). Note that *E*_*hd*_ = 1/*E*_*dh*_.

| Scenario | Description | Condition |
| --- | --- | --- |
| $R^*$ Rule | Either type is excluded primarily due to its inferior exploitative ability. | $E_{dh} > I_{dh}$ <u>while</u> $E_{dh} > 1/I_{dh}$<br>OR<br>$E_{hd} > I_{hd}$ <u>while</u> $E_{hd} > 1/I_{hd}$ |
| Hawk victory | Dove is excluded primarily due to inter-type interference | $E_{dh} > I_{dh}$ <u>while</u> $E_{dh} < 1/I_{dh}$ |
| Dove victory | Hawk is excluded primarily due to opportunity cost | $E_{hd} > I_{hd}$ <u>while</u> $E_{hd} < 1/I_{hd}$ |
| Hawk-dove coexistence | Hawk invades due to inter-type interference, dove due to intra-hawk interference. | $1/I_h < E_{dh} < I_d$ <u>while</u><br>$E_{dh} > 1/I_{dh}$ and $E_{hd} > 1/I_{hd}$ |
| Dominance-Discovery trade-off | Dove invades due to superior exploitation, hawk invades due to inter-type interference. | $1/I_h < E_{dh} < I_d$ <u>while</u><br>$E_{dh} < 1/I_{dh}$ and $E_{hd} > 1/I_{hd}$ |
| Dove-Discovery trade-off | Hawk invades due to superior exploitation, dove invades due to intra-hawk interference. | $1/I_h < E_{dh} < I_d$ <u>while</u><br>$E_{dh} > 1/I_{dh}$ and $E_{hd} < 1/I_{hd}$ |

Firstly, differences in exploitative ability may be predominantly responsible for competitive exclusion of a consumer (the dove or hawk is excluded while *E*_*dh*_ > 1/*I*_*dh*_ or *E*_*hd*_ > 1/*I*_*hd*_, respectively). This scenario resembles the *R*^∗^ rule because the exploitative 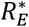 ratio predominantly underlies exclusion.

Secondly, interference competition can cause competitive exclusion. This can occur via two mechanisms. (1) “Hawk victory” in which hawks exclude doves via inter-type interference when *E*_*dh*_ < 1/*I*_*dh*_. (2) “Dove victory” in which doves exclude hawks when the opportunity costs from interference are too large, when *E*_*hd*_ < 1/*I*_*hd*_. Unlike the simplified case of *β*_*h*_ = ∞, the hawk may be excluded even if the dove is not superior at exploitation. It is therefore possible for a consumer who is both superior at exploitation and more likely to win contests to be competitively excluded.

Coexistence can occur via three mechanisms. Firstly, “hawk-dove coexistence” describes when interference considerations enable invasion of both types: *E*_*dh*_ > 1/*I*_*dh*_ and *E*_*hd*_ > 1/*I*_*hd*_. Those considerations are the hawk’s gain from stealing (intra-type interference), and the dove’s gain from hawk-hawk contests (intra-type interference), which exceed any benefit of superior exploitative ability. This generalizes the inter-type vs intra-type interference trade-off beyond the previous section’s treatment of *β*_*d*_ = ∞.

Coexistence can also occur via a trade-off between exploitative and interference abilities. This trade-off occurs via two distinct mechanisms. First, doves invade primarily due to superior exploitative ability (*E*_*dh*_ < 1/*I*_*dh*_) while hawks invade predominantly due to interference effects, meaning stealing resources from doves (*E*_*hd*_ > 1/*I*_*hd*_). This scenario resembles the dominance-discovery trade-off, generalizing our treatment above of the *β*_*d*_ = ∞ case. This scenario can also be thought of as a “hawk-discovery” trade-off.

The direction of the exploitative vs. interference trade-off is reversed in the second mechanism: doves primarily invade due to the increase in resource density caused by hawk-hawk interference (*E*_*dh*_ > 1/*I*_*dh*_), while hawks invade primarily due to superior exploitative ability (*E*_*hd*_ < 1/*I*_*hd*_). This constitutes a novel intra-type-interference vs. exploitation trade-off, which we call the “dove-discovery trade-off”.

Finally, a priority effect (founder control) occurs when 1/*I*_*hd*_ > *E*_*dh*_ > *I*_*dh*_. However, we do not discuss this scenario in detail, nor include it in Table 1 because priority effects require unlikely parameter value choices in our model (Supplementary Materials Section H).

#### Outcome classification based on mechanistic parameters

*I*_*dh*_, *I*_*hd*_, and *E*_*dh*_ (Table 1) are not independent: changes to a single model parameter alter multiple key properties (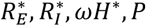; Equations (10-11)). We next dig deeper into the causal effects of mechanistic parameters on competitive outcomes and which scenario causes them.

First, consider the relatively simple case of hawk and dove equal in all parameters except giving up rate, *β*, resulting in equal exploitative abilities. Hawk-dove coexistence (Table 1) occurs if and only if *β*_*h*_ and *β*_*d*_ are on either side of a central value (henceforth *β*_0_; Fig. 2, yellow star). *β*_0_ is a complicated (though mathematically tractable) expression that depends on every parameter of the model (Supplementary Materials Section G.6).

**Fig. 2:**
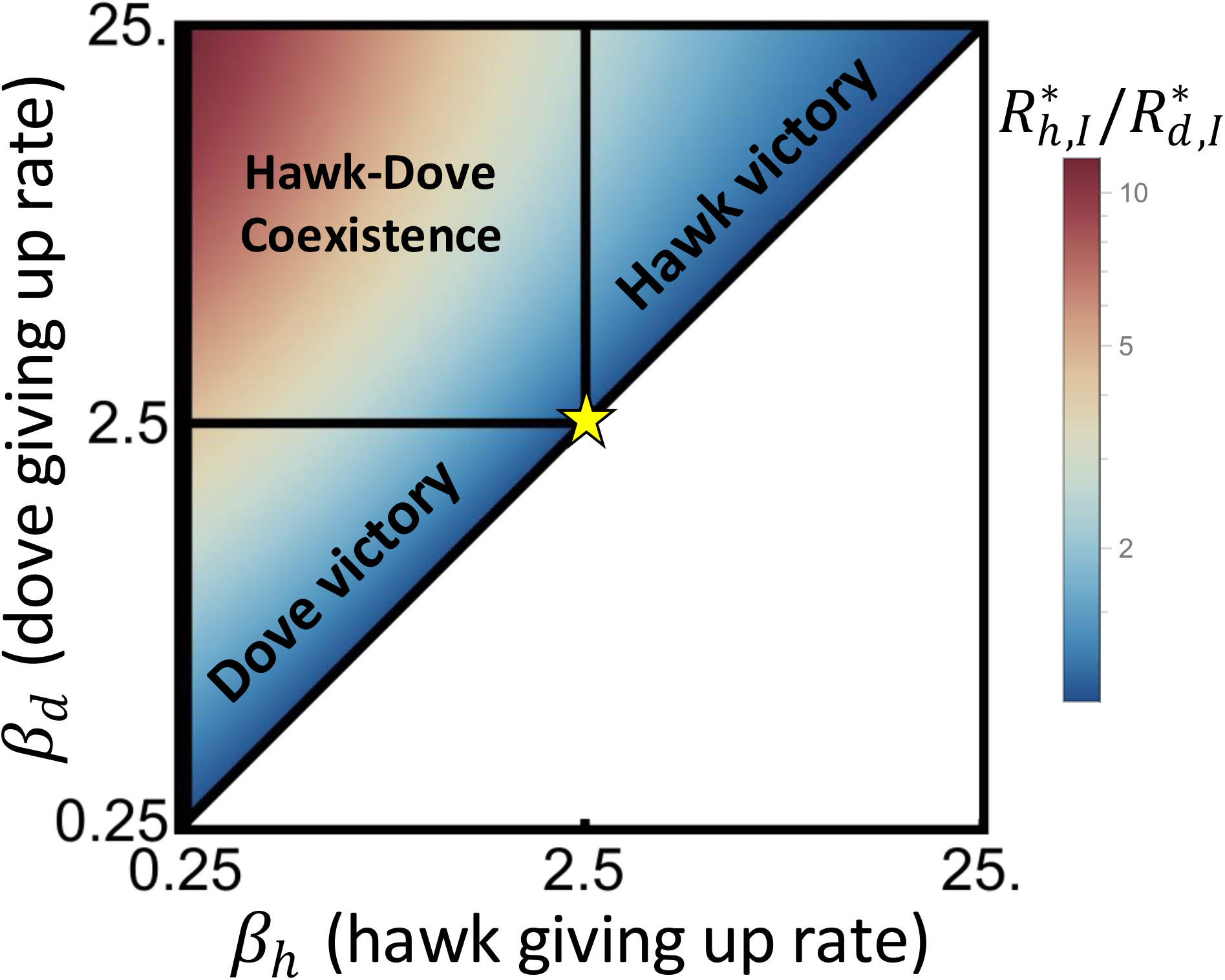
The hawk-dove game for types that vary only in giving up rate *β*. Coexistence requires *β*_*h*_ < *β*_0_ < *β*_*d*_, where *β*_0_ is shown as a yellow star. *β*_*h*_ and *β*_*d*_ strongly modify the ratio of resources set by each type at equilibrium,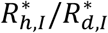. Coexistence occurs in the upper-left, where 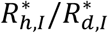 is the farthest from unity. Note the log scale for color. *τ* = 1, *a* = 0.1, *m* = 0.015, *e* = 0.035, *m*_*C*_ = 0, *r* = 0.26, *K* = 250.

*R*^∗^ theory predicts the exclusion of the consumer that sets a higher equilibrium resource abundance. However, coexistence predominantly occurs when the equilibrium resources abundances differ by a large factor (Fig. 2, top-left corner), while small differences are associated with competitive exclusion. Coexistence corresponds to a greater *R*^∗^ ratio 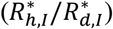 because dove invasion requires intra-hawk interference to increase the equilibrium resource density set by the hawk. The ratio of equilibrium resources abundances 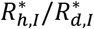 should not be confused with the exploitative *R*^∗^ ratio; 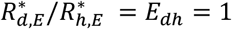 throughout Fig. 2).

We now consider types that also differ in exploitative ability (in contrast to Fig. 2 above, in which all parameters except *β* were identical). Instead of denoting types by their giving up rates as hawk *h* and dove *d*, we distinguish types by their relative exploitative ability. We denote type 2 as better at exploitation than type 1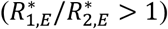; this implies nothing about which type is the hawk vs. dove with respect to *β*. The sometimes dramatic extent to which exploitative differences change the region of coexistence (Figs. 3-4), depends on the relative impact of mechanistic parameters on interference vs. exploitative differences.

**Fig. 3:**
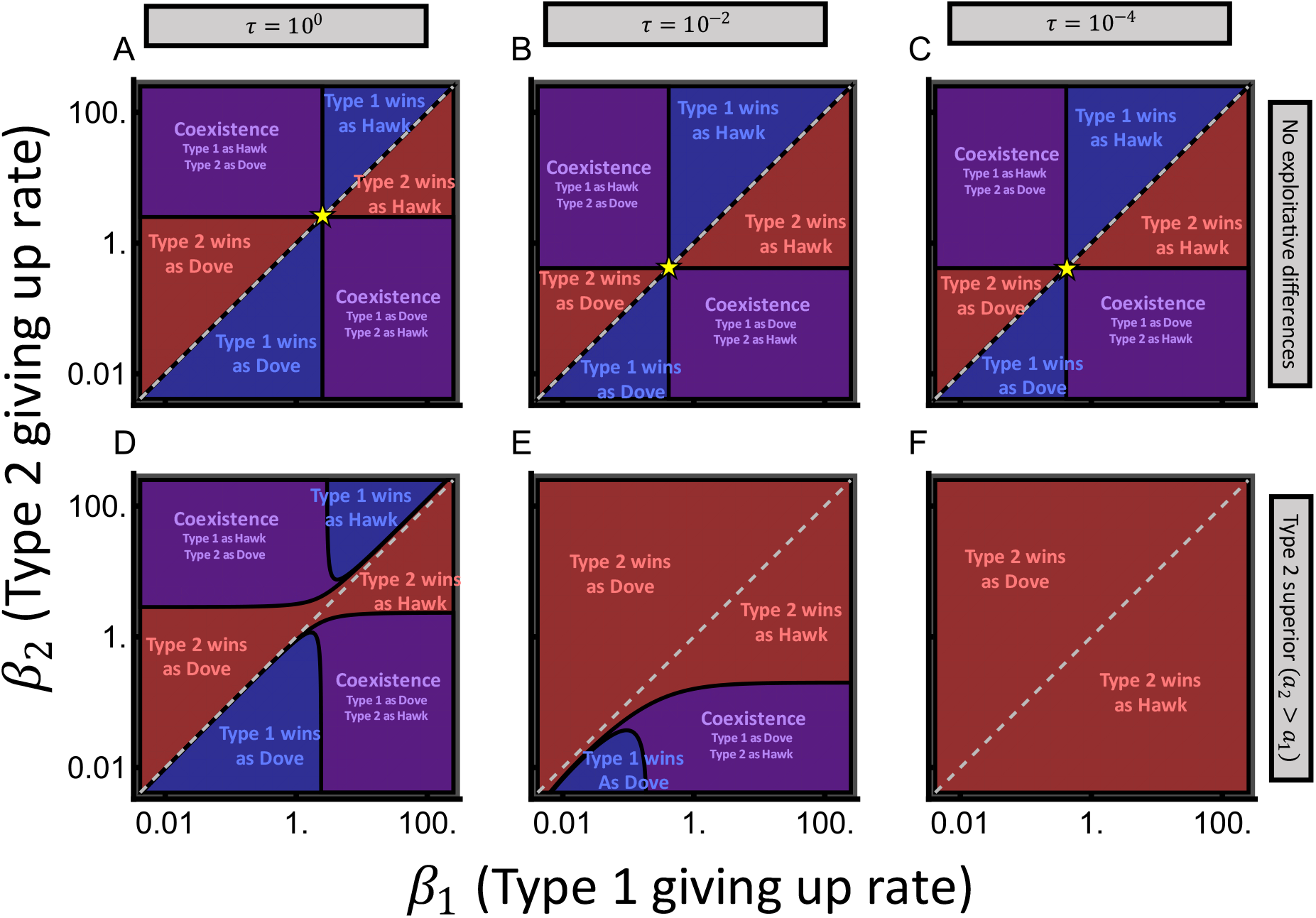
Short handling time erodes coexistence when types differ in exploitative ability. Consumers 1 and 2 differ in search rate (*a*, in bottom row only) and giving up rate (*β*, both rows). Below the dashed diagonal line, *β*_1_ > *β*_2_ makes type 1 the dove, while type 1 is the hawk above the line. Columns vary the handling time. Panels A-C depict identical exploitative abilities; gold stars are as defined in Fig. 2. In panels D-F, type 2 is better at exploitation (*a*_2_ = 0.12 > *a*_2_ = 0.10). Other parameters are: *m*_1_ = *m*_2_ = 0.015, *e*_1_ = *e*_2_ = 0.035, *m*_*C*1_ = *m*_*C*2_ = 0, *r* = 0.26, *K* = 250.

**Fig. 4:**
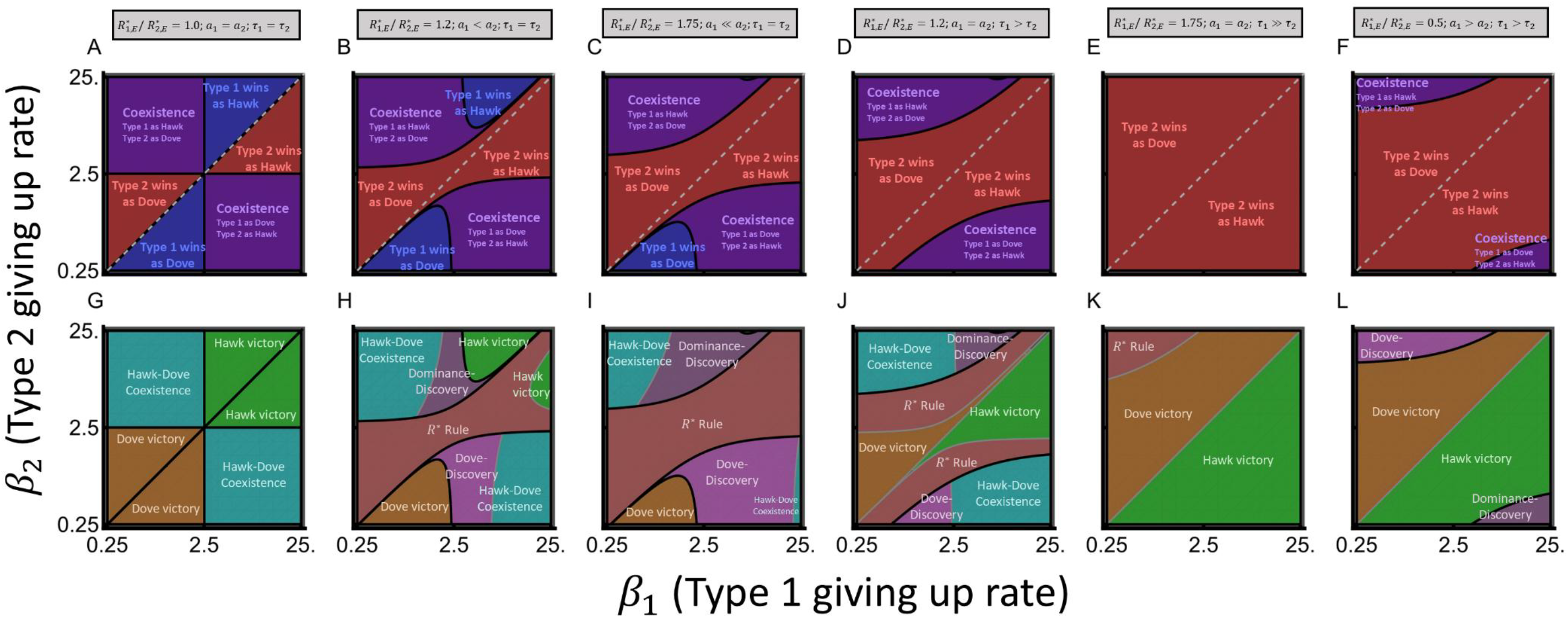
Differences in exploitative ability effected by search speed *a* permit more coexistence than those effected by handling time *τ*. Top row (A-F) depicts outcomes, bottom row 2 (G-H) the corresponding dominant mechanism from Table 1. Type 1 is the dove below the dashed diagonal line, and is the hawk above. The first column shows consumers that are identical in every exploitative parameter. The second and third columns show differences in search rate (*a*_2_ = 0.12 or *a*_2_ = 0.175 > *a*_1_ = 0.1). The fourth and fifth columns show differences in handling rate (*τ*_2_ = 1/1.35 or *τ*_2_ = 1/30 < *τ*_1_ = 1). Both vary in the rightmost column (0.35 = *a*_1_ > *a*_2_ = 0.1; 1/5.19 = *τ*_2_ < *τ*_1_ = 1). When not specified above, *a*_1_ = *a*_2_ = 0.1, *τ*_1_ = *τ*_2_ = 1, *m*_1_ = *m*_2_ = 0.015, *e*_1_ = *e*_2_ = 0.035, *m*_*C*1_ = *m*_*C*2_ = 0, *r* = 0.26, *K* = 250. Results are robust to alternative choices (Supplementary Figures I3-I10).

For example, rapid handling of resources (small *τ*) decreases the frequency of contests, making exploitative competition relatively more important. Correspondingly, the region of coexistence declines with decreasing *τ* (Fig. 3, left to right) given a constant difference in exploitative ability (Fig. 3 bottom row,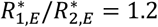 via *a*_2_ > *a*_1_), but not when exploitative ability is identical (Fig. 3 top row).

Differences in exploitative ability can be mediated by different mechanistic parameters (e.g. *a* vs. *τ*), which have different effects on interference competition, and hence on competitive outcomes (Fig. 4). Faster search *a* slightly increases opportunities to interfere with others, while shorter handling time *τ* dramatically decreases vulnerability to interference. In contrast, in traditional *R*^∗^ theory, only relative exploitative ability matters (i.e., *a* and *τ* only matter insomuch as they affect 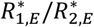).

When different search rates are responsible for a moderate difference in exploitative ability (e.g.,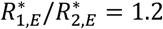 because *a*_2_ > *a*_1_; Fig 4B, 4H), outcomes are modestly different from when there are no exploitative differences (compare Figs. 4A and 4B), at least when *τ* is not very small (Fig. 3). However, the dominant mechanism behind the same outcome (coexistence or competitive exclusion) may be different (compare Figs. 4G and 4H). When giving up rates (*β*) are similar, the *R*^∗^ rule is the most influential mechanism (Fig. 4H, 4I). When coexistence occurs, different search rates shift the predominant mechanism from an inter-type vs. intra-type interference trade-off (hawk-dove coexistence) to an exploitation-interference trade-off (dominance-discovery or dove-discovery), noting mechanisms are not mutually exclusive. Coexistence remains possible even with larger differences in exploitative ability, e.g., when 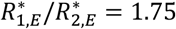 via *a*_2_ > *a*_1_in Fig 4C, 4I.

When differences in handling time *τ* drive the same difference in exploitative ability (e.g. 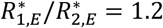 via *τ*_2_ < *τ*_1_; *a*_2_ = *a*_1_), coexistence becomes less likely than when driven by differences in search rate *a* (compare Figs. 4B with 4D and Figs. 4C with 4E). Coexistence largely evaporates when large exploitative differences are caused by *τ* (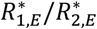 = 1.75 in Fig. 4E). Hawk or dove victories dominate the parameter space in which competitive exclusion occurs, rather than the *R*^∗^ rule (compare Fig. 4I and Fig. 4K).

When differences in base mortality rate *m* or conversion efficiency *e* drive differences in exploitative ability, coexistence patterns resemble those derived from differences in search rate *a* (Supplementary Figure I2). In these scenarios, the *R*^∗^ rule predominantly drives competitive exclusion (Supplementary Figure I2, E-H), reflecting the fact that *m* and *e* have minimal impact on interference.

The difference in exploitative ability 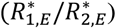 does not reliably explain outcomes. Fig. 4F shows an example where type 1 is superior at exploitation 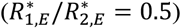 yet is excluded via interference by type 2, which searches more slowly (*a*_1_ > *a*_2_) but uses resources faster (*τ*_2_ < *τ*_1_). Type 1 is excluded in this scenario irrespective of whether it has a hawk or dove strategy (Fig. 4L). The disconnect between 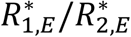 and outcomes reflects how mechanistic parameters that affect 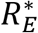 also impact interference.

## DISCUSSION

Exploitative resource competition is the foundation of much ecological theory, often in the form of *R*^∗^ theory (Tilman 1982), but its co-occurrence with interference competition has received less focus. Here, we analyze a mechanistic framework in which consumers’ handling of resources can be interrupted by interference from competitors, and time invested in contesting a resource incurs an opportunity cost of not searching for uncontested resources. Outcomes rarely follow *R*^∗^ theory, whether *R*^∗^ is interpreted as the equilibrium resource level 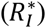 or as the hypothetical equilibrium resource level in the absence of interference (the minimum resource level required for persistence 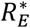). Coexistence can stem from a trade-off between inter-type interference and intra-type interference (hawk-dove coexistence), from a trade-off between exploitation and inter-type interference (dominance-discovery), or from a novel trade-off between exploitation and intra-type interference that we call dove-discovery. Which outcome occurs via which form(s) of competition is a complex outcome of parameter value combinations, with search rate and handling time simultaneously affecting both exploitation and interference.

### Interference competition and R^∗^ Theory

A key result of *R*^∗^ theory – the *R*^∗^ rule – breaks down with interference competition for two basic reasons. Firstly, interference allows the equilibrium resource density to rise, making the minimum tolerated resource density less important. Within *R*^∗^ theory, the minimum tolerated resource density 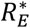 is equal to the equilibrium resource density 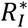. With interference, this is no longer true. Secondly, when invading consumers steal resources, their growth rate is higher than expected from the equilibrium density of free resources.

Our mechanistic terms for search rate and resource handling time affect not just exploitative ability, but also interference competition. Genotypes or species that achieve low 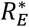 by handling resources more rapidly also gain substantial protection from interference. Genotypes or species that achieve the same low 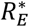 through faster search benefit more modestly from an increased ability to interfere with others. Previous models used handling time to describe relative non-linearity in consumers’ functional responses (Armstrong and McGehee 1980), and explored its effects on intrinsically driven consumer-resource oscillations (Johnson and Amarasekare 2015; Rosenzweig 1971). We emphasize for the first time the effect of handling time on interference rather than exploitation.

*R*^∗^ theory and the *R*^∗^ rule hold within some regions of parameter space (Fig. 4, Supplementary Figure I2); how common these are is an empirical question. While *R*^∗^ theory has experimental support (Wilson et al. 2007), this generally comes from well-mixed chemostats. Interference invokes co-localization of at least two individuals and a resource; well-mixed chemostats minimize the duration of co-localization in which a contest can occur. Outside the laboratory, spatial structure is ubiquitous and interference competition is widespread; perhaps most obviously in animals, often over territories (Drury et al. 2020; Schoener 1983), but also among plants via soil interactions and allelopathy (Inderjit et al. 2011; Meiners et al. 2012; Rice 2012), and among microbes through mechanisms such as stabbing and toxin-antitoxin systems (Böck et al. 2017; Boynton 2019; García-Bayona and Comstock 2018; Ghoul and Mitri 2016).

Measuring outcomes plus exploitative ability provides insufficient evidence in favor of the *R*^∗^ rule, because the same consumer can also be favored by interference competition. Ideally, experiments and models should be tailored to the mechanistic biology of the organisms in question. Comparing 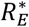(e.g. as implied by measured trait values *a, m, e*, and *τ*) to 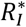 (as measured by the equilibrium resource abundance) could quantify the strength of negative intra-specific density-dependence driven by interference.

### Comparison with Past Coexistence Models

High consumer density in our model both depletes resources (exploitation), and increases interaction frequency (interference). In contrast, because every round of the classic hawk-dove game (Maynard Smith 1974) involves a contest, the payoff matrix does not depend on density. Our direct dependence on consumer density also differs from payoff matrices of interacting competitors that depend on environmental variables such as resource density (Tilman et al. 2020). Our underlying model is more similar to work incorporating density-dependence into game theory models, to position hawk-dove dynamics within more ecologically realistic scenarios (Auger et al. 2006; Galanthay et al. 2023; Kooi 2015; Křivan and Cressman 2022; Křivan et al. 2018). Our work formally connects this class of models and, by extension, the hawk-dove game, to *R*^∗^ theory.

We distinguish between two different exploitation-interference trade-offs. One resembles the dominance-discovery trade-off described by Adler et al. (2007), in which inter-type interference enables hawk invasion, and superior exploitation enables “dove” invasion (typically via faster resource discovery/search rate). In contrast, it is intra-hawk interference that enables dove invasion under hawk-dove coexistence. The dominance-discovery trade-off can alternatively be thought of as a “hawk-discovery” trade-off in our model.

Competition-colonization trade-off models (Hastings 1980; Horn and MacArthur 1972; Tilman 1994) are formally similar. Our model has two key differences. Firstly, as in Adler et al. (2007), we consider consumption of an exhaustible resource instead of durable territories. Secondly, “competition” in our model is interpreted as interference, rather than as local superiority in exploitation (e.g., a plant species with a lower *R*^∗^ for nitrogen that locally excludes its competitor on a patch of a metacommunity; Tilman 1994). Classically, coexistence requires the inferior exploiter (with higher local *R*^∗^ for nitrogen) to colonize patches more rapidly. This can be interpreted as having a lower “global” *R*^∗^, despite higher local *R*^∗^, where global *R*^∗^ represents the equilibrium proportion of unoccupied patches set by a species in monoculture. Coexistence in these classic models thus requires a trade-off between local and global exploitation competition with corresponding *R*^∗^s. These local and global *R*^∗^*s* should not be confused with 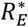 and 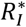 introduced in our model.

We discovered a second and perhaps counterintuitive exploitation-interference trade-off: “dove-discovery”. Here the hawk invades due to its superior exploitative abilities, in addition to its (by definition) superiority at inter-type interference. The dove can nevertheless invade because intra-hawk interference frees up resources. Hawk invasion can require an exploitative advantage because winning inter-type contests might not be enough to counter the high opportunity costs of those inter-type contests. This emphasizes how relatively doveish behavior can yield a competitive advantage.

Dove-discovery complicates empirical efforts to detect and interpret trade-offs in interference and exploitative abilities, e.g. in ants suspected to be subject to the discovery-dominance trade-off (Davidson 1998; Fellers 1987; Holway 1999). Under dominance-discovery, coexisting species/types who are better at obtaining resources are expected to be less likely to win contests, at least in the absence of additional trade-offs (e.g., vulnerability to parasitoids Adler et al. 2007; LeBrun 2005). However, under dove-discovery, species or genotypes that are more likely to win contests are also better at obtaining resources – the exact opposite pattern. Similarly, under hawk-dove coexistence, either the hawk or dove may be superior at exploitation. Empirical investigations looking for trends consistent with dominance-discovery and competition-colonization trade-offs (often yielding mixed support; for a recent review, see Ferzoco and McCauley 2023) may therefore overlook key interference-related coexistence mechanisms. Hawk-dove coexistence and the dove-discovery trade-off have not received similar levels of attention in this context, perhaps because previous theory had not outlined the conditions under which they occur. Hawk-dove and dove-discovery make up a large region of coexistence space (Fig. 4) and may turn out to be as or more important than dominance-discovery for maintaining coexistence in empirical communities (although the mechanisms are not mutually exclusive).

### Previous Models of Interference Competition

Most previous models of interference competition use Lotka-Volterra-like terms in phenomenological models to summarize the strengths of intra- and inter-specific interference (Case and Gilpin 1974; Hsu 1982; Schoener 1976; Schoener 1978; Vance 1984). Papers such as Vance (1984) suggest that coexistence requires intra-specific interference to exceed inter-specific interference. Our model shows this is not necessary: coexistence occurs even if both the high-interference and the low-interference genotypes/species are each equally prone to interfere with their own type as with the other type.

McPeek (2012) incorporated a phenomenological intra-specific density-dependent term into a consumer-resource model, which increases the resource level set by a resident, similarly producing 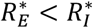. Adding this term allows multi-consumer coexistence on a single resource (also see Kuang et al., 2003). Our model provides a mechanistic basis by which this might occur.

Amarasekare (2002), analyzing a model similar to ours, assumed no intra-specific interference (implying 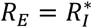), and concluded that (1) mutually costly forms of interference (e.g., aggression, territoriality) cannot maintain coexistence, and (2) interference-exploitation trade-offs can facilitate coexistence when interference is directly beneficial to the inferior exploiter. Amarasekare (2002) then interprets directly beneficial interference as intraguild predation or parasitism, to argue that the restricted nature of these scenarios limits the role of interference in promoting coexistence. An alternative interpretation is that aggression, territoriality, and interference over resources within dominance-discovery also qualify as beneficial because the aggressor can profit from acquiring stolen resources. Interference also promotes coexistence via hawk-dove dynamics and the dove-discovery trade-off, which are not examined in Amarasekare (2002). This suggests that the breadth of conditions under which costly interference competition maintains coexistence is much larger than previously recognized.

Previous models of interference (e.g., Amarasekare 2002; Hsu 1982; Vance 1984) found priority effects to be more prevalent than we did. Priority effects in these models were driven by stronger inter-specific interference than intra-specific interference. We did not depart from equal interference (*β*_*ii*_ = *β*_*ij*_), because this was not part of our focus on mechanistically linking *R*^∗^ theory, the hawk-dove game, and the dominance-discovery trade-off.

### Interference, Character Displacement, and the Evolution of Niche Differences

Our theory provides foundations with which one might understand how interference competition modifies character displacement and the evolution of niche differences (Brown and Wilson 1956; Grant and Grant 2006). Classic work on character displacement focuses on exploitative competition, an emphasis perhaps stemming from the remarkable associations of beak size/shape and seed diet observed in Darwin’s finches (Grant 1999; Grant and Grant 2006). However, several recent studies highlight so-called Agonistic Character Displacement (ACD), broadly defined as phenotypic evolution due to interference competition among sympatric species (Grether et al. 2013; Grether et al. 2009; Grether et al. 2017). ACD enjoys growing empirical support (Anderson and Grether 2010; McEachin et al. 2024; Moran and Fuller 2018). Our model provides a basis with which to contextualize the role of interference competition, especially ACD, in niche evolution.

The fact that handling time modifies both exploitation and interference has implications for niche evolution as depicted in classic ecological models that only consider exploitative competition. For example, MacArthur’s consumer–resource model (Chesson 1990; MacArthur 1970) describes a 2-consumer-multiple-resource exploitative competition model in which niche differences are defined in terms of a trait (e.g., beak shape) that determines its search rate for each resource (e.g., different seeds sizes/shapes). The evolution of niche differences is construed as the selective pressure driven by inter-specific overlap in resource exploitation (Pastore et al. 2021; Song et al. 2019). MacArthur’s consumer–resource model ignores handling time, which does not matter for exploitation-only models. However, shorter handling times decrease vulnerability to interference. Many specialized morphological features (e.g. bird beak morphology and its compatibility with seed shape/size) do more to decrease handling time than to make search more rapid. Selection for trait specialization might be driven in part by escaping interference.

### Caveats, Future Directions, and Conclusion

We considered only two consumers competing for a single limiting resource. It remains an open question precisely how much more diversity interference helps maintain in more complex scenarios of more consumers and more resources.

Our model considers a spontaneously generated, consumable resource, rather than a biotic resource that undergoes a replicative form of growth from a starting population. The same general principles likely apply for other resource types, with an interesting caveat. Because our model introduces resource handling, consumers have non-linear functional responses (Holling 1965). For a replicating resource, this allows the possibility of stable limit cycles. When different non-linear functional responses result in a trade-off between fitness in low-resource environments (gleaners) and high-resource environments (opportunists), resource oscillations may contribute to coexistence (Armstrong and McGehee 1980; Yamamichi and Letten 2022).

Our modeling of interference as a delay before a motile organism “gives up” is not an obvious fit to sessile plants, although it is, following precedent in classic game theory (Maynard Smith and Price 1973), a good fit for motile animals (Ferzoco and McCauley 2023; Rodríguez et al. 2007; Stanton et al. 2002). Specifically, temporal investment in interference incurs an opportunity cost to foraging, facilitating coexistence. Plants also invest in local interference strategies, e.g., root competition, overgrowth, undercutting, and allelopathy, likely incurring an opportunity cost to reproduction and resource acquisition, rather than to foraging. Such trade-offs are likely ubiquitous, and process-based models of interference tailored to plant biology may yield results similar to our study.

Our model considers a consumable resource. Bertram and Masel (2019) considered competition for a durable resource such as territory, via three traits within a variable-density lottery model: the rate *b* that adults give birth to and disperse juveniles, juveniles’ ability *c* to contest each other to take possession of an unoccupied territory and thus become adults, and adult survival 1/*d*. Our searching term *a* is somewhat analogous to *b*, our persistence in interference competition 1/*β* is somewhat analogous to *c*, and our mortality *m* is directly analogous to *d*.

These similarities perhaps reveal profound patterns. (Grime 1977; Grime 1988; Grime 2006) hypothesized that trade-offs shaped species into three types of specialization: “ruderals” tolerate low resource levels, “competitors” excel at interference competition, and “colonizers” rapidly disperse to ephemeral resources. Grime’s three-dimensional scheme is often contrasted with *R*^∗^ models of exploitative competition (the “Grime-Tilman debate”; (Aerts 1999; Craine 2005; Jabot and Pottier 2012). A major obstacle to synthesis has been the lack of explicit mathematical models of Grime’s scheme (Jabot and Pottier 2012; Tilman 2007). Our framework mathematically unites ideas of Grime with ideas of Tilman. Grime imagined low density environments in which intraspecific competition (both interference and exploitation) was weak and ruderals would prevail, whereas Tilman’s view was that organisms in apparently low density environments simply had larger territories, with resource levels still set by consumption down to the lowest tolerated level. Our work provides loose support for Grime’s view, by highlighting that resource levels need not be set by exploitation alone.

In the ant literature, Grime’s colonizers are called discoverers, Grime’s competitors are called dominators, and a slightly different take on Grime’s ruderals is known as the insinuator foraging strategy (Hölldobler and Wilson 1990; LeBrun 2005). Insinuators adopt non-aggressive strategies to capitalize on a small portion of resources. In our model, discoverers are characterized by a high search rate, and dominators by a low giving up rate. To incorporate insinuators, we could expand our model to include multiple resources of differing quality. Discoverers are more likely to discover high-value resources, dominators to steal them, while insinuators rely on low-quality resources that are not worth the handling time of dominators and are therefore uncontested. Insinuators are thus characterized by both non-aggressive behavior, and better yield from low value resources.

The aggressive behavior that we capture as lower *β* is likely to co-evolve with other critical parameters (e.g. *m, a, τ*). Concerted trait combinations seem likely, including traits not considered by our model. E.g., types may differ in intrinsic fighting abilities, as reflecting in the death rate of their opponent. Physiological constraints might impose trade-offs, e.g., between fighting, reducing vulnerability to interference *τ*, dispersing *a*, and persisting at low resource levels *m* and *e*. Incorporating these into adaptive dynamic models (Dieckmann et al. 2006; Leimar 2009) may help understand the evolution of aggression in motile organisms, and which trade-offs promote or inhibit species coexistence.

## CODE AVAILABILITY

All code is available in Supplementary Files and on GitHub: https://github.com/DanielSmithEcology/R_Star_Interference_Competition

## ACKNOWLEDGEMENTS

We thank Jason Bertram and Matt McCaskey for early conceptual work in this area, and Sasha Dall for helpful discussions on conceptual issues related to the hawk-dove game. We thank three anonymous reviewers and Gaku Takimoto for suggestions that greatly improved the manuscript. We thank the John Templeton Foundation (62220) for funding. DJBS also thanks the Peter O’Donnell Jr. Postdoc Fellowship for funding.

