## Supplementary Materials for "A Mechanistically Integrated Model of Exploitative and Interference Competition over a Single Resource Produces Widespread Coexistence"

#### Section A: Lack of Oscillations with $f(R) = r(K - R)$

In the main text, we claim that the 1-consumer-1-resouce system

$$\frac{dS_i}{dt} = \frac{1}{\tau_i}(1 + e_i)H_i + 2\beta_i C_{ii} - m_i S_i - a_i S_i H_i - a_i S_i R$$

$$\frac{dH_i}{dt} = a_i S_i R + (2\beta_i + 2m_i)C_{ii} - m_i H_i - H_i/\tau_i - a_i S_i H_i \qquad (1A)$$

$$\frac{dC_{ii}}{dt} = a_i S_i H_i - (2\beta_i + 2m_i)C_{ii}$$

$$\frac{dR}{dt} = f(R) - a_i S_i R + m_i H_i$$

does not exhibit oscillations at the non-trivial equilibrium when $f(R) = r(K - R)$. In principle, this can be determined by examining the eigenvalues of the Jacobian matrix of the $4 \times 4$ system described by 1A, evaluated at the non-trivial equilibrium:

$$S_i^* = \frac{r\big(a_i K(e_i - m_i\tau_i) - m_i(m_i\tau_i + 1)\big)}{a_i m_i \left(1 + \frac{Kra_i\tau_i}{\beta_i + m_i}\right)}$$

$$H_i^* = \frac{r\tau_i(a_i(e_i K - Km_i\tau_i) - m_i(1 + m_i\tau_i))}{a_i\left(\frac{m_i r\tau_i(1 + m_i\tau_i)}{\beta_i + m_i} + e_i - m_i\tau_i\right)} \qquad (2A)$$

$$C_{ii}^* = \frac{r^2\tau_i(\beta_i + m_i)\big(a_i K(m_i\tau_i - e_i) + m_i(m_i\tau_i + 1)\big)^2}{2a_i m_i\big(e_i(\beta_i + m_i) + m_i\tau_i(-\beta_i + m_i(r\tau_i - 1) + r)\big)(Kra_i\tau_i + \beta_i + m_i)}$$

$$R_{i,I}^* = \frac{m_i(m_i\tau_i + 1)\left(K\frac{a_i r\tau_i}{\beta_i + m_i} + 1\right)}{a_i\left(e_i - m_i\tau_i + \frac{m_i r\tau_i(m_i\tau_i + 1)}{\beta_i + m_i}\right)}$$

If all the eigenvalues of the Jacobian matrix have negative real parts, the system will converge to a stable limit point (i.e., not exhibit persistent oscillations). Unfortunately, we find that the complexity of the model precludes analytically tractable methods such as the Routh–Hurwitz criterion.

Instead, we use a numerical approach to examine the stability/oscillatory nature of the system. We numerically investigate stability using a Latin Hypercube Sampling (LHS) method, sampling a wide variety of parameter combinations. In all feasible parameter combinations (parameters that permit positive equilibria), we find the Jacobian matrix always exhibits four negative eigenvalues (Fig. A1). Therefore, by the stable manifold theorem, the system converges to a stable (non-oscillating) point. See "Supplementary_Material_A_Eigenvalues.R"

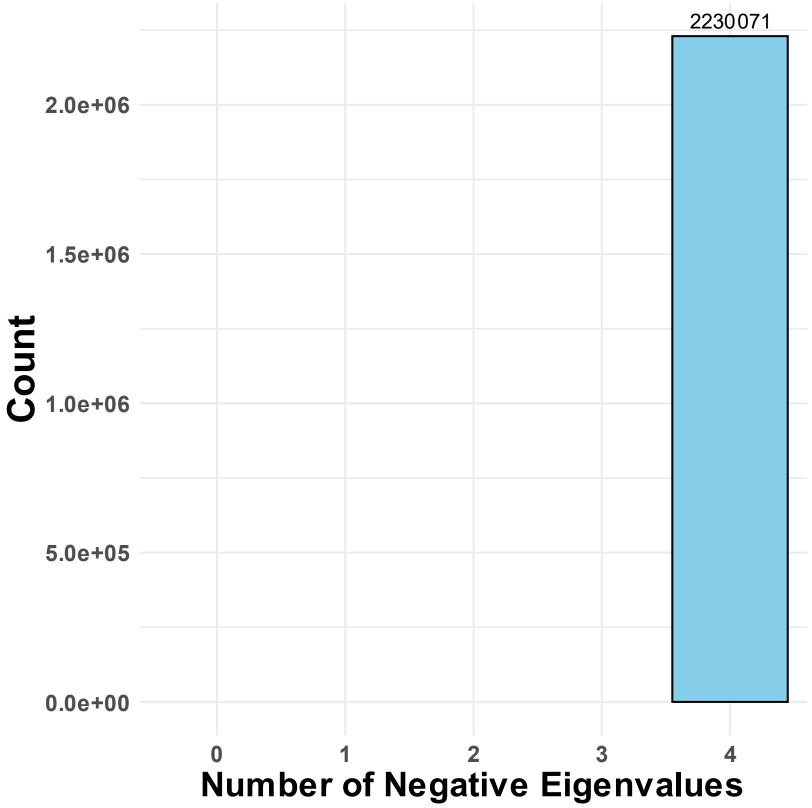


**Fig. A1:** All the eigenvalues from the sampled feasible and non-trivial equilibria of the system in equations in $2A$ are negative. Hence, persistent oscillations do not occur. Parameter values were sampled using Latin Hypercube Sampling wherein: $a_i \sim \{.01,100\}$, $\tau_i \sim \{.001,100\}$, $\beta_i \sim \{.01,1000\}$, $e_i \sim \{.01,100\}$, $m_i \sim \{.0001,10\}$, $r \sim \{.01,1000\}$ and $K \sim \{5,5000\}$. Sampling was performed on a log scale. From an initial $5 \times 10^6$ samples of parameter space, the plot depicts the ~$2.2 \times 10^6$ feasible samples.

### Section B: Logistic Resource Growth Gives Similar Results

In the main text, we claim that a logistically growing resource yields qualitatively similar results (in terms of coexistence) to the spontaneously regenerating consumable resource examined in the main text.

Here, we show an example of the similarity. Using the same parameter values, we present ODE simulations of the system in Equation (1) in the main text (also Equation 1A above) under two scenarios. (1) Using logistic resource growth according to $f(R) = rR(1 - R/K)$, and (2) using $f(R) = r(K - R)$ as in the main text. Fig. B1 shows that patterns of competitive exclusion and coexistence have quantitative differences, but are qualitatively similar. Hence, the functional form of resource growth is not likely to qualitatively alter our conclusions. Simulations were run using the DifferentialEquations package in Julia. See "Supplementary_Material_B_ODE_Simulations.jl" for more simulation details.

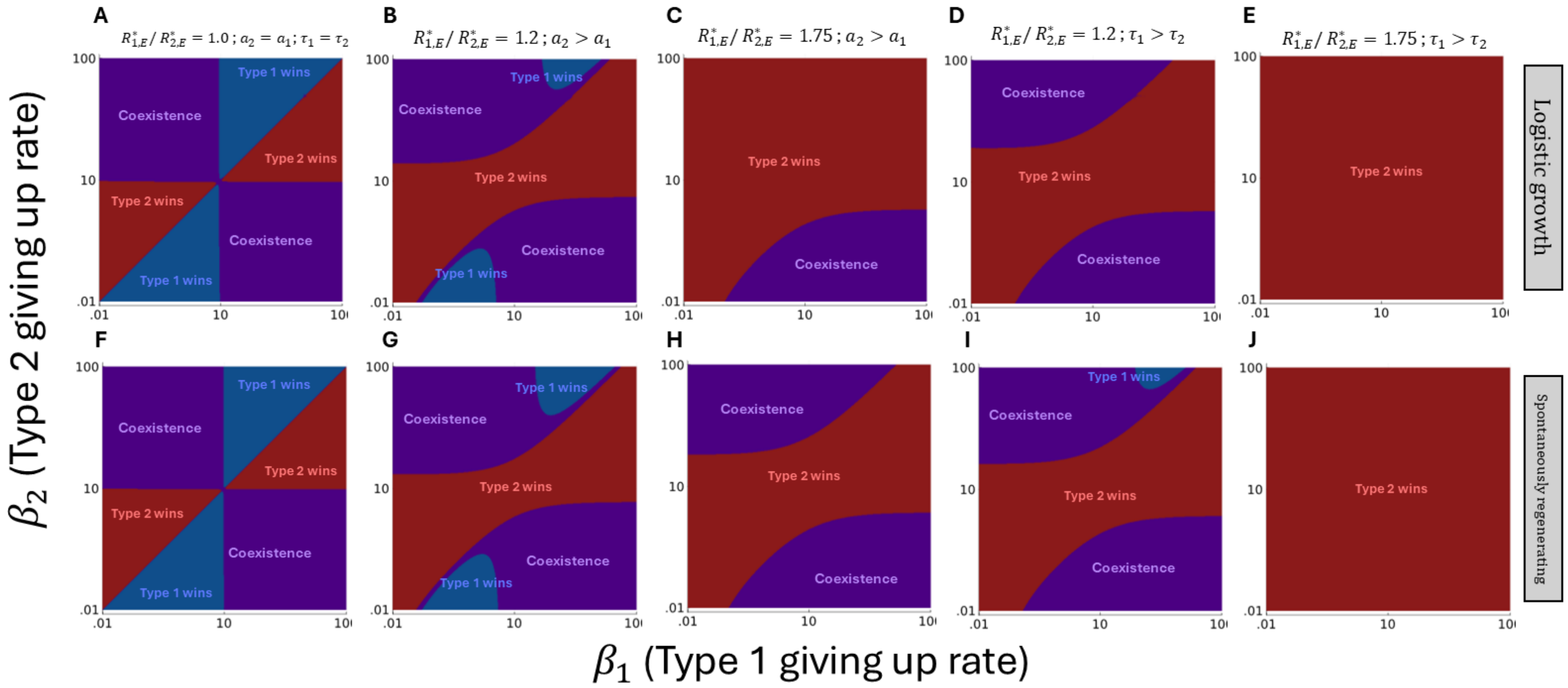


**Fig. B1:** Logistic resource growth and spontaneously regenerating resources yield similar result. Row 1 (A-E) depicts logistic resource growth; F-J show a spontaneously regenerating resource. The first column shows consumers that are identical in every exploitative parameter. The second and third columns show differences in search rate ($a_2 = 0.06$ or $a_2 = 0.0875 > a_1 = 0.05$). The fourth and fifth columns show differences in resource use rate ($\tau_2 = 1/1.35$ or $\tau_2 = 1/30 < \tau_1 = 1$). When not specified above, $a_i = 0.05$, $\tau_i = 1$, $m = 0.015$, $e = 0.035$, $m_C = 0$, $r = 1.0$, $K = 25$.

### Section C: Coexistence Follows Mutual Invasibility in Simulations

In the main text, we claim that numerical simulations of the system of Ordinary Differential Equations in equation (1) from the main text yield coexistence results identical to the invasion analysis. Here, we show the convergence of ODE simulations and the invasion criteria.

Using parameters identical to Fig. 4 in the main text, we ran numerical simulations of the ODE described by the system in equation (1). Results are shown in Fig. C1. Note that the results (coexistence, type $i$ wins, type $j$ wins) are identical to those in Fig. 4A-5F. This demonstrates that the invasion criteria as defined by Equation (11) accurately delineates competitive outcomes. Simulations were run using the DifferentialEquations package in Julia. See "Supplementary_Material_C_Simulations.jl" for simulation details.

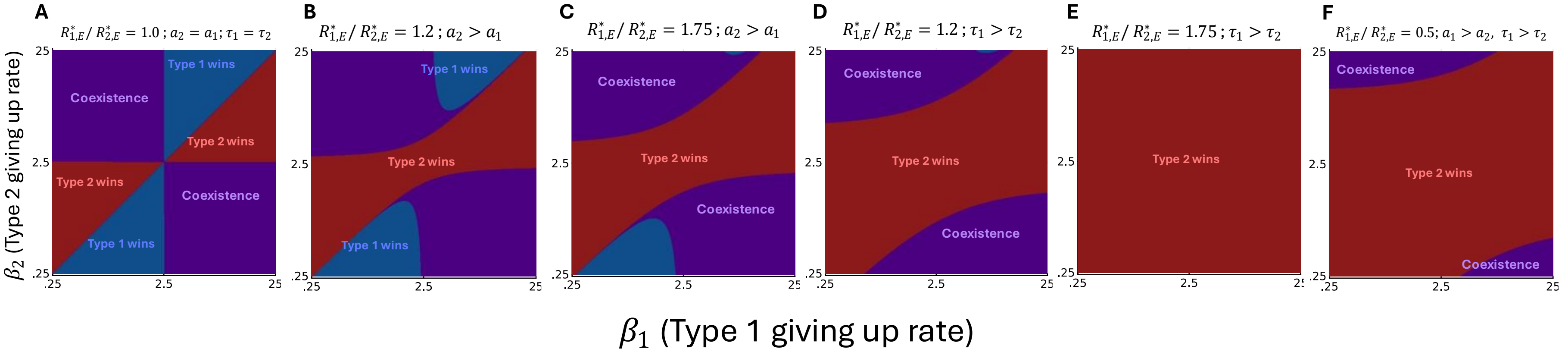


**Fig. C1** ODE simulations yield coexistence results identical to those predicted by the invasion analysis. Compare this figure to Fig. 4A-4F in the main text. As in Fig. 4, **A** shows identical exploitative ability. **B** and **C** show differences in search rate ($a_2 = 0.12$ or $a_2 = 0.175 > a_1 = 0.1$). The **D** and **E** show differences in resource use rate ($\tau_2 = 1/1.35$ or $\tau_2 = 1/30 < \tau_1 = 1$), and **F** shows differences in both ($0.35 = a_1 > a_2 = 0.1;\ 1/5.19 = \tau_2 < \tau_1 = 1$). When not specified above, $a_i = 0.1, \tau_i = 1, m = 0.015, e = 0.035, m_C = 0, r = 0.26, K = 250$. Compare to Fig. 4A-4F from the main text.

### Section D: Invasion Criteria, No Interference Competition

#### Introduction:

We show a step-by-step derivation of the invasion criteria for the "No Interference Competition" case -- when neither consumer partakes in interference competition. Here, we provide all details in the derivation of equations 2-3 in the main text.

As noted in the main text, we employ invasion analysis to derive the invasion criteria. To do so, we derive the Jacobian Matrix of the system when a single consumer (the resident) is at equilibrium and the other consumer (the invader) is at low abundance (i.e., $\approx 0$). If any eigenvalues of the Jacobian Matrix are positive, it implies the system is unstable and the invader tends to increase in abundance when rare. If both consumers increase in abundance when rare, it implies mutual invasibility (i.e., coexistence).

We examine the system of Ordinary Differential Equations given by:

$$\begin{aligned} \frac{dS_i}{dt} &= \frac{1}{\tau_i}(1+e)H_i - m_i S_i - a_i S_i R \\ \frac{dH_i}{dt} &= a_i S_i R - m_i H_i - \frac{H_i}{\tau_i} \qquad (1D) \\ \frac{dR}{dt} &= f(R) - R\sum_{j=1}^{2} a_j S_j + \sum_{j=1}^{2} m_j H_j \end{aligned}$$

which describe the dynamics of Searchers, Handlers, and the Resource; $j = 1, 2$ and $f(R) = r(K - R)$.

### D.1 Consumer-Resource Equilibrium Calculation

We solve for the equilibrium of the ODE system describing the 1-Consumer-1-Resource system (with only consumer type $i$). There is a single non-trivial equilibrium:

```
(*Solve for the equilibria*)
ConResEquil = Simplify[Solve[ 1/τi (1 + ei) Hi − mi*Si − ai*R*Si == 0 &&
        ai*R*Si − mi*Hi − Hi/τi == 0 &&
        r*(K − R) − ai*R*Si + mi*Hi == 0,
          {Si, Hi, R}]];
(*Save variables of the equilibria (searching consumers,handling consumers,resource level),
which are used for invasion analysis.*)
Sstari = Si /. Part[Part[ConResEquil, 2], 1];
Hstari = Hi /. Part[Part[ConResEquil, 2], 2];
Rstari = R /. Part[Part[ConResEquil, 2], 3];
(*Print the non−trivial equilibria*)
Part[ConResEquil, 2] // TraditionalForm
```

$$\left\{S_i \to r\left(-\frac{m_i\,\tau_i+1}{a_i}+\frac{K\,e_i}{m_i}-K\,\tau_i\right),\ H_i \to -\frac{r\,\tau_i\,(a_i\,(K\,m_i\,\tau_i-K\,e_i)+m_i\,(m_i\,\tau_i+1))}{a_i\,(e_i-m_i\,\tau_i)},\ R \to \frac{m_i\,(m_i\,\tau_i+1)}{a_i\,(e_i-m_i\,\tau_i)}\right\}$$

That is, the equilibria are:

$$S_i^* = r\left(-\frac{m_i\,\tau_i+1}{a_i}+\frac{K\,e_i}{m_i}-K\,\tau_i\right)$$
$$H_i^* = -\frac{r\,\tau_i\,(a_i\,(K\,m_i\,\tau_i-K\,e_i)+m_i\,(m_i\,\tau_i+1))}{a_i(e_i-m_i\,\tau_i)}$$
$$R_i^* = \frac{m_i(m_i\,\tau_i+1)}{a_i(e_i-m_i\,\tau_i)}$$

### D.2 Calculation of the Jacobian Matrix

We derive the invasion condition of type $j$ ($j \neq i$) at low density, invading a community of type $i$. To do so, we derive the eigenvalues of the Jacobian matrix taken at the equilibria of consumer $i$ and with type $j$ at low abundance. The Jacobian is given by:

$$J_j = \begin{pmatrix} \partial S_j \big/ \partial S_j & \partial S_j \big/ \partial H_j \\ \partial H_j \big/ \partial S_j & \partial H_j \big/ \partial H_j \end{pmatrix} \qquad (2\,D)$$

which is taken at the points: $S_j = 0,\ H_j = 0,\ R_j = 0$ and $S_i = S_i^*,\ H_i = H_i^*,\ R_i = R_i^*$:

```
(* This code will take the Jacobian Matrix of J_j,
and substitute the values S_j=0,H_j=0 and S_i=S_i^*,H_i=H_i^*,R=R_i^* *)
ConsumerjMatrix = {(*Put the dynamics of type j into matrix form*)
        -a_j * R * S_j + (H_j / τ_j) + (e_j * H_j / τ_j) - m_j * S_j,
         a_j * R * S_j - (H_j / τ_j) - m_j * H_j};

(*Calculate the Jacobian Matrix (J_j in the above equation)*)
JacMat = ResourceFunction["JacobianMatrix"][ConsumerjMatrix, {S_j, H_j}];

(*Take the limit when type j is rare and S_i=S_i^*,H_i=H_i^*,R_i=R_i^* *)
JacMat2 = Limit[JacMat, {S_j → 0, H_j → 0, S_i → Sstar_i, H_i → Hstar_i, R → Rstar_i}];
(*note that we use the notation e.g. Rstar_i to represent R_i^* for syntax reasons*)

(*Show the Jacobian Matrix*)
JacMat2 // TraditionalForm
```

$$\begin{pmatrix} -a_j \,\text{Rstar}_i - m_j & \frac{e_j}{\tau_j} + \frac{1}{\tau_j} \\ a_j \,\text{Rstar}_i & -m_j - \frac{1}{\tau_j} \end{pmatrix}$$

Thus,

$$J_j = \begin{pmatrix} -a_j R_i^* - m_j & e_j / \tau_j + 1 / \tau_j \\ a_j R_i^* - m_j & -m_j - 1 / \tau_j \end{pmatrix} \qquad (3D)$$

### D.3 Calculation of Eigenvalues

The eigenvalues of matrix $J$ ($\lambda_k$, $k = 1, 2$) are:

```
(*Calculate both eigenvalues of 2 x 2 matrix*)
λ1 = Part[Eigenvalues[JacMat2], 1]; (*eigenvalue 1*)
λ2 = Part[Eigenvalues[JacMat2], 2]; (*eigenvalue 2*)
(*Show eigenvalues*)
{λ1 // TraditionalForm, λ2 // TraditionalForm}
```

$$\left\{\frac{-\sqrt{4\,a_j\,e_j\,\text{Rstar}_i\,\tau_j + a_j^2\,\text{Rstar}_i^2\,\tau_j^2 + 2\,a_j\,\text{Rstar}_i\,\tau_j + 1} - a_j\,\text{Rstar}_i\,\tau_j - 2\,m_j\,\tau_j - 1}{2\,\tau_j}, \frac{\sqrt{4\,a_j\,e_j\,\text{Rstar}_i\,\tau_j + a_j^2\,\text{Rstar}_i^2\,\tau_j^2 + 2\,a_j\,\text{Rstar}_i\,\tau_j + 1} - a_j\,\text{Rstar}_i\,\tau_j - 2\,m_j\,\tau_j - 1}{2\,\tau_j}\right\}$$

That is, the eigenvalues are:

$$\lambda_k = \frac{\pm\sqrt{4\,a_j\,e_j\,R_i^*\,\tau_j + a_j^2\,(R_i^*)^2\,\tau_j^2 + 2\,a_j\,R_i^*\,\tau_j + 1} - a_j\,R_i^*\,\tau_j - 2\,m_j\,\tau_j - 1}{2\,\tau_j} \qquad (4D)$$

where

$$\text{Rstar}_i = R_i^* = \frac{m_i(m_i\,\tau_i + 1)}{a_i(e_i - m_i\,\tau_i)}$$

$i = 1, 2$. Recall that consumer $j$ will invade if there is at least one positive eigenvalue. $\lambda_k$ is always negative when the square root term is subtracted. Thus, we focus on when the square root term is added, $\lambda_2$.

### D.4 Rearrangement of Eigenvalues into Invasion Criteria

Here, we solve for when $\lambda_2 > 0$ in terms of $R_i^*$ (i.e., $\text{Rstar}_i$):

```
(* Find when λ2>0 in terms of Ri* (Rstari) *)
InvCrit = Reduce[λ2 > 0 && aj > 0 && ej > 0 && τj > 0 && Rstari > 0 && mj > 0 && ej > mj * τj, {Rstari}, Reals];
(*Solve the equation in terms of Rstari; assume that all the parameters are positive*)
(*Print solution*)
Simplify[Part[InvCrit, 5]] // TraditionalForm
```

$$\text{Rstar}_i > \frac{m_j\,(m_j\,\tau_j + 1)}{a_j\,(e_j - m_j\,\tau_j)}$$

This expression (substituting in $R_i^*$)

$$\frac{m_j(m_j\,\tau_j + 1)}{a_j(e_j - m_j\,\tau_j)} < \frac{m_i(m_i\,\tau_i + 1)}{a_i(e_i - m_i\,\tau_i)}$$

or

$$R_j^* < R_i^*$$

which is the invasion condition noted in the main text. Coexistence then requires

$$1 < \frac{R_j^*}{R_i^*} < 1$$

which cannot be fulfilled. This leads to competitive exclusion of the weaker exploitative competitor (the R* Rule).

### Section E: Derivation of $R^*_{i,I}$

In the main text, we show two different values of the equilibrium resource abundance when consumer $i$ interferes $R^*_{i,I}$ (Equation (4) and the expression below it). Here, we show where they originate from.

We consider the system of ODEs describing the consumer-resource dynamics of type $i$. The system is given by:

$$\frac{dS_i}{dt} = \frac{1}{\tau_i}(1+e_i)h_i + 2\beta_i C_{ii} - m_i S_i - a_i S_i i_i - a_i S_i R$$
$$\frac{dH_i}{dt} = a_i S_i R + (2\beta_i + 2m_i) C_{ii} - m_i i_i - H_i/\tau_i - a_i S_i H_i \qquad (1E)$$
$$\frac{dC_{ii}}{dt} = a_i S_i i_i - (2\beta_i + 2m_i) C_{ii}$$
$$\frac{dR}{dt} = f(R) - a_i S_i R + m_i H_i$$

where $f(R) = r(K - R)$.

$R^*_{i,I}$ is derived by solving the non-trivial equilibrium for $1E$:

```
(*Solve for the equilibria*)
ConResEquilHawk = Simplify[Solve[-a_i*R*S_i + (H_i/τ_i) + (e_i*H_i/τ_i) - m_i*S_i + 2 C_ii*β_i - a_i*H_i*S_i == 0 &&

        a_i*R*S_i - (H_i/τ_i) - m_i*H_i + 2 C_ii*β_i + 2 m_i*C_ii + 2*m_Ci*C_ii - a_i*H_i*S_i == 0 &&
                                a_i*H_i*S_i - 2*β_i*C_ii - 2*m_i*C_ii - 2*m_Ci*C_ii == 0 &&
                                r*(K - R) - a_i*R*S_i + m_i*H_i == 0, {S_i, H_i, C_ii, R}]];

(*Print the non-trivial equilibria*)
Part[ConResEquilHawk, 2] // TraditionalForm
```

$$\Big\{S_i \to -\frac{r(m_{Ci}+\beta_i+m_i)(a_i(K m_i \tau_i - K e_i) + m_i(m_i\tau_i+1))}{a_i(m_{Ci}(K r a_i \tau_i + m_i) + m_i(K r a_i \tau_i + \beta_i + m_i))},$$
$$H_i \to -\frac{r\tau_i(m_{Ci}+\beta_i+m_i)(a_i(K m_i \tau_i - K e_i) + m_i(m_i\tau_i+1))}{a_i(e_i(m_{Ci}+\beta_i+m_i) + \tau_i(m_{Ci}(m_i(r\tau_i-1)+r) + m_i(-\beta_i + m_i(r\tau_i - 1) + r)))},$$
$$C_{ii} \to \left(r^2\tau_i(m_{Ci}+\beta_i+m_i)(a_i(K m_i\tau_i - K e_i) + m_i(m_i\tau_i+1))^2\right)\Big/ (2a_i(e_i(m_{Ci}+\beta_i+m_i) + \tau_i(m_{Ci}(m_i(r\tau_i-1)+r) + m_i(-\beta_i+m_i(r\tau_i-1)+r))) (m_{Ci}(K r a_i\tau_i + m_i) + m_i(K r a_i \tau_i + \beta_i + m_i))),$$
$$R \to \frac{(m_i\tau_i+1)(m_{Ci}(K r a_i \tau_i + m_i) + m_i(K r a_i\tau_i + \beta_i + m_i))}{a_i(e_i(m_{Ci}+\beta_i+m_i) + \tau_i(m_{Ci}(m_i(r\tau_i-1)+r) + m_i(-\beta_i + m_i(r\tau_i-1)+r)))}\Big\}$$

That is, the equilibria are:

$$S_i^* = -\frac{r(m_{Ci}+\beta_i+m_i)(a_i(K m_i \tau_i - K e_i) + m_i(m_i\tau_i+1))}{a_i(m_{Ci}(K r a_i \tau_i + m_i) + m_i(K r a_i \tau_i + \beta_i + m_i))}$$

$$H_i^* = -\frac{r\tau_i(m_{Ci}+\beta_i+m_i)(a_i(K m_i \tau_i - K e_i) + m_i(m_i\tau_i+1))}{a_i(e_i(m_{Ci}+\beta_i+m_i) + \tau_i(m_{Ci}(m_i(r\tau_i-1)+r) + m_i(-\beta_i + m_i(r\tau_i - 1) + r)))}$$

$$C_{ii}^* = \frac{r^2\tau_i(m_{Ci}+\beta_i+m_i)(a_i(K m_i\tau_i - K e_i) + m_i(m_i\tau_i+1))^2}{2a_i(e_i(m_{Ci}+\beta_i+m_i) + \tau_i(m_{Ci}(m_i(r\tau_i-1)+r) + m_i(-\beta_i+m_i(r\tau_i-1)+r)))(m_{Ci}(K r a_i\tau_i + m_i) + m_i(K r a_i \tau_i + \beta_i + m_i))}$$

$$R_{i,I}^* = \frac{(m_i\tau_i+1)(m_{Ci}(K r a_i \tau_i + m_i) + m_i(K r a_i\tau_i + \beta_i + m_i))}{a_i(e_i(m_{Ci}+\beta_i+m_i) + \tau_i(m_{Ci}(m_i(r\tau_i-1)+r) + m_i(-\beta_i + m_i(r\tau_i-1)+r)))}$$

$R^*_{i,I}$ can be rearranged and the term $R^*_{i,E} = \dfrac{m_i(m_i\,\tau_i + 1)}{a_i(e_i - m_i\,\tau_i)}$ can be substituted in several places. This yields:

$$R^*_{i,I} = R^*_{i,E}\,\frac{1 + K\,\frac{r\,a_i\,\tau_i}{\beta_i + m_i + m_{C\,i}}\left(1 + \frac{m_{C\,i}}{m_i}\right)}{1 + R^*_{i,E}\,\frac{r\,a_i\,\tau_i}{\beta_i + m_i + m_{C\,i}}\left(1 + \frac{m_{C\,i}}{m_i}\right)}$$

which is the same as Equation (4) from the main text.

Secondly, in the main text we consider the case in which contests that are very fast (large $\beta_i$) but very deadly (large $m_{C\,i}$, recalling $m_{C\,i}$ is the increase in mortality due to contests). This can be achieved by making the substitution of parameters $\beta_i \to x\,\beta_i$, $m_{C\,i} \to x\,m_{C\,i}$ and taking the limit of $R^*_{i,I}$ as $x \to \infty$. We do so below:

```
(*Take the limit; the below code first substitutes βi→xβi and mCi→xmCi and then takes the limit as x→∞*)
RStarLimit =
 Simplify[Limit[Limit[(mi τi + 1) (mCi (K r ai τi + mi) + mi (K r ai τi + βi + mi)) /
      (ai (ei (mCi + βi + mi) + τi (mCi (mi (r τi - 1) + r) + mi (-βi + mi (r τi - 1) + r)))),
    {βi → x βi, mCi → x mCi}], x → Infinity]]
```

$$\frac{(1 + m_i\,\tau_i)\,(m_i\,\beta_i + m_{Ci}\,(m_i + K\,r\,a_i\,\tau_i))}{a_i\,(e_i\,(m_{Ci} + \beta_i) + \tau_i\,(-m_i\,\beta_i + m_{Ci}\,(r + m_i\,(-1 + r\,\tau_i))))}$$

Which, with some rearranging, gives

$$R^*_{i,I} = \;\to R^*_{i,E}\,\frac{1 + K\,c_i\,\frac{r\,a_i\,\tau_i}{m_i}}{1 + R^*_{i,E}\,c_i\,\frac{r\,a_i\,\tau_i}{m_i}}$$

where $c_i = \dfrac{m_{C\,i}}{m_{C\,i} + \beta_i}$. This is the same expression from the main text. Note that $c_i$ can be interpreted as the probability of mortality due to a contest. To see this, recall that constant rates in Ordinary Differential Equations (e.g., $d\,N / d\,t = -k * N$) are the deterministic mass action limit of exponentially distributed times to events. If two random variables, A and B, are exponentially distributed with rates a and b, P[A happens before B] = a/(a+b). In our context, B represents the probability a contest ends due to mortality and A represents the probability a contest ends due to giving up. Substituting the appropriate rates for A and B yields $m_{C\,i} / (m_{C\,i} + \beta_i)$.

### Section F: Invasion Criteria, Only One Type Interferes

#### Introduction:

Here, we show the derivation of the invasion criteria when one consumer (henceforth the "dove") never partakes in interference competition ($\beta_d = \infty$) and the other consumer (henceforth the "hawk") does interfere ($\beta_h < \infty$). We provide details for the derivation of equations 5-9 in the main text.

As noted in the main text, we employ invasion analysis. To do so, we derive the Jacobian Matrix of the system when a single consumer (the resident) is at equilibrium and the other consumer (the invader) is at low abundance (i.e., approximately 0). If any eigenvalues of the Jacobian Matrix are positive, it implies the system is unstable and the invader tends to increase in abundance when rare. If both consumers increase in abundance when rare, it implies mutual invisibility (i.e., coexistence).

### F.1 Consumer-Resource Equilibrium Calculation

The full dynamics of the system (from equation (1) from the main text with $\beta_d = \infty$) are:

$$\frac{dS_h}{dt} = \frac{1}{\tau_h}(1+e_h)H_h + 2\beta_h C_{hh} - m_h S_H - a_h S_h H_h - a_h S_h R$$
$$\frac{dS_d}{dt} = \frac{1}{\tau_d}(1+e_d)H_d - m_d S_d - a_d S_d R$$
$$\frac{dH_d}{dt} = a_d S_d R - m_d H_d - \frac{H_d}{\tau_d} - a_h S_h H_d \qquad (1F)$$
$$\frac{dH_h}{dt} = a_h S_h R + a_h S_h H_d + (2\beta_h + 2m_h) C_{hh} - m_h H_h - H_h/\tau_h - a_h S_h H_h$$
$$\frac{dC_{hh}}{dt} = a_h S_h H_h - (2\beta_h + 2m_h) C_{hh}$$
$$\frac{dR}{dt} = f(R) - a_h S_h R - a_d S_d R + m_h H_h + m_d H_d$$

where $f(R) = r(K - R)$. We analyze this system below.

First, we derive the equilibrium of the Ordinary Differential Equation (ODE) Consumer-Resource system in which a single consumer (hawk or dove) persists with the resource.

#### F.1.1 Analysis for Hawk

We consider the system of ODEs describing the consumer-resource dynamics for the hawk alone (without the dove). The system is given by:

$$\frac{dS_h}{dt} = \frac{1}{\tau_h}(1+e_h)\,h_h + 2\beta_h C_{hh} - m_h S_h - a_h S_h h_h - a_h S_h R$$
$$\frac{dH_h}{dt} = a_h S_h R + (2\beta_h + 2m_h) C_{hh} - m_h h_h - H_h/\tau_h - a_h S_h H_h \qquad (2F)$$
$$\frac{dC_{hh}}{dt} = a_h S_h h_h - (2\beta_h + 2m_h) C_{hh}$$
$$\frac{dR}{dt} = f(R) - a_h S_h R + m_h H_h$$

Note that in contrast to the exploitative competition only case (see Supplementary Material Section D), we include an extra state variable of hawks in conflict $C_{hh}$. At the non-trivial equilibrium

```
(*Solve for the equilibria*)
ConResEquilHawk =
    Simplify[Solve[-ah*R*Sh + (Hh/τh) + (eh*Hh/τh) - mh*Sh + 2 Chh*βh - ah*Hh*Sh == 0 &&
                   ah*R*Sh - (Hh/τh) - mh*Hh + 2 Chh*βh + 2 mh*Chh - ah*Hh*Sh == 0 &&
                   ah*Hh*Sh - 2*βh*Chh - 2*mh*Chh == 0 &&
                   r*(K - R) - ah*R*Sh + mh*Hh == 0, {Sh, Hh, Chh, R}]];

(*Print the non-trivial equilibria*)
Part[ConResEquilHawk, 2] // TraditionalForm
```

$$\left\{S_h \to -\frac{r(\beta_h+m_h)(a_h(K m_h \tau_h - K e_h) + m_h(m_h\tau_h+1))}{a_h m_h (K r a_h \tau_h + \beta_h + m_h)},\; H_h \to -\frac{r\tau_h(\beta_h+m_h)(a_h(K m_h\tau_h - K e_h) + m_h(m_h\tau_h+1))}{a_h(e_h(\beta_h+m_h) + m_h\tau_h(-\beta_h + m_h(r\tau_h-1)+r))},\right.$$
$$C_{hh} \to \frac{r^2\tau_h(\beta_h+m_h)(a_h(K m_h\tau_h - K e_h) + m_h(m_h\tau_h+1))^2}{2a_h m_h(e_h(\beta_h+m_h) + m_h\tau_h(-\beta_h + m_h(r\tau_h-1)+r))(K r a_h\tau_h + \beta_h + m_h)},$$
$$\left. R \to \frac{m_h(m_h\tau_h+1)(K r a_h\tau_h + \beta_h + m_h)}{a_h(e_h(\beta_h+m_h) + m_h\tau_h(-\beta_h + m_h(r\tau_h-1)+r))}\right\}$$

That is, the equilibria are:

$$S_h^* = -\frac{r(\beta_h+m_h)(a_h(K m_h\tau_h - K e_h) + m_h(m_h\tau_h+1))}{a_h m_h(K r a_h\tau_h + \beta_h + m_h)}$$
$$H_h^* = -\frac{r\tau_h(\beta_h+m_h)(a_h(K m_h\tau_h - K e_h) + m_h(m_h\tau_h+1))}{a_h(e_h(\beta_h+m_h) + m_h\tau_h(-\beta_h + m_h(r\tau_h-1)+r))} \qquad (3F)$$
$$C_{hh} = \frac{r^2\tau_h(\beta_h+m_h)(a_h(K m_h\tau_h - K e_h) + m_h(m_h\tau_h+1))^2}{2a_h m_h(e_h(\beta_h+m_h) + m_h\tau_h(-\beta_h + m_h(r\tau_h-1)+r))(K r a_h\tau_h + \beta_h + m_h)}$$
$$R_{h,I}^* = \frac{m_h(m_h\tau_h+1)(K r a_h\tau_h + \beta_h + m_h)}{a_h(e_h(\beta_h+m_h) + m_h\tau_h(-\beta_h + m_h(r\tau_h-1)+r))}$$

recalling that we refer to $R_{h,I}^*$ as the Interference $R^*$. $R_{h,I}^*$ is equivalent to equation (4) from the main text, with $i$ substituted with $h$ and $m_C = 0$ (here, contests do not increase mortality rate).

#### F.1.2 Analysis for Dove

The system of ODEs describing the consumer-resource dynamics for the dove is:

$$\frac{dS_d}{dt} = \frac{1}{\tau_d}(1+e_d)H_d - m_d S_d - a_d S_d R$$
$$\frac{dH_d}{dt} = a_d S_d R - m_d H_d - \frac{H_d}{\tau_d} \qquad (4F)$$
$$\frac{dR}{dt} = f(R) - a_d S_d R + m_d H_d$$

The absence of interference competition means there is no state variable describing Doves in conflict (i.e. there is no $C_{dd}$). The non-trivial equilibria are:

```
(*Solve for the equilibria*)
ConResEquilDove = Simplify[Solve[-ad*R*Sd + (Hd/τd) + (ed*Hd/τd) - md*Sd == 0 &&
                    ad*R*Sd - (Hd/τd) - md*Hd == 0 &&
                    r*(K - R) - ad*R*Sd + md*Hd == 0, {Sd, Hd, R}]];
(*Print the non-trivial equilibria*)
Part[ConResEquilDove, 2] // TraditionalForm
```

$$\left\{S_d \to r\left(-\frac{m_d\,\tau_d+1}{a_d}+\frac{K\,e_d}{m_d}-K\,\tau_d\right),\ H_d \to -\frac{r\,\tau_d\,(a_d\,(K\,m_d\,\tau_d-K\,e_d)+m_d\,(m_d\,\tau_d+1))}{a_d\,(e_d-m_d\,\tau_d)},\ R \to \frac{m_d\,(m_d\,\tau_d+1)}{a_d\,(e_d-m_d\,\tau_d)}\right\}$$

That is, the equilibria are:

$$S_d^* = r\left(-\frac{m_d\,\tau_d+1}{a_d}+\frac{K\,e_d}{m_d}-K\,\tau_d\right)$$
$$H_d^* = -\frac{r\,\tau_d\,(a_d\,(K\,m_d\,\tau_d-K\,e_d)+m_d\,(m_d\,\tau_d+1))}{a_d(e_d-m_d\,\tau_d)} \qquad (5\,F)$$
$$R_{d,E}^* = \frac{m_d(m_d\,\tau_d+1)}{a_d(e_d-m_d\,\tau_d)}$$

noting that $R_{d,E}^* = R_{d,I}^*$ for the dove. We use the former because it emphasizes that the equilibrium resource density set by the dove is also the dove's minimum resource requirement

### F.2 Calculation of Jacobian Matrices

#### F.2.1 Analysis for dove invader, resident hawk

First, we derive the Jacobian Matrix for the dove, $J_d$, where

$$J_d = \begin{pmatrix} \partial S_d\ /\ \partial S_d & \partial S_d\ /\ \partial H_d \\ \partial H_d\ /\ \partial S_d & \partial H_d\ /\ \partial H_d \end{pmatrix} \qquad (6\,F)$$

We consider $J_d$ when the dove is rare $S_d, H_d \to 0$ and the system is at the equilibrium set by the hawk, as defined by 3F. We calculate this below:

**(*This code will generate the Jacobian $J_d$,**
**and substitute the values $S_d$=0, $H_d$=0 and $S_h=S_h^*$, $H_h=H_h^*$, $C_{hh}=C_{hh}^*$, $R=R_{h,I}^*$ *)**

```
ConsumerDoveMatrix = {(*Put the dynamics of the dove into matrix form*)
    -ad * R * Sd + (Hd / τd) + (ed * Hd / τd) - md * Sd + ah * Hd * Sh,
     ad * R * Sd - (Hd / τd) - md * Hd - ah * Hd * Sh};

(*Calculate the Jacobian Matrix (Jd in the above equation)*)
JacMatDove = D[ConsumerDoveMatrix, {{Sd, Hd}}];

(*Take the limit when the dove is rare Sd=0, Hd=0 and Sh=S*h, Hh=H*h, Chh=C*hh, R=R*h,I *)
JacMatDove2 = Limit[JacMatDove, {Sd → 0, Hd → 0, Sh → Sstarh, Hh → Hstarh, R → RstarhI}];
(* note that we use the notation e.g. RstarhI for R*h,I for syntax reasons*)

(* print the Jacobian Matrix of Jd*)
JacMatDove2 // TraditionalForm
```

$$\begin{pmatrix} -a_d \, \text{Rstar}_{\text{hI}} - m_d & a_h \, \text{Sstar}_h + \frac{e_d}{\tau_d} + \frac{1}{\tau_d} \\ a_d \, \text{Rstar}_{\text{hI}} & -a_h \, \text{Sstar}_h - m_d - \frac{1}{\tau_d} \end{pmatrix}$$

That is,

$$J_d = \begin{pmatrix} -a_d \, R_{h,I}^* - m_d & a_h \, S_h^* + e_d \,/\, \tau_d + 1 \,/\, \tau_d \\ a_d \, R_{h,I}^* & -a_h \, S^* - m_d - 1 \,/\, \tau_d \end{pmatrix} \qquad (7\,F)$$

#### F.2.2 Analysis for Hawk invader, resident Dove

Now, we derive the Jacobian Matrix for the hawk, $J_h$:

$$J_h = \begin{pmatrix} \partial S_h \,/\, \partial S_h & \partial S_h \,/\, \partial H_h & \partial S_h \,/\, \partial C_{hh} \\ \partial H_h \,/\, \partial S_h & \partial H_h \,/\, \partial H_h & \partial H_h \,/\, \partial C_{hh} \\ \partial C_{hh} \,/\, \partial C_{hh} & \partial C_{hh} \,/\, \partial H_h & \partial C_{hh} \,/\, \partial C_{hh} \end{pmatrix} \qquad (8\,F)$$

We consider $J_h$ when the hawk is rare ($S_h \to 0$, $H_h \to 0$, $C_{hh} \to 0$) and the dove is at equilibrium with the resource as defined by 5F. This gives:

```
(*This code will generate the Jacobian J_h,
and substitute the values S_h=0,H_h=0,C_hh=0 and S_d=S*_d,H_d=H*_d,R=R*_{d,E}*)
ConsumerHawkMatrix = {(*Put the dynamics of the Hawk into matrix form*)
    -a_h*R*S_h + (H_h/τ_h) + (e_h*H_h/τ_h) - m_h*S_h - a_h*H_h*S_h - a_h*H_d*S_h + C_hh*β_h,
     a_h*R*S_h + a_h*H_d*S_h - (H_h/τ_h) - m_h*H_h - a_h*H_h*S_h + C_hh*β_h + m_h*C_hh,
     a_h*H_h*S_h - 2*β_h*C_hh - 2*m_h*C_hh};

(*Calculate the Jacobian Matrix (J_h in the above equation)*)
JacMatHawk = D[ConsumerHawkMatrix, {{S_h, H_h, C_hh}}];

(*Take the limit when the dove is rare S_d=0,H_d=0 and S_h=S*_h,H_h=H*_h,C_hh=C*_hh,R=R*_{h,I}*)
JacMatHawk2 = Limit[JacMatHawk, {S_h → 0, H_h → 0, C_hh → 0, S_d → Sstar_d, H_d → Hstar_d, R → Rstar_dE}];
(* we use the notation e.g. Rstar_dE for R*_{d,E} for syntax reasons*)

(* print the Jacobian Matrix of J_h*)
JacMatHawk2 // TraditionalForm
```

$$\begin{pmatrix} -a_h \, \mathrm{Hstar}_d - a_h \, \mathrm{Rstar}_{\mathrm{dE}} - m_h & \frac{e_h}{\tau_h} + \frac{1}{\tau_h} & \beta_h \\ a_h \, \mathrm{Hstar}_d + a_h \, \mathrm{Rstar}_{\mathrm{dE}} & -m_h - \frac{1}{\tau_h} & \beta_h + m_h \\ 0 & 0 & -2\beta_h - 2\, m_h \end{pmatrix}$$

That is,

$$J_h = \begin{pmatrix} -a_h H_d^* - a_h R_{d,E}^* - m_h & e_h / \tau_h + 1 / \tau_h & \beta_h \\ a_h H_d^* + a_h R_{d,E}^* & -m_h - 1/\tau_h & \beta_h + m_h \\ 0 & 0 & -\beta_h - 2\, m_h \end{pmatrix} \qquad (9F)$$

### F.3 Calculation of Eigenvalues

It is straightforward to calculate the eigenvalues for both dove and hawk invaders. We do so below.

#### F.3.1 Eigenvalues, Dove

We first calculate the eigenvalues for the dove from the $2 \times 2$ Jacobian matrix $J_d$:

```
(*Calculate both eigenvalues of 2 x 2 matrix*)
λd,1 = Part[Eigenvalues[JacMatDove2], 1]; λd,2 = Part[Eigenvalues[JacMatDove2], 2];

(*Show eigenvalues*)
{λd,1 // TraditionalForm, λd,2 // TraditionalForm}
```

$$\left\{\frac{1}{2\,\tau_d}\left(-\sqrt{4\,a_d\,e_d\,\tau_d\,\mathrm{Rstar}_{\mathrm{hI}} + 2\,a_d\,a_h\,\tau_d^2\,\mathrm{Sstar}_h\,\mathrm{Rstar}_{\mathrm{hI}} + a_h^2\,\tau_d^2\,\mathrm{Sstar}_h^2 + 2\,a_h\,\tau_d\,\mathrm{Sstar}_h + a_d^2\,\tau_d^2\,\mathrm{Rstar}_{\mathrm{hI}}^2 + 2\,a_d\,\tau_d\,\mathrm{Rstar}_{\mathrm{hI}} + 1} - a_h\,\tau_d\,\mathrm{Sstar}_h - a_d\,\tau_d\,\mathrm{Rstar}_{\mathrm{hI}} - 2\,m_d\,\tau_d - 1\right), \frac{1}{2\,\tau_d}\left(\sqrt{4\,a_d\,e_d\,\tau_d\,\mathrm{Rstar}_{\mathrm{hI}} + 2\,a_d\,a_h\,\tau_d^2\,\mathrm{Sstar}_h\,\mathrm{Rstar}_{\mathrm{hI}} + a_h^2\,\tau_d^2\,\mathrm{Sstar}_h^2 + 2\,a_h\,\tau_d\,\mathrm{Sstar}_h + a_d^2\,\tau_d^2\,\mathrm{Rstar}_{\mathrm{hI}}^2 + 2\,a_d\,\tau_d\,\mathrm{Rstar}_{\mathrm{hI}} + 1} - a_h\,\tau_d\,\mathrm{Sstar}_h - a_d\,\tau_d\,\mathrm{Rstar}_{\mathrm{hI}} - 2\,m_d\,\tau_d - 1\right)\right\}$$

That is, the eigenvalues are:

$$\lambda_{d,k} = \pm\frac{\sqrt{4\,a_d\,e_d\,\tau_d\,R^*_{h,I} + 2\,a_d\,a_h\,\tau_d^2\,S^*_h\,R^*_{h,I} + a_h^2\,\tau_d^2\,S^*_h\,2 + 2\,a_h\,\tau_d\,S^*_h + a_d^2\,\tau_d^2\,R^*_{h,I}\,2 + 2\,a_d\,\tau_d\,R^*_{h,I} + 1} - a_h\,\tau_d\,S^*_h - a_d\,\tau_d\,R^*_{h,I} - 2\,m_d\,\tau_d - 1}{2\,\tau_d}$$

Recall that the Dove will invade if there is at least one positive eigenvalue. $\lambda_{d,k}$ is always negative if the square root term is subtracted. Thus, we focus on when the root term is added (see below)

#### F.3.2 Eigenvalues, Hawk

We calculate the eigenvalues for the invading hawk from the $3 \times 3$ Jacobian matrix $J_h$:

```
(*Calculate all 3 eigenvalues of 3 x 3 matrix*)
λh,1 = Part[Eigenvalues[JacMatHawk2], 1];
λh,2 = Part[Eigenvalues[JacMatHawk2], 2];
λh,3 = Part[Eigenvalues[JacMatHawk2], 3];

(*Show eigenvalues*)
{TraditionalForm[λh,1], TraditionalForm[λh,2], TraditionalForm[λh,3]}
```

$$\left\{-2\,(\beta_h + m_h), \frac{1}{2\,\tau_h}\left(-\sqrt{\left(2\,a_h^2\,\mathrm{Hstar}_d\,\mathrm{Rstar}_{\mathrm{dE}}\,\tau_h^2 + 4\,a_h\,\mathrm{Hstar}_d\,e_h\,\tau_h + a_h^2\,\mathrm{Hstar}_d^2\,\tau_h^2 + 2\,a_h\,\mathrm{Hstar}_d\,\tau_h + 4\,a_h\,\mathrm{Rstar}_{\mathrm{dE}}\,e_h\,\tau_h + a_h^2\,\mathrm{Rstar}_{\mathrm{dE}}^2\,\tau_h^2 + 2\,a_h\,\mathrm{Rstar}_{\mathrm{dE}}\,\tau_h + 1\right)} - a_h\,\mathrm{Hstar}_d\,\tau_h - a_h\,\mathrm{Rstar}_{\mathrm{dE}}\,\tau_h - 2\,m_h\,\tau_h - 1\right), \frac{1}{2\,\tau_h}\left(\sqrt{\left(2\,a_h^2\,\mathrm{Hstar}_d\,\mathrm{Rstar}_{\mathrm{dE}}\,\tau_h^2 + 4\,a_h\,\mathrm{Hstar}_d\,e_h\,\tau_h + a_h^2\,\mathrm{Hstar}_d^2\,\tau_h^2 + 2\,a_h\,\mathrm{Hstar}_d\,\tau_h + 4\,a_h\,\mathrm{Rstar}_{\mathrm{dE}}\,e_h\,\tau_h + a_h^2\,\mathrm{Rstar}_{\mathrm{dE}}^2\,\tau_h^2 + 2\,a_h\,\mathrm{Rstar}_{\mathrm{dE}}\,\tau_h + 1\right)} - a_h\,\mathrm{Hstar}_d\,\tau_h - a_h\,\mathrm{Rstar}_{\mathrm{dE}}\,\tau_h - 2\,m_h\,\tau_h - 1\right)\right\}$$

The eigenvalues of $J_h$ are:

$$\lambda_{h,1} = -2\,(\beta_h + m_h)$$

$$\lambda_{h,2} = \frac{-\sqrt{4\,a_h\,H_d^*\,e_h\,\tau_h + 4\,a_h\,H_d^*\,\tau_h + 4\,a_h\,R_{d,E}^*\,e_h\,\tau_h + a_h^2\,(R_{d,E}^*)^2\,\tau_h^2 + 2\,a_h\,R_{d,E}^*\,\tau_h + 1} - a_h\,R_{d,E}^*\,\tau_h - 2\,m_h\,\tau_h - 1}{2\,\tau_h}$$

$$\lambda_{h,2} = \frac{\sqrt{4\,a_h\,H_d^*\,e_h\,\tau_h + 4\,a_h\,H_d^*\,\tau_h + 4\,a_h\,R_{d,E}^*\,e_h\,\tau_h + a_h^2\,(R_{d,E}^*)^2\,\tau_h^2 + 2\,a_h\,R_{d,E}^*\,\tau_h + 1} - a_h\,R_{d,E}^*\,\tau_h - 2\,m_h\,\tau_h - 1}{2\,\tau_h}$$

The hawk invades if there is at least one positive eigenvalue. $\lambda_{h,1}$ and $\lambda_{h,2}$ are clearly always negative. Thus, we examine the conditions in which $\lambda_{h,3}>0$.

### F.4 Rearrangement of Eigenvalues into Invasion Criteria

#### F.4.1 Dove Invasion Criterion

Here, we solve for when $\lambda_{d,2} > 0$ in terms of $R_{h,I}^*$. This yields the invasion criterion.

```
(* Find when λd,2>0 in terms of R*h,I (RstarhI) *)
(* parameters and equilibria are all positive *)
InvCritDove = Reduce[λd,2 > 0 && ad > 0 && ed > 0 && τd > 0 &&
    RstarhI > 0 && md > 0 && ed > md * τd && Sstarh > 0 && ah > 0, {RstarhI}, Reals];
Part[InvCritDove, 7]
```

$$\text{Rstar}_{hI} > \frac{-m_d - m_d^2\,\tau_d - a_h\,m_d\,\text{Sstar}_h\,\tau_d}{-a_d\,e_d + a_d\,m_d\,\tau_d}$$

Multiplying the numerator and denominator of the right hand side by $-1$, the dove invades if

$$R_{h,I}^* > \frac{m_d + m_d^2\,\tau_d + a_h\,m_d\,\tau_d\,S_h^*}{a_d(e_d - m_d\,\tau_d)}$$

The right hand side can rearranged as

$$\begin{aligned}\frac{m_d + m_d^2\,\tau_d + a_h\,m_d\,\tau_d\,S_h^*}{a_d(e_d - m_d\,\tau_d)} &= \frac{m_d(m_d\,\tau_d + 1)}{a_d(e_d - m_d\,\tau_d)} + \frac{a_h\,m_h\,\tau_h\,S_h^*}{a_d(e_d - m_d\,\tau_d)} \\ &= \frac{m_d(m_d\,\tau_d + 1)}{a_d(e_d - m_d\,\tau_d)}\left(1 + \frac{a_h\,\tau_d\,S_h^*}{m_d\,\tau_d + 1}\right) \\ &= R_{d,E}^*\left(1 + \frac{a_h\,S_h^*}{m_d + 1/\tau_d}\right)\end{aligned}$$

The inequality above can be rewritten as

$$R_{h,I}^* > R_{d,E}^*\left(1 + \frac{a_h\,S_h^*}{m_d + 1/\tau_d}\right)$$

Dividing each side by $\left(1 + \frac{a_h\,S_h^*}{m_d + 1/\tau_d}\right)$ yields:

$$R^*_{d,E} < R^*_{h,I}\,\frac{1}{1+\frac{a_h\,S^*_h}{m_d+1/\tau_d}} \qquad (10\,F)$$

In the main text we substitute

$$P'_d = \frac{1}{1+\frac{a_h\,S^*_h}{m_d+1/\tau_d}}$$

Thus, the dove invades if

$$R^*_{d,E} < R^*_{h,I}\,P'_d \qquad (11\,F)$$

which is the same as equation 5 from the main text.

#### F.4.2 Hawk Invasion Criterion

Here, we solve for when $\lambda_{h,3} > 0$ in terms of $R^*_{d,E}$. This yields:

```
(* Find when λ2>0 in terms of R*_{d,E} (Rstar_dE) *)
(* parameters and equilibria are all positive; the last inequality ensures that the most general form is output *)
InvCritHawk = Reduce[λh,3 > 0 && ah > 0 && eh > 0 && τh > 0 &&
    RstardE > 0 && mh > 0 && eh > mh * τh && 0 < Hstard ≤ (−mh − mh^2 τh)/(−ah eh + ah mh τh), {RstardE}, Reals];
Part[InvCritHawk, 6]
```

$$\mathrm{Rstar_{dE}} > \frac{\mathrm{a_h\,e_h\,Hstar_d - m_h - a_h\,Hstar_d\,m_h\,\tau_h - m_h^2\,\tau_h}}{\mathrm{-a_h\,e_h + a_h\,m_h\,\tau_h}}$$

The hawk can invade if

$$R^*_{d,E} > \frac{a_h\,H^*_d\,e_h - a_h\,H^*_d\,m_h\,\tau_h + m_h^2\,(-\tau_h) - m_h}{a_h\,m_h\,\tau_h - a_h\,e_h}$$

The right-hand side term can be rearranged as

$$\frac{a_h\,H^*_d\,e_h - a_h\,H^*_d\,m_h\,\tau_h - \tau_h\,m_h^2 - m_h}{a_h\,m_h\,\tau_h - a_h\,e_h} = \frac{m_h(m_h\,\tau_h+1)}{a_h(e_h - m_h\,\tau_h)} - H^*_d\,\frac{a_H(e_H - m_H\,\tau_H)}{a_H(e_H - m_H\,\tau_H)} = R^*_{h,E} - H^*_d$$

Substituting this value into the inequality and adding $H^*_d$ to both sides gives

$$R^*_{h,E} < R^*_{d,E} + H^*_d \qquad (12\,F)$$

which is the same as equation 6 in the main text.

F.4.3 Coexistence Criteria

By dividing each side of equation 11F by $R^*_{h,E}$ and multiplying each side of 12F by $\frac{R^*_{h,E}/R^*_{d,E}}{R^*_{d,E}+H^*_d}$, we derive the expression:

$$\frac{1}{1 + H_d^* / R_{d,E}^*} < \frac{R_{d,E}^*}{R_{h,E}^*} < \frac{R_{h,I}^*}{R_{h,E}^*} P_d' \qquad (13\,F)$$

which is the same as equation (7) from the main text.

### F.5 Interpretation of $P_d'$

In the main text, it is claimed that $P_d'$ is the probability that a dove does not experience interference from a hawk while handling a resource. To see this, recall that constant rates in Ordinary Differential Equations (e.g., $d N / d t = -k * N$) are the deterministic mass action limit of exponentially distributed times to events. If two random variables, A and B, are exponentially distributed with rates a and b, P[A happens before B] = a/(a+b). In our context, B represents the initiation of hawk interference, and A represents all the other ways in which doves exit the Handling class. The per-dove rate of A is $m_d + 1 / \tau_d$. The per-dove rate of B is $a_h S_h^*$ (the search rate of hawks multiplied by the equilibrium abundance of hawk searchers). It then follows that

$$P_d' = \frac{1}{1 + \frac{a_h S_h^*}{m_d + \left(\frac{1}{\tau_d}\right)}} = \frac{m_d + \frac{1}{\tau_d}}{m_d + \frac{1}{\tau_d} + a_h S_h^*} = \text{Pr[dove does not experience interference].}$$

### Section G: Invasion Criteria, General Case (Both Consumers Interfere)

#### Introduction:

In this Section, we derive invasion criteria for the general case in which both consumers interfere $(\beta_i,\ \beta_j < \infty)$. We show the derivation of equations 10-11. In addition, we provide explanations for several claims we make in the main text pertaining to the derivation and interpretation of terms in the invasion criteria (the derivation and interpretation of $P_j$, $\omega_{ji}$, and $\beta_0$).

As noted in the main text, we employ invasion analysis. To do so, we derive the Jacobian Matrix of the system when a single consumer (the resident) is at equilibrium and the other consumer (the invader) is at low abundance. If any eigenvalues of the Jacobian Matrix are positive, it implies the invader tends to increase in abundance when rare. If both consumers do so, it implies mutual invisibility (i.e., coexistence).

### G.1 Consumer-Resource Equilibrium Calculation

#### G.1.1 Model when both types interfere

As shown in the main text, the full dynamics of the system are given by:

$$\begin{aligned}
\frac{dS_i}{dt} &= \frac{1}{\tau_i}(1+e_i)H_i + 2\beta_i C_{ii} + \beta_i\left(C_{ij}+C_{ji}\right) - m_i S_i - a_i S_i \sum_{j=1}^{2} H_j - a_i S_i R \\
\frac{dH_i}{dt} &= a_i S_i R + (2\beta_i + 2m_i + 2m_{Ci})C_{ii} + \left(\beta_j + m_j + m_{Cj}\right)\left(C_{ij}+C_{ji}\right) - m_i H_i - \frac{H_i}{\tau_i} - H_i \sum_{j=1}^{2} a_j S_j \\
\frac{dC_{ii}}{dt} &= a_i S_i H_i - (2\beta_i + 2m_i + 2m_{Ci})C_{ii} \qquad (1G)\\
\frac{dC_{ij}}{dt} &= a_j S_j H_i - C_{ij}\left(\beta_i + \beta_j + m_i + m_j + m_{Ci} + m_{Cj}\right) \\
\frac{dR}{dt} &= f(R) - R\sum_{j=1}^{2} a_j S_j + \sum_{j=1}^{2} m_j H_j
\end{aligned}$$

where $i, j = 1, 2$ and $i \neq j$. $f(R) = r(K-R)$. See the description of the model in the main text for more details.

Below, we derive the equilibrium of the Consumer-Resource system (which contains a single non-trivial equilibrium) for consumer i (i.e., in the absence of consumer j). For simplicity, we examine when $e_i = e_j = e$ and $m_i = m_j = m$.

#### G.1.2 1-consumer-1-resource system

We solve for the equilibria of the set of Ordinary Differential Equations describing the consumer-resource dynamics of a single type with the resource. We consumer type 1 the resident and type 2 the invader. 1 and 2 are used instead of $i$ and $j$ because they are less easily visually confused. However, the same results apply.

The dynamics of the 1-consumer-1-resource system are:

$$\frac{dS_1}{dt} = \frac{1}{\tau_1}(1+e_1)H_1 + 2\beta_1 C_{11} - m_1 S_1 - a_1 S_1 H_1 - a_1 S_1 R$$

$$\frac{dH_1}{dt} = a_1 S_1 R + (2\beta_1 + 2m_1) C_{11} - m_1 H_1 - \frac{H_1}{\tau_1} - a_1 S_1 H_1 \qquad (2G)$$

$$\frac{dC_{11}}{dt} = a_1 S_1 H_1 - (2\beta_1 + 2m_1) C_{11}$$

$$\frac{dR}{dt} = f(R) - a_1 S_1 R + m H_1$$

From which the non-trivial equilibrium of the system is:

```
(*Solve for the equilibria*)
ConResEquilC1 = Simplify[Solve[-a1*R*S1 + (H1/τ1) + (e1*H1/τ1) - m1*S1 + 2 C11*β1 - a1*H1*S1 == 0 &&
                          a1*R*S1 - (H1/τ1) - m1*H1 + 2 C11*β1 + 2 m1*C11 - a1*H1*S1 == 0 &&
                          a1*H1*S1 - 2*β1*C11 - 2*m1*C11 == 0 &&
                          r*(K - R) - a1*R*S1 + m1*H1 == 0, {S1, H1, C11, R}]];

(*Print the non-trivial equilibria*)
Part[ConResEquilC1, 2] // TraditionalForm
```

$$\left\{S_1 \to -\frac{r(\beta_1 + m_1)(a_1(K m_1 \tau_1 - e_1 K) + m_1(m_1 \tau_1 + 1))}{a_1 m_1 (a_1 K r \tau_1 + \beta_1 + m_1)}, H_1 \to -\frac{r \tau_1 (\beta_1 + m_1)(a_1(K m_1 \tau_1 - e_1 K) + m_1(m_1 \tau_1 + 1))}{a_1(e_1(\beta_1 + m_1) + m_1 \tau_1(-\beta_1 + m_1(r\tau_1 - 1) + r))},\right.$$
$$C_{11} \to \frac{r^2 \tau_1 (\beta_1 + m_1)(a_1(K m_1 \tau_1 - e_1 K) + m_1(m_1 \tau_1 + 1))^2}{2 a_1 m_1 (e_1(\beta_1 + m_1) + m_1 \tau_1(-\beta_1 + m_1(r\tau_1 - 1) + r))(a_1 K r \tau_1 + \beta_1 + m_1)},$$
$$\left. R \to \frac{m_1(m_1 \tau_1 + 1)(a_1 K r \tau_1 + \beta_1 + m_1)}{a_1(e_1(\beta_1 + m_1) + m_1 \tau_1(-\beta_1 + m_1(r\tau_1 - 1) + r))}\right\}$$

That is:

$$S_1^* = \frac{r(\beta_1 + m_1)(a_1(K m_1 \tau_1 - e_1 K) + m_1(m_1 \tau_1 + 1))}{a_1 m_1 (a_1 K r \tau_1 + \beta_1 + m_1)}$$

$$H_1^* = -\frac{r \tau_1 (\beta_1 + m_1)(a_1(K m_1 \tau_1 - e K) + m_1(m_1 \tau_1 + 1))}{a_1\left(\beta_1(e_1 - m_1 \tau_1) + m_1\left(e_1 + m r \tau_1^2 + \tau_1(r - m_1)\right)\right)} \qquad (3G)$$

$$C_{11} = \frac{r^2 \tau_1 (\beta_1 + m_1)(a_1(K m_1 \tau_1 - e_1 K) + m_1(m_1 \tau_1 + 1))^2}{2 a_1 m_1 \left(\beta_1(e_1 - m_1 \tau_1) + m_1\left(e_1 + m_1 r \tau_1^2 + \tau_1(r - m_1)\right)\right)(a_1 K r \tau_1 + \beta_1 + m_1)}$$

$$R_{1,I}^* \frac{m_1(m_1 \tau_1 + 1)(a_1 K r \tau_1 + \beta_1 + m_1)}{a_1\left(\beta_1(e_1 - m_1 \tau_1) + m_1\left(e_1 + m_1 r \tau_1^2 + \tau_1(r - m_1)\right)\right)}$$

### G.2 Calculation of Jacobian Matrix

We calculate the Jacobian Matrix for consumer 2, the invader. The Jacobian matrix is given by:

$$J_2 = \begin{pmatrix} \partial S_2 / \partial S_2 & \partial S_2 / \partial H_2 & \partial S_2 / \partial C_{22} & \partial S_2 / \partial C_{\{12\}} \\ \partial H_2 / \partial S_2 & \partial H_2 / \partial H_2 & \partial H_2 / \partial C_{22} & \partial H_2 / \partial C_{\{12\}} \\ \partial C_{22} / \partial S_2 & \partial C_{22} / \partial H_2 & \partial C_{22} / \partial C_{22} & \partial C_{22} / \partial C_{\{12\}} \\ \partial C_{\{12\}} / \partial S_2 & \partial C_{\{12\}} / \partial H_2 & C_{\{12\}} / \partial H_2 & \partial C_{\{12\}} / \partial C_{\{12\}} \end{pmatrix}$$

where $C_{\{12\}} = C_{12} + C_{21}$.

We consider $J_2$ when the type 2 is rare $S_2, H_2, C_{22}, C_{\{12\}} \to 0$ and the system is at the equilibrium set by the hawk, as defined by 3G. We calculate this below:

```
(*This code will generate the Jacobian J2,
and substitute the values S2=0,H2=0,C22=0,C{12}=0 and S1=S1*,H1=H1*,C11=C11*,R=R1,I*)
ConsumerC2Matrix = { (*Put the dynamics of consumer type 2 into matrix form*)
        -a2*R*S2 +(H2/τ2) +(e2*H2/τ2) -m2*S2 -a2*H1*S2 -a2*H2*S2 +2 C22*β2 +C{12}*β2,

    a2*R*S2 -(H2/τ2) -m2*H2 -a1*H2*S1 -  a2*H2*S2 +2 C22*β2 +2 m2*C22 +C{12}*β1 +C{12}*m1,
        a2*H2*S2 -2*β2*C22 -2*m2*C22,
        a2*H1*S2 + a1*H2*S1 -β1*C{12} -β2*C{12} -(m1 + m2)*C{12}};

(*Calculate the Jacobian Matrix (J2  in the above equation)*)
JacMatC = ResourceFunction["JacobianMatrix"][ConsumerC2Matrix, {S2, H2, C22, C{12}}];

(*Take the limit when type 2 is rare  S2=0,H2=0,C22=0,C{12}=0
  and 1 is at equilibrium S1=S1*,H1=H1*,C11=C11*,R=R1,I*)
JacMatC2 = Limit[JacMatC, {S2 → 0, H2 → 0, C22 → 0, C12 → 0, S1 → Sstar1, H1 → Hstar1, R → Rstar1I}];

(* print the Jacobian Matrix of J2*)JacMatC2 // TraditionalForm
```

$$\begin{pmatrix} -a_2\,\text{Hstar}_1 + a_2\,(-\text{Rstar1I}) - m_2 & \frac{e_2}{\tau_2} + \frac{1}{\tau_2} & 2\beta_2 & \beta_2 \\ a_2\,\text{Rstar1I} & -a_1\,\text{Sstar}_1 - m_2 - \frac{1}{\tau_2} & 2\beta_2 + 2m_2 & \beta_1 + m_1 \\ 0 & 0 & -2\beta_2 - 2m_2 & 0 \\ a_2\,\text{Hstar}_1 & a_1\,\text{Sstar}_1 & 0 & -\beta_1 - \beta_2 - m_1 - m_2 \end{pmatrix}$$

That is,

$$J_2 = \begin{pmatrix} -a_2 H_1^* - a_2 R_{1,I}^* - m_2 & e_2/\tau_2 + 1/\tau_2 & 2\beta_2 & \beta_2 \\ a_2 R_{1,I}^* & -a_1 S_1^* - m_2 - 1/\tau_2 & 2\beta_2 + 2m_2 & \beta_1 + m_1 \\ 0 & 0 & -2\beta_2 - 2m_2 & 0 \\ a_2 H_1^* & a_1 S_1^* & 0 & -\beta_1 - \beta_2 - m_1 + m_2 \end{pmatrix}$$

### G.3 Invasion Analysis

In contrast to Supplementary Material Sections D and F we were unable to find an analytically tractable solution for the eigenvalues ($\lambda_k$, $k = 1, 2, 3, 4$) of 2. Instead, we use a combination of numerical and analytical arguments to determine if any eigenvalue of the Jacobian matrix (henceforth) is positive. We take advantage of the property that $\text{Det}(J_2) = \prod_{k=1}^{4} \lambda_k$ (the determinant of a matrix is equal to the product of its eigenvalues). Based on the characteristic equation of $J_2$, it is easy to see that one eigenvalue is always negative, at it is always equal to $-2\,\beta_j - 2\,m_j$. Numerically, we find that two additional eigenvalues of the matrix are always negative throughout feasible parameter space, as depicted in Fig. G1. However, we find a single eigenvalue, henceforth $\lambda_4$, is either positive

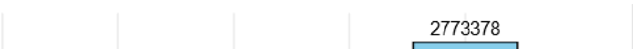

or negative depending on the parameter choices. Thus, $J_j$always contains three negative eigenvalues and a fourth eigenvalue that varies in sign. If all four eigenvalues are negative, $\text{Det}(J_j) > 0$and $\text{Det}(J_j) < 0$ if $\lambda_4 > 0$. We therefore conjecture that consumer j invades if $\text{Det}(J_j) < 0$. See "Supplementary_Material_G_Eigenvalues.R" for the code that generated Fig. G1.

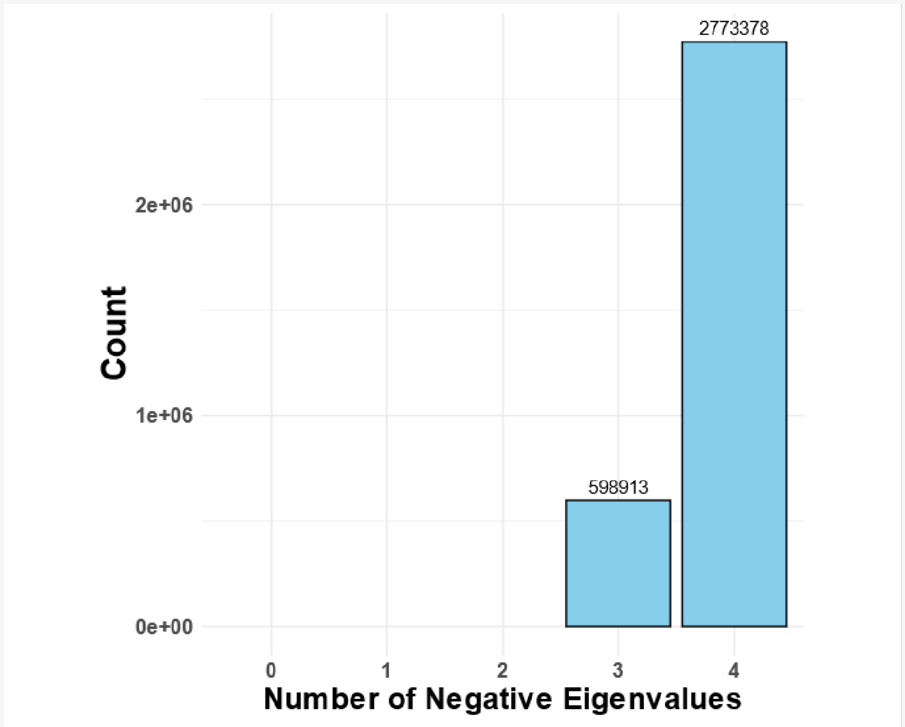


**Fig. G1:** The Jacobian Matrix always has either 3 or 4 negative eigenvalues. We numerically calculated the eigenvalues of the Jacobian Matrix using Latin Hypercube Sampling to sample from an extensive range of parameter space wherein: $a_i$~{.01,100}, $a_j$~{.01,100}, $\tau_i$~{.001,100}, $\tau_j$~{.001,100}, $\beta_i$~{.01,1000}, $\beta_j$~{.01,1000}, $e$~{.01,100}, $m$~{.0001,10}, $r$~{.01,1000} and $K$~{5,5000}. Sampling was performed on a log scale. See "Supplementary_Material_G_Eigenvalues.R" for more details.

Below, we show algebraic manipulation of Det($J_2$) into equations 10-11 from the main text.

#### G.3.1 Calculation and algebraic treatment of the determinant

Here, we take the Determinant of $J_2$ and derive the invasion criterion of type 2. The determinant is:

```
(* Take the determinant)*)
Det[JacMatC2] // TraditionalForm
```

$$(-2\beta_2 - 2 m_2)\left(\frac{a_2 \beta_1 e_2 \text{Hstar}_1}{\tau_2} + \frac{a_2 e_2 \text{Hstar}_1 m_1}{\tau_2} + \frac{a_2 e_2 m_2 \text{Rstar1I}}{\tau_2} + \frac{a_2 e_2 m_1 \text{Rstar1I}}{\tau_2} + \frac{a_2 \beta_1 e_2 \text{Rstar1I}}{\tau_2} + \frac{a_2 \beta_2 e_2 \text{Rstar1I}}{\tau_2} - a_2 \beta_1 \text{Hstar}_1 m_2 - a_1 a_2 \text{Hstar}_1 m_2 \text{Sstar}_1 - \frac{a_2 \text{Hstar}_1 m_2}{\tau_2} - a_2 \text{Hstar}_1 m_2^2 - a_2 \text{Hstar}_1 m_1 m_2 - a_2 \beta_1 m_2 \text{Rstar1I} - a_2 \beta_2 m_2 \text{Rstar1I} - a_1 a_2 m_2 \text{Rstar1I}\, \text{Sstar}_1 - a_2 m_2^2 \text{Rstar1I} - a_2 m_1 m_2 \text{Rstar1I} - a_1 \beta_2 m_2 \text{Sstar}_1 - a_1 m_2^2 \text{Sstar}_1 - \frac{\beta_1 m_2}{\tau_2} - \frac{\beta_2 m_2}{\tau_2} - \beta_1 m_2^2 - \beta_2 m_2^2 - \frac{m_2^2}{\tau_2} - \frac{m_1 m_2}{\tau_2} - m_2^3 - m_1 m_2^2\right)$$

where consumer 2 invades if

$$(-2\beta_2 - 2 m_2)\left(\frac{a_2 \beta_1 e_2 \text{Hstar}_1}{\tau_2} + \frac{a_2 e_2 \text{Hstar}_1 m_1}{\tau_2} + \frac{a_2 e_2 m_2 \text{Rstar1I}}{\tau_2} + \frac{a_2 e_2 m_1 \text{Rstar1I}}{\tau_2} + \frac{a_2 \beta_1 e_2 \text{Rstar1I}}{\tau_2} + \frac{a_2 \beta_2 e_2 \text{Rstar1I}}{\tau_2} - a_2 \beta_1 \text{Hstar}_1 m_2 - a_1 a_2 \text{Hstar}_1 m_2 \text{Sstar}_1 - \frac{a_2 \text{Hstar}_1 m_2}{\tau_2} - a_2 \text{Hstar}_1 m_2^2 - a_2 \text{Hstar}_1 m_1 m_2 - a_2 \beta_1 m_2 \text{Rstar1I} - a_2 \beta_2 m_2 \text{Rstar1I} - a_1 a_2 m_2 \text{Rstar1I}\, \text{Sstar}_1 - a_2 m_2^2 \text{Rstar1I} - a_2 m_1 m_2 \text{Rstar1I} - a_1 \beta_2 m_2 \text{Sstar}_1 - a_1 m_2^2 \text{Sstar}_1 - \frac{\beta_1 m_2}{\tau_2} - \frac{\beta_2 m_2}{\tau_2} - \beta_1 m_2^2 - \beta_2 m_2^2 - \frac{m_2^2}{\tau_2} - \frac{m_1 m_2}{\tau_2} - m_2^3 - m_1 m_2^2\right) < 0$$

Note: to make the below notation less cumbersome, we below substitute:

$\text{Hstar}_1 = H_1^* = H_1$
$\text{Sstar}_1 = S_1^* = S_1$
$\text{Rstar1I} = R_{1,I}^* = \text{R1I}$

Note that the initial term of Det($J_2$), $(-2\beta_2 - 2\,\mathrm{m}_2)$, is always negative. Thus, dividing each side of Det($J_2$)<0 by $(-2\beta_2 - 2\,\mathrm{m}_2)$ yields the expression:

$$\frac{a_2 \beta_1 e_2 H_1}{\tau_2} + \frac{a_2 e_2 H_1 m_1}{\tau_2} + \frac{a_2 e_2 m_2 R1I}{\tau_2} + \frac{a_2 e_2 m_1 R1I}{\tau_2} + \frac{a_2 \beta_1 e_2 R1I}{\tau_2} + \frac{a_2 \beta_2 e_2 R1I}{\tau_2} - a_2 \beta_1 H_1 m_2 - a_1 a_2 H_1 m_2 S_1 - \frac{a_2 H_1 m_2}{\tau_2} - a_2 H_1 m_2^2 - a_2 H_1 m_1 m_2 - a_2 \beta_1 m_2 R1I - a_2 \beta_2 m_2 R1I - a_1 a_2 m_2 R1I S_1 - a_2 m_2^2 R1I - a_2 m_1 m_2 R1I - a_1 \beta_2 m_2 S_1 - a_1 m_2^2 S_1 - \frac{\beta_1 m_2}{\tau_2} - \frac{\beta_2 m_2}{\tau_2} - \beta_1 m_2^2 - \beta_2 m_2^2 - \frac{m_2^2}{\tau_2} - \frac{m_1 m_2}{\tau_2} - m_2^3 - m_1 m_2^2 > 0$$

noting that the inequality was reversed because both sides were divided by a negative number $-2\beta_2 - 2m$. It is then convenient to separate terms containing R1I. Hence

**(* Separate expression in terms of R1I*)**

$$\textbf{Collect}\Big[\textbf{Simplify}\Big[\Big(\frac{a_2\beta_1 e_2 H_1}{\tau_2}+\frac{a_2 e_2 H_1 m_1}{\tau_2}+\frac{a_2 e_2 m_2\,\mathrm{R1I}}{\tau_2}+\frac{a_2 e_2 m_1\,\mathrm{R1I}}{\tau_2}+\frac{a_2\beta_1 e_2\,\mathrm{R1I}}{\tau_2}+\frac{a_2\beta_2 e_2\,\mathrm{R1I}}{\tau_2}-a_2\beta_1 H_1 m_2-a_1 a_2 H_1 m_2 S_1-\frac{a_2 H_1 m_2}{\tau_2}-a_2 H_1 m_2^2-a_2 H_1 m_1 m_2-a_2\beta_1 m_2\,\mathrm{R1I}-a_2\beta_2 m_2\,\mathrm{R1I}-a_1 a_2 m_2\,\mathrm{R1I}\,S_1-a_2 m_2^2\,\mathrm{R1I}-a_2 m_1 m_2\,\mathrm{R1I}-a_1\beta_2 m_2 S_1-a_1 m_2^2 S_1-\frac{\beta_1 m_2}{\tau_2}-\frac{\beta_2 m_2}{\tau_2}-\beta_1 m_2^2-\beta_2 m_2^2-\frac{m_2^2}{\tau_2}-\frac{m_1 m_2}{\tau_2}-m_2^3-m_1 m_2^2\Big)\Big],\mathrm{R1I}\Big]\ \textbf{// TraditionalForm}$$

$$\frac{1}{\tau_2}\big(a_2\beta_1 e_2 H_1+a_2 e_2 H_1 m_1-a_2 H_1 m_2(a_1 S_1\tau_2+\beta_1\tau_2+m_1\tau_2+m_2\tau_2+1)-m_2\big(m_2(a_1 S_1\tau_2+\beta_1\tau_2+\beta_2\tau_2+1)+a_1\beta_2 S_1\tau_2+\beta_1+\beta_2+m_2^2\tau_2+m_1(m_2\tau_2+1)\big)\big)+\frac{\mathrm{R1I}\,(a_2\beta_1 e_2+a_2\beta_2 e_2+a_2 e_2 m_1+a_2 e_2 m_2-a_2 m_2\tau_2(a_1 S_1+\beta_1+\beta_2+m_1+m_2))}{\tau_2}$$

or

$$\frac{a_2\beta_1 e_2 H_1+a_2 e_2 H_1 m_1-a_2 H_1 m_2(a_1 S_1\tau_2+\beta_1\tau_2+m_1\tau_2+m_2\tau_2+1)-m_2\big(m_2(a_1 S_1\tau_2+\beta_1\tau_2+\beta_2\tau_2+1)+a_1\beta_2 S_1\tau_2+\beta_1+\beta_2+m_2^2\tau_2+m_1(m_2\tau_2+1)\big)}{\tau_2}>-\mathrm{R1I}\frac{(a_2\beta_1 e_2+a_2\beta_2 e_2+a_2 e_2 m_1+a_2 e_2 m_2-a_2 m_2\tau_2(a_1 S_1+\beta_1+\beta_2+m_1+m_2))}{\tau_2}$$

after some rearranging. Dividing each side by $-\frac{(a_2\beta_1 e_2+a_2\beta_2 e_2+a_2 e_2 m_1+a_2 e_2 m_2-a_2 m_2\tau_2(a_1 S_1+\beta_1+\beta_2+m_1+m_2))}{\tau_2}$ simplifies the equation. Note that we do not pay attention to the sign of this expression (which matters for the direction of the inequality). This is because we next multiply by an expression with the same sign (in which case, the inequality will not change). Dividing by the expression gives

**(* Divide left hand side by fraction: *)**

$$\textbf{Simplify}\left[\frac{\frac{a_2\beta_1 e_2 H_1+a_2 e_2 H_1 m_1-a_2 H_1 m_2(a_1 S_1\tau_2+\beta_1\tau_2+m_1\tau_2+m_2\tau_2+1)-m_2(m_2(a_1 S_1\tau_2+\beta_1\tau_2+\beta_2\tau_2+1)+a_1\beta_2 S_1\tau_2+\beta_1+\beta_2+m_2^2\tau_2+m_1(m_2\tau_2+1))}{\tau_2}}{-\frac{(a_2\beta_1 e_2+a_2\beta_2 e_2+a_2 e_2 m_1+a_2 e_2 m_2-a_2 m_2\tau_2(a_1 S_1+\beta_1+\beta_2+m_1+m_2))}{\tau_2}}\right]\ \textbf{// TraditionalForm}$$

$$\big(a_2 H_1(m_2(a_1 S_1\tau_2+\beta_1\tau_2+m_1\tau_2+m_2\tau_2+1)-e_2(\beta_1+m_1))+m_2\big(m_2(a_1 S_1\tau_2+\beta_1\tau_2+\beta_2\tau_2+1)+a_1\beta_2 S_1\tau_2+\beta_1+\beta_2+m_2^2\tau_2+m_1(m_2\tau_2+1)\big)\big)/(a_2(e_2(\beta_1+\beta_2+m_1+m_2)-m_2\tau_2(a_1 S_1+\beta_1+\beta_2+m_1+m_2)))$$

The algebra is made simpler by making the substitution $g_i=1/\tau_i,\ i=1,\ 2$. This yields

$$\frac{a_2 H_1(e_2 g_2(\beta_1+m_1)-m_2(a_1 S_1+\beta_1+g_2+m_1+m_2))-m_2(a_1 m_2 S_1+a_1\beta_2 S_1+g_2(\beta_1+\beta_2+m_1+m_2)+\beta_1 m_2+\beta_2 m_2+m_2^2+m_1 m_2)}{a_2(a_1 m_2 S_1+(\beta_1+\beta_2+m_1+m_2)(m_2-e_2 g_2))}>\mathrm{R1I}$$

It is then convenient to multiply each side of the equation by $\frac{a_1 m_2 S_1}{e_2 g_2 - m_2} - (\beta_1 + \beta_2 + m_1 + m_2)$. Note that it is relatively straightforward to show that this expression $\frac{a_1 m_2 S_1}{e_2 g_2 - m_2} - (\beta_1 + \beta_2 + m_1 + m_2)$ has the same sign as the above expression

$$-\frac{(a_2 \beta_1 e_2 + a_2 \beta_2 e_2 + a_2 e_2 m_1 + a_2 e_2 m_2 - a_2 m_2 \tau_2 (a_1 S_1 + \beta_1 + \beta_2 + m_1 + m_2))}{\tau_2}$$

that we previously divided both sides by. Thus the direction of the inequality remains the same. This yields:

$$\frac{a_2 H_1 (e_2 g_2 (\beta_1 + m_1) - m_2 (a_1 S_1 + \beta_1 + g_2 + m_1 + m_2)) - m_2 (a_1 m_2 S_1 + a_1 \beta_2 S_1 + g_2 (\beta_1 + \beta_2 + m_1 + m_2) + \beta_1 m_2 + \beta_2 m_2 + m_2^2 + m_1 m_2)}{a_2(e_2 g_2 - m_2)} > \text{R1I}\left(\frac{a_1 m_2 S_1}{e_2 g_2 - m_2} - (\beta_1 + \beta_2 + m_1 + m_2)\right)$$

Subtracting $\text{R1I}\left(\frac{a_1 m_2 S_1}{e_2 g_2 - m_2} - (\beta_1 + \beta_2 + m_1 + m_2)\right)$ from the left hand side, some extensive rearranging yields:

$$H_1 (\beta_1 + m_1) - \frac{a_1 S_1 (a_2 H_1 + \beta_2 + m_2)}{g_2 + m_2} \frac{m_2(g_2 + m_2)}{a_2(e_2 g_2 - m_2)} - (\beta_1 + \beta_2 + m_1 + m_2) \frac{m_2(g_2 + m_2)}{a_2(e_2 g_2 - m_2)} +$$
$$a_2 H_1 \frac{m_2(g_2 + m_2)}{a_2(e_2 g_2 - m_2)} - \text{R1I} \frac{a_1 S_1}{m_2 + g_2} \frac{m_2(g_2 + m_2)}{a_2(e_2 g_2 - m_2)} + \text{R1I} (\beta_1 + \beta_2 + m_1 + m_2) > 0$$

Note that with the substitution of $g_2 = 1/\tau_2$, $R^*_{2,E} = \frac{m_2(g_2 + m_2)}{a_2(e_2 g_2 - m_2)}$. Using the simplified notation $R^*_{2,E}$ = R2E, R2E can be substituted several times in the above expression, which yields:

$$H_1 (\beta_1 + m_1) - \frac{a_1 S_1 (a_2 H_1 + \beta_2 + m_2)}{g_2 + m_2} \text{R2E} - (\beta_1 + \beta_2 + m_1 + m_2) \text{R2E} + a_2 H_1 \text{R2E} - \text{R1I} \frac{a_1 S_1}{m_2 + g_2} \text{R2E} + \text{R1I} (\beta_1 + \beta_2 + m_1 + m_2) > 0$$

Solving for R2E by adding $\left(\frac{a_1 S_1 (a_2 H_1 + \beta_2 + m_2)}{g_2 + m_2} + \frac{\text{R1I}(a_1 a_2 S_1)}{g_2 + m_2} + a_2 H_1 + (\beta_1 + \beta_2 + m_1 + m_2)\right)$ R2E from both sides and then dividing both sides by the terms in the parentheses (which does not flip the inequality because all terms are positive) yields:

**(* Do the above−described algebra *)**

$$\textbf{Simplify}\left[\frac{H_1 (m_1 + \beta_1) + \text{R1I} (\beta_1 + \beta_2 + m_1 + m_2)}{(\beta_1 + \beta_2 + m_1 + m_2) + a_2 H_1 + \frac{a_1 S_1 (a_2 H_1 + m_2 + \beta_2)}{g_2 + m_2} + \text{R1I}\left(\frac{a_1 S_1 a_2}{(m_2 + g_2)}\right)}\right] \text{// TraditionalForm}$$

$$\frac{H_1 (\beta_1 + m_1) + \text{R1I} (\beta_1 + \beta_2 + m_1 + m_2)}{\frac{a_1 S_1 (a_2 H_1 + \beta_2 + m_2)}{g_2 + m_2} + \frac{a_1 a_2 \text{R1I} S_1}{g_2 + m_2} + a_2 H_1 + \beta_1 + \beta_2 + m_1 + m_2}$$

for the left hand side of the equation (the right hand side remains R2E). With a little rearranging, this yields

$$\text{R2E} < \frac{1}{1 + \frac{a_2 H_1}{\beta_1 + \beta_2 + m_1 + m_2} + \frac{a_1 S_1}{g_2 + m_2}\left(\frac{a_2 (H_1 + \text{R1I})}{\beta_1 + \beta_2 + m_1 + m_2} + \frac{\beta_2 + m_2}{\beta_1 + \beta_2 + m_1 + m_2}\right)} \left(H_1 \frac{\beta_1 + m_1}{\beta_1 + \beta_2 + m_1 + m_2} + \text{R1I}\right)$$

Then, substituting $g_i = 1/\tau_i$ (i = 1,2):

$$R2E < \frac{1}{1 + \frac{a_2 H_1}{\beta_1+\beta_2+m_1+m_2} + \frac{a_1 S_1}{m_2+\frac{1}{\tau_2}}\left(\frac{a_2(H_1+\text{R1I})}{\beta_1+\beta_2+m_1+m_2} + \frac{\beta_2+m_2}{\beta_1+\beta_2+m_1+m_2}\right)}\left(H_1 \frac{\beta_1+m_1}{\beta_1+\beta_2+m_1+m_2} + \text{R1I}\right)$$

In the main text, we use the notation $\omega_{21} = \frac{\beta_1 + m_1}{\beta_1 + \beta_2 + m_1 + m_2}$ and $\omega_{12} = \frac{\beta_2 + m_2}{\beta_1 + \beta_2 + m_1 + m_2}$ in which $\omega_{21}$ and $\omega_{12}$ represent the probability that type 2 wins or loses a contest against type 1, respectively (see below for a justification). In addition, we define the term $P_2$

$$P_2 = \frac{1}{1 + \frac{a_2 H_1}{\beta_1+\beta_2+m_1+m_2} + \frac{a_1 S_1}{m_2+\frac{1}{\tau_2}}\left(\frac{a_2(H_1+\text{R1I})}{\beta_1+\beta_2+m_1+m_2} + \frac{\beta_2+m_2}{\beta_1+\beta_2+m_1+m_2}\right)}.$$

$P_2$ represents the negative effects of interference on an invader of consumer 2. The above invasion criterion can thus expressed as (substituting in full notation):

$$R^*_{2,E} < P_2 \left(H^*{}_1\, \omega_{21} + R^*{}_{1,I}\right) \qquad (4\,G)$$

which is identical to equation 10 in the main text, substituting 2 with $j$ and 1 with $i$.

Note that if consumer 2 never engages in interferences ($\beta_2 \to \infty$), then

```
(*Take limit as β2→∞*)
Simplify[Limit[1/(1 + (a2 H1)/(β1+β2+m1+m2) + (a1 S1)/(m2 + (1/τ2)) ((β2+m2)/(β1+β2+m1+m2) + (a2 (H1+R1I))/(β1+β2+m1+m2))), β2 → Infinity]] // TraditionalForm
```

$$\frac{m_2\,\tau_2 + 1}{a_1\,S_1\,\tau_2 + m_2\,\tau_2 + 1}$$

or, in more familiar form

$$P_2 \to \frac{m_2\,\tau_2+1}{a_1\,S_1\,\tau_2+m_2\,\tau_2+1} = \frac{1}{1+\frac{a_1\,S_1}{m_2+(1/\tau_2)}}$$

which is identical to $P'_d$ in case where one consumer (the "dove) never interferes (see Supplementary Materials F, Section 3-5).

### G.4 Coexistence Criteria

Consumer 2 invades when

$$R^*_{2,E} < P_2 \left(H^*{}_1\, \omega_{21} + R^*{}_{1,I}\right)$$

and, analogously, consumer 1 invades if

$$R^*_{1,E} < P_1 \left(H^*{}_2\, \omega_{12} + R^*{}_{2,I}\right)$$

To derive the mutual invasion criteria, we divide each side of the first inequality by $R^*{}_{1,E}$. For the first inequality, this gives

$$\frac{R^*_{2,E}}{R^*_{1,E}} < P_2\left(\frac{H^*{}_1}{R^*_{1,E}}\,\omega_{21} + \frac{R^*{}_{1,I}}{R^*_{1,E}}\right)$$

For the second inequality, we divide each side by $R^*{}_{1,E}\,P_1\,(\,R^*{}_{2,I}\; + \;H^*{}_2\,\omega_{12})$ and multiply each side by $R^*{}_{2,E}$. This yields

$$\frac{1}{P_1\left(\frac{H^*{}_2}{R^*_{2,E}}\,\omega_{12} + \frac{R^*{}_{2,I}}{R^*_{2,E}}\right)} < \frac{R^*_{2,E}}{R^*_{1,E}}$$

Putting these expression together, mutual  invasion thus occurs when

$$\frac{1}{P_1\left(\frac{H^*{}_2}{R^*_{2,E}}\,\omega_{12} + \frac{R^*{}_{2,I}}{R^*_{2,E}}\right)} < \frac{R^*_{2,E}}{R^*_{1,E}} < P_2\left(\frac{H^*{}_1}{R^*_{1,E}}\,\omega_{21} + \frac{R^*{}_{1,I}}{R^*_{1,E}}\right) \qquad (5G)$$

which is identical to equation 11 in the main text, substituting 2 with *j* and *i* with 1.

### G.5 Interpretation of $\omega_{21}$

In the main text, we claim that $\omega_{21}$ is the probability that consumer 2 wins a contest against consumer 1. That is,

$$\omega_{21} = \frac{\beta_1 + m_1}{\beta_1 + \beta_2 + m_1 + m_2} = \text{P[consumer type 2 wins a contest against type 1]} \qquad (6G)$$

To see this, recall that Ordinary Differential Equations (e.g., dN/dt = -k*N) are the deterministic mass action limit of exponentially distributed times to events. If two random variables, A and B, are exponentially distributed with rates a and b, P[A happens before B] = a/(a+b). Type j wins if type i gives up or dies before type i gives up or dies. Thus, the rate of j victory within a contest is $\beta_i + m_i$, and the rate of j loss is $\beta_j + m_j$. Note that this assumes m is identical between types. The interpretation in Equation 6G follows, where $\beta_i + m_i$ and $\beta_j + \beta_i + m_j + m_i$ are equivalent to a and a+b, respectively.

### G.6 Comment on $\beta_0$

In the main text, we refer to a "central" value of giving up rate, $\beta_0$. When consumers have identical exploitative ability, we state that $\beta_0$ represents a key transition such that coexistence always occurs if $\beta_i < \beta_0 <$ (see Fig. 2, main text). Here, we derive $\beta_0$.

Consider equation the invasion condition:

$$R^*{}_{2,E} \;<\; P_2\left(H^*{}_1\,\omega_{21} \;+\; R^*{}_{1,I}\right)$$

with some minor rearranging, type 2 invades if

$$f\,(\beta_1, \beta_2) = P_2\left(H^*{}_1\,\omega_{21} + R^*{}_{1,I}\right) - R^*{}_{2,E} > 0$$

Now, consider when $a_1 = a_2 = a$ and $\tau_1 = \tau_2 = \tau$ (consumers have identical exploitative ability). The invasion of j can be visualized as a "Pairwise Invasion Plot" or PIP used in adaptive dynamics (Brännström et al. 2013). Fig. G2 depicts the PIP, where $\beta_0$ is the gold star. $\beta_0$ corresponds to an evolutionary singular strategy (Brännström et al. 2013; Geritz et al. 1998), which for the value of $\beta_1$ for which

$$\frac{\partial f(\beta_1, \beta_2)}{\partial \beta_2} \Big|_{\beta_1=\beta_2} = 0$$

i.e., the value of $\beta_1$ for which partial derivative of $f(\beta_1, \beta_2)$ with respect to $\beta_2$ at the point $\beta_2 = \beta_1$ is equal to zero gives $\beta_0$. We show the derivation of $\beta_0$ below:

**(*Substitute all the equilibrium values, making exploitative parameters equal*)**

$$\mathbf{S1star} = \text{Limit}\left[-\frac{r(\beta_1+m)(a_1(k\,m\,\tau_1 - e\,k) + m(m\,\tau_1+1))}{a_1\,m(a_1\,k\,r\,\tau_1 + \beta_1 + m)}, \{a_1 \to a, \tau_1 \to \tau, a_2 \to a, \tau_2 \to \tau\}\right];$$

$$\mathbf{H1star} = \text{Limit}\left[-\frac{r\,\tau_1(\beta_1+m)(a_1(k\,m\,\tau_1 - e\,k) + m(m\,\tau_1+1))}{a_1(\beta_1(e - m\,\tau_1) + m(e + m\,r\,\tau_1^2 + \tau_1(r-m)))}, \{a_1 \to a, \tau_1 \to \tau, a_2 \to a, \tau_2 \to \tau\}\right];$$

$$\mathbf{R1Istar} = \text{Limit}\left[\frac{m(m\,\tau_1+1)(a_1\,k\,r\,\tau_1 + \beta_1 + m)}{a_1(\beta_1(e - m\,\tau_1) + m(e + m\,r\,\tau_1^2 + \tau_1(r-m)))}, \{a_1 \to a, \tau_1 \to \tau, a_2 \to a, \tau_2 \to \tau\}\right];$$

$$\mathbf{fBeta1Beta} = \text{Limit}\left[\frac{1}{1 + \frac{a_2\,H_1}{\beta_1+\beta_2+2\,m} + \frac{a_1\,S_1}{m + (1/\tau_2)}\left(\frac{\beta_2+m}{\beta_1+\beta_2+2\,m} + \frac{a_2(H_1+R1I)}{\beta_1+\beta_2+2\,m}\right)}\left(H_1\left(\frac{\beta_1+m}{\beta_1+\beta_2+2\,m}\right) + R1I\right) - \frac{m\,\tau_2+1}{\tau_2(a_1\,S_1+m)+1},\right.$$
$$\left.\{a_1 \to a, \tau_1 \to \tau, a_2 \to a, \tau_2 \to \tau\}\right];$$

**(* $f(\beta_1,\beta_2)$ calculation*)**
**fBeta1Beta = Limit[fBeta1Beta, {$S_1$ → S1star, $H_1$ → H1star, R1I → R1Istar}];**

**(*Take derivative of $f(\beta_1,\beta_2)$ with respect to $\beta_2$ and set $\beta_1 \to \beta_2$ *)**
**PartialDerivativefBeta1Beta = Simplify[Limit[D[fBeta1Beta, $\beta_2$], {$\beta_2 \to \beta_1$}]];**

**(*Solve for $\beta_0$ and print value*)**
**Simplify[Solve[PartialDerivativefBeta1Beta == 0, $\beta_1$]]**

$$\left\{\left\{\beta_1 \to \frac{1}{2(e - m\,\tau)}\left(m^2\,\tau(3 - r\,\tau) - m(-1 + 2\,e + r\,\tau) + \sqrt{m}\,\sqrt{1 + m\,\tau}\,\sqrt{m + 4\,a\,e\,k\,r\,\tau + m\,r^2\,\tau^2 - 2\,m\,r\,\tau(1 + 2\,a\,k\,\tau) + m^2\,\tau(-1 + r\,\tau)^2}\right)\right\},\right.$$
$$\left.\left\{\beta_1 \to -\frac{1}{2(e - m\,\tau)}\left(m^2\,\tau(-3 + r\,\tau) + m(-1 + 2\,e + r\,\tau) + \sqrt{m}\,\sqrt{1 + m\,\tau}\,\sqrt{m + 4\,a\,e\,k\,r\,\tau + m\,r^2\,\tau^2 - 2\,m\,r\,\tau(1 + 2\,a\,k\,\tau) + m^2\,\tau(-1 + r\,\tau)^2}\right)\right\}\right\}$$

where the positive root is:

$$\beta_0 = \frac{\sqrt{m}\,\sqrt{m\,\tau+1}\,\sqrt{4\,a\,e\,k\,r\,\tau - 2\,m\,r\,\tau(2\,a\,k\,\tau+1) + m^2\,\tau(r\,\tau-1)^2 + m\,r^2\,\tau^2 + m} - m(2\,e + r\,\tau - 1) + m^2\,\tau(3 - r\,\tau)}{2(e - m\,\tau)}$$

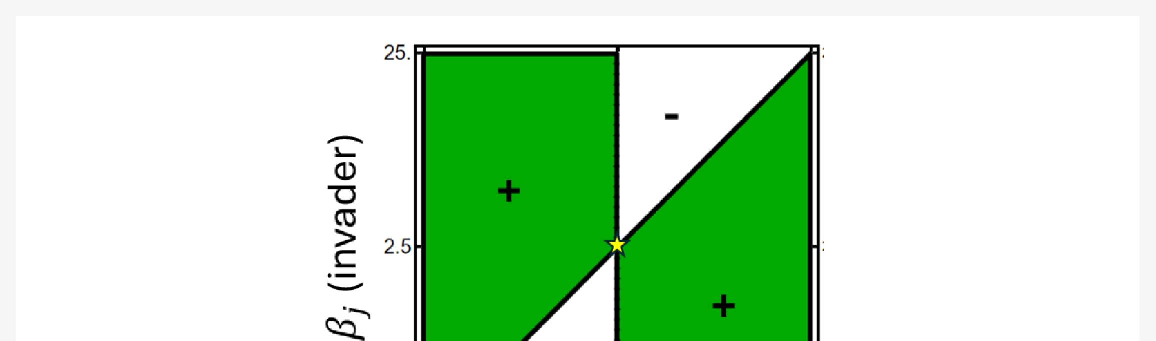

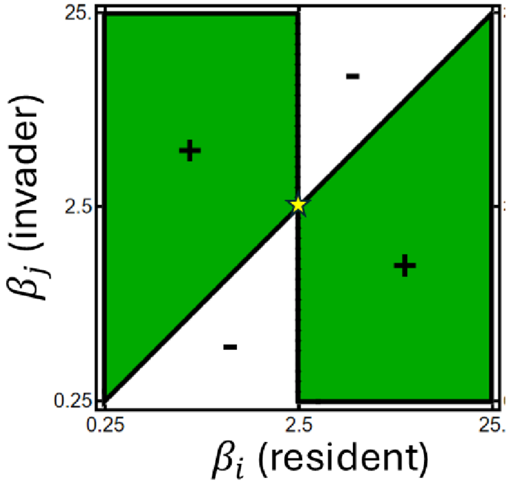


**Fig G2:** Pairwise Invasion Plot with respect to giving up rate $\beta$. The golden star is $\beta_0$. The invader $j$ with giving up rate $\beta_j$ invades against a resident with giving up rate $\beta_i$ in the green regions and fails to invade in the white regions. Parameters are the same as Fig. 3 in the main text.

### Section H: Invasion Criteria, Both Consumers Interfere

In the main text, we claim that priority effects are extremely rare in our model and ignore them on that basis. Here, we provide numerical evidence for this claim.

The lack (or possibility) of priority effects is traditionally demonstrated by performing a stability analysis of the 2-consumer equilibrium. However, we found that the equilibria of the full 8-equation system of differential equations (Equation (1) in the main text) were not analytically tractable. We therefore resort to numerical methods, described below.

##### H.1 Conditions for priority effects

We derive mutual invasibility condition in Section G (also see Supplementary Notebook C). Mutual invasion occurs if:

$$\frac{1}{I_{ij}} < E_{ji} < Ij_i \tag{1H}$$

where

$$I_{ij} = P_i\left(\frac{H_j^*}{R_{j,E}^*}\omega_{ij} + \frac{R_{j,I}^*}{R_{j,E}^*}\right)$$

$$E_{ij} = \frac{R_{j,E}^*}{R_{i,E}^*} \tag{2H}$$

$$I_{ji} = P_j\left(\frac{H_i^*}{R_{i,E}^*}\omega_{ji} + \frac{R_{i,I}^*}{R_{i,E}^*}\right)$$

$i, j = 1,2$, $i \neq j$. See Equation (11) in the main text and Section G for more details on each term. Conversely, a priority effect (founder control) would occur if

$$\frac{1}{I_{ij}} > E_{ji} > I_{ji} \tag{3H}$$

i.e., both consumers $i$ and $j$ are unable to invade the other when rare at the 1-consumer-1 resource equilibrium.

##### H.2 Numerical approach

We use a Latin Hypercube Sampling (LHS) approach. Using LHS, we numerically investigate a broad range of the parameters $a_i$, $a_j$, $\tau_i$, $\tau_j$, $\beta_i$, $\beta_j$, $e$, $m$, $r$, and $K$ (attack rates, handling times, giving up rates, conversion efficiency, mortality rate, resource renewal rate, and resource carry capacity, respectively). We $5 \times 10^7$ samples, sub-setting values that yield feasible 1-consumer-1-resource systems (i.e., $e_i > \tau_i m_i$ and $R_{i,E}^* < K$; see below equation (2) in the main text). See "Supplementary_Material_H_Priority_Effects_Code.R" for more details on the sampling.

For each sampled feasible set of parameter values, we calculated $I_j$, $E_{ji}$, and $I_i$. We then counted every parameter combination in which $1/I_j < E_{ji} < I_i$ (coexistence), $1/I_j > E_{ji} < I_i$ (type $i$ is excluded), $1/I_j < E_{ji} > I_i$ (type $j$ is excluded), and $1/I_j > E_{ji} > I_i$ (a priority effect).

##### H.3 The (near) absence of priority effects

We find that only coexistence or the exclusion of $i$ or $j$ occurs. Sampling over $> 10^7$ feasible parameter combinations, Equation 3H was fulfilled in approximately 0.003% of cases (Fig. 1H). Interestingly, these extraordinarily rare events are not a numerical artifact: ODE simulations confirm that priority effects indeed occur in these rarified parameter combinations.

Typically, we find priority effects tend to occur under extreme parameter combinations, such as when consumers differ in search rate $a$ by multiple orders of magnitude. Overall, priority effects are quantitively negligible and are unlikely to be biologically relevant where they do. We therefore ignore them in our analysis in the main text. All the parameter space examined in the main text yields no priority effects.

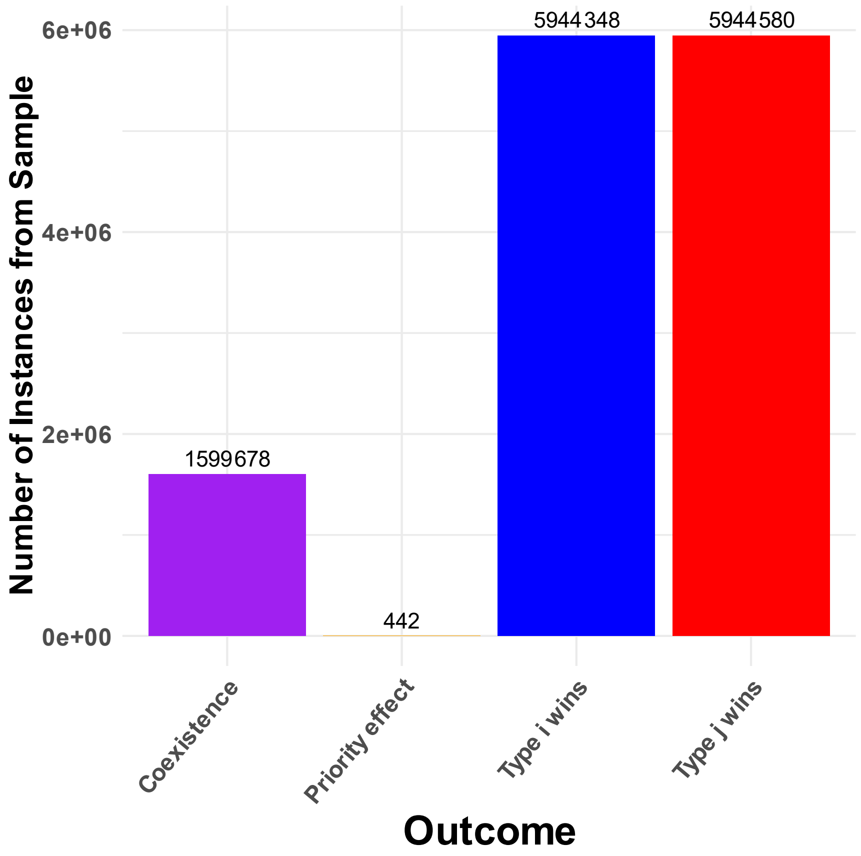


**Fig. H1:** priority effects are extremely rare. The bars show the frequency of each competitive outcome (coexistence, priority effect, type $i$ victory, and type $j$ victory) from the Latin Hypercube Sampling. We sampled $\sim 1.3 \times 10^7$ sets of parameters of the model for which 1-consumer-1-resource equilibria were feasible. Parameter were sampled using log-scaling as follows: $a_i \sim \{.01,100\}$, $a_j \sim \{.01,100\}$, $\tau_i \sim \{.001,100\}$, $\tau_j \sim \{.001,100\}$, $\beta_i \sim \{.01,1000\}$, $\beta_j \sim \{.01,1000\}$, $e \sim \{.01,100\}$, $m \sim \{.0001,10\}$, $r \sim \{.01,100\}$ and $K \sim \{5,5000\}$. For each set of parameter combinations from the Latin Hypercube Sampling, we calculated the values of the quantities in 2D, which were subsequently classified (see Table 1 of the main text). Modifying the parameter range of the sampling regime had negligible effects on the frequency of priority effects.

### Section I: Additional Supplementary Figures

We provide additional supplementary figures references in the main text. See the following pages.

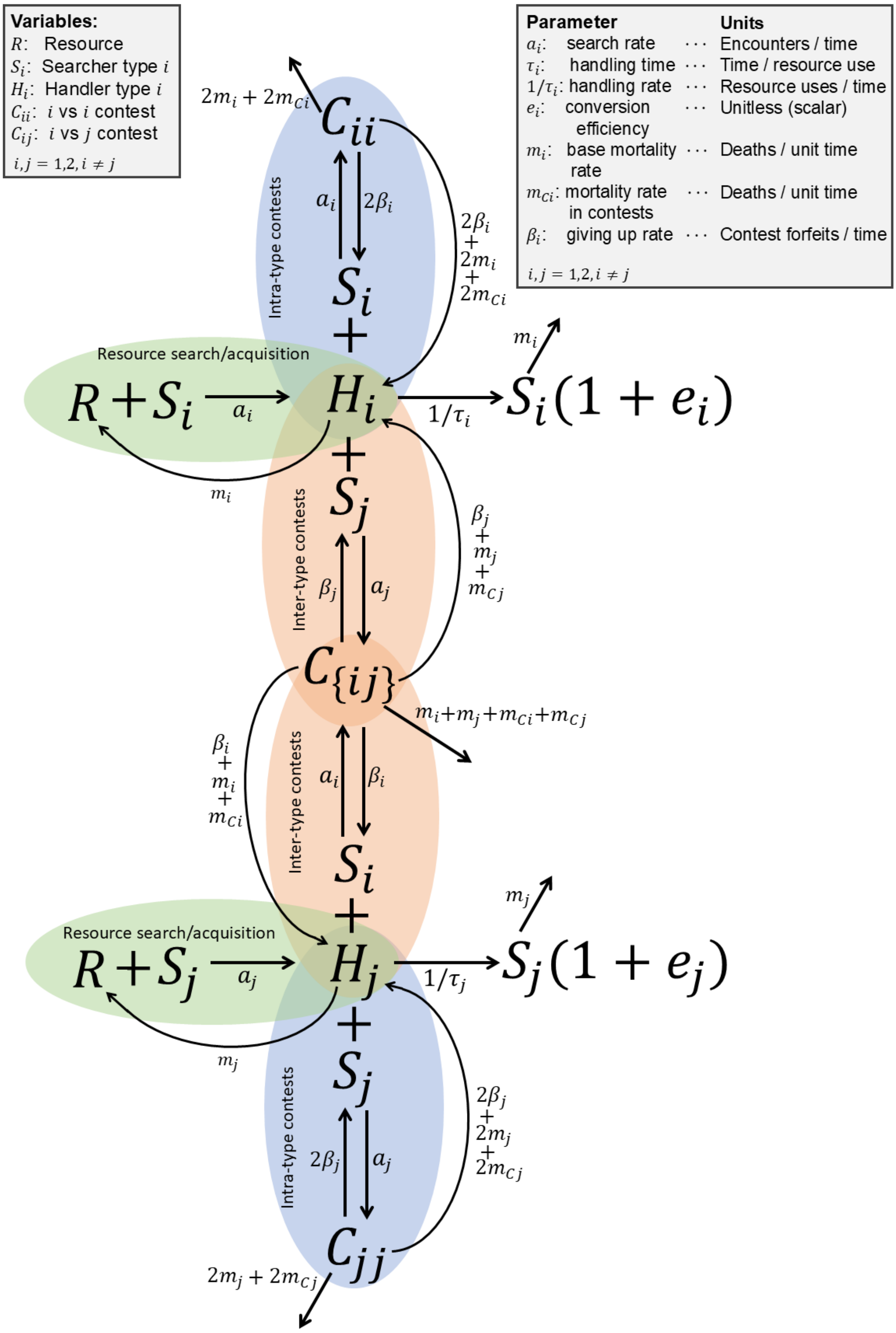


**Fig. I1:** Full depiction of the model, including species $i$ and $j$. See Figure. 1 and the main text for more details on the parameters and model.

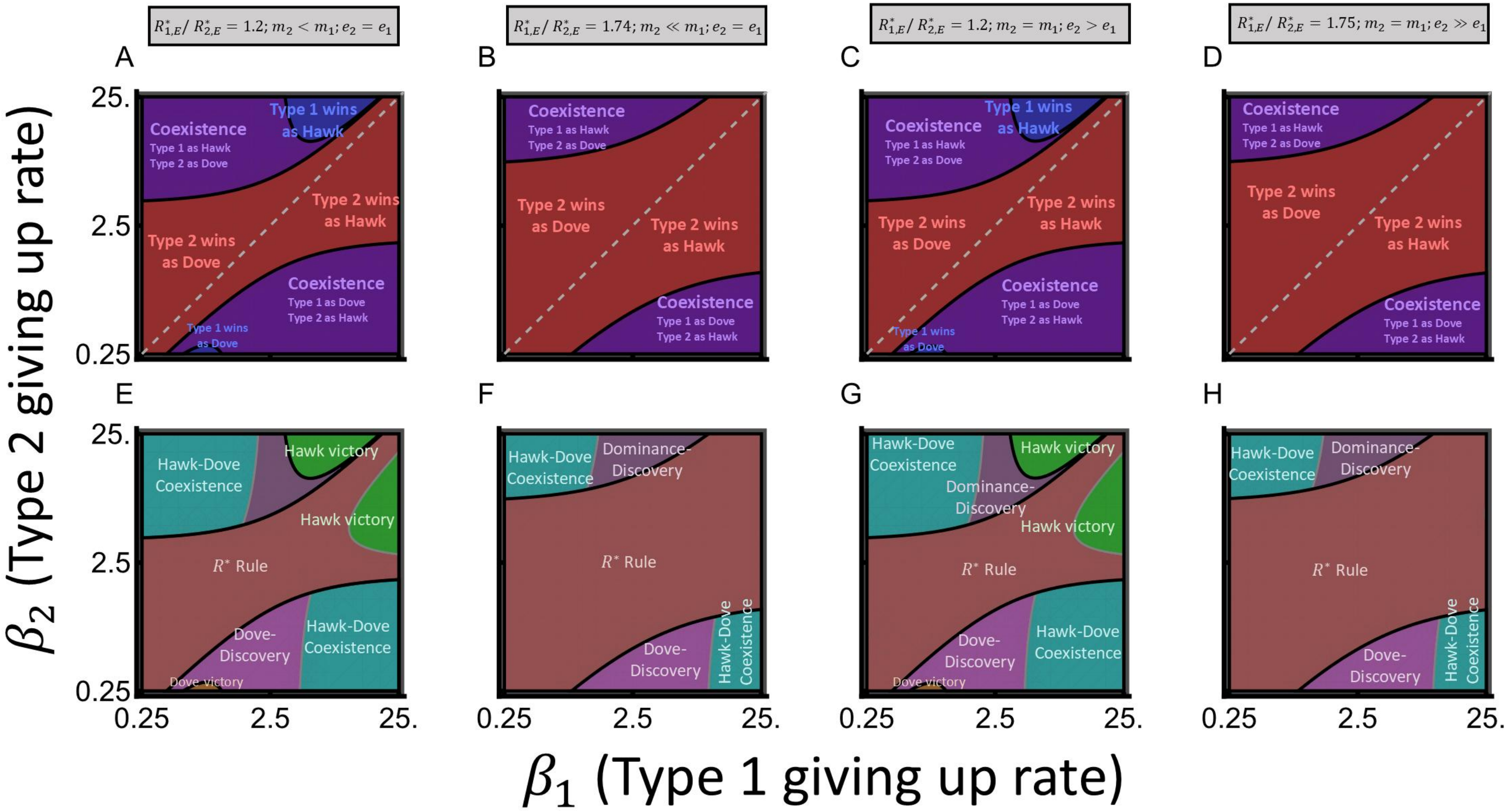


**Fig. I2:** Difference in exploitative ability driven by mortality rate $m$ and conversion efficiency $e$. In panels A and E, $m_1 = 0.015 > m_2 = 0.0135$. In panels B and F, $m_1 = 0.015 > m_2 = 0.0105$. In panels C and G, $e_1 = 0.035 < e_2 = 0.039$. In panels D and H, $e = 0.035 > e_2 = 0.05$. When not specified, $a_1 = a_2 = 0.1, \tau_1 = \tau_2 = 1, m_1 = m_2 = 0.015, e_1 = e_2 = 0.035, m_{C1} = m_{C2} = 0, r = 0.26, K = 250$. As compared to the cases in the main text, the $R^*$ drives most competitive exclusion.

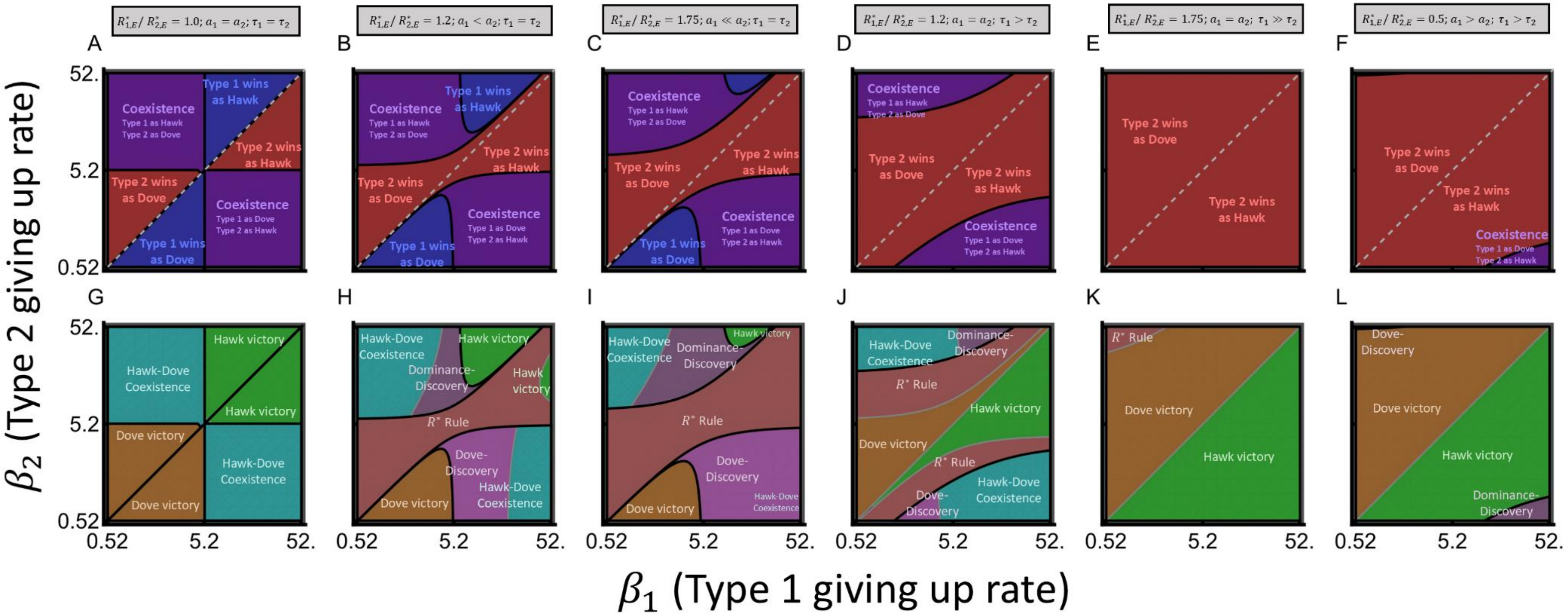


**Fig. I3:** The same as Fig. 4 from the main text, except $K = 1250$ instead of $K = 250$.

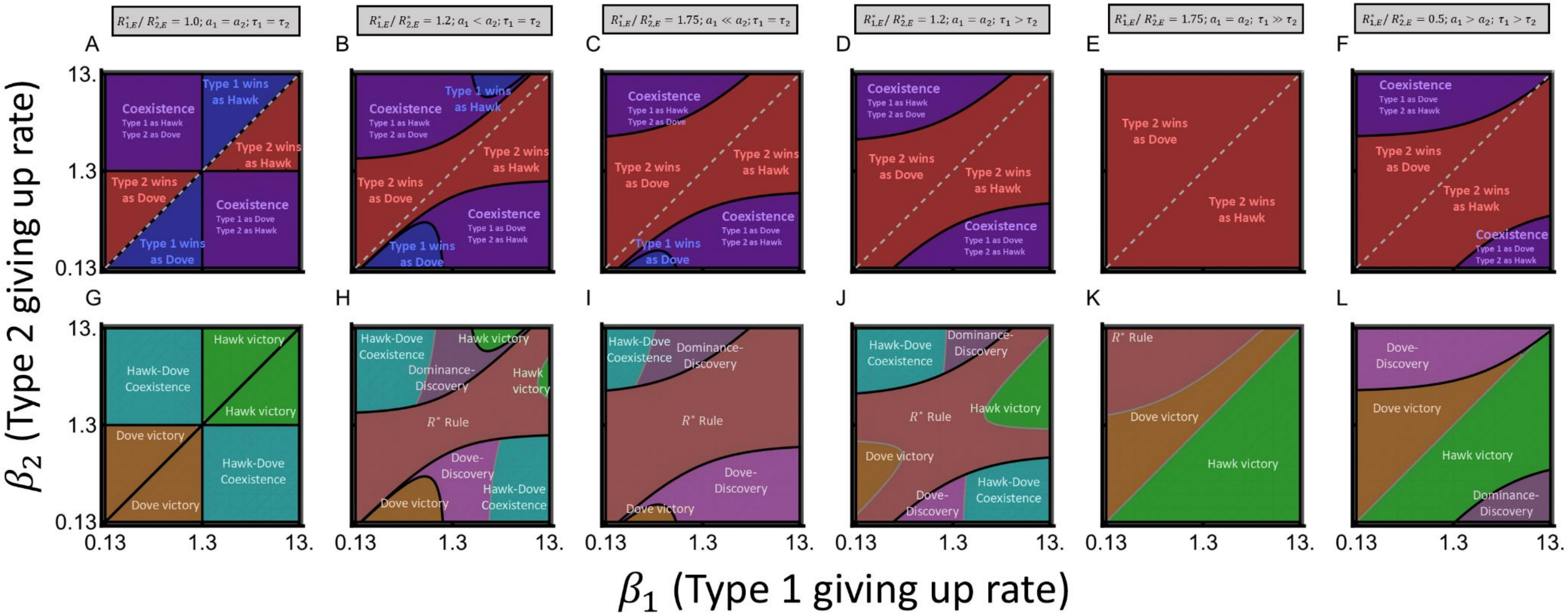


**Fig. I4:** The same as Fig. 4 from the main text, except $K = 50$ instead of $K = 250$.

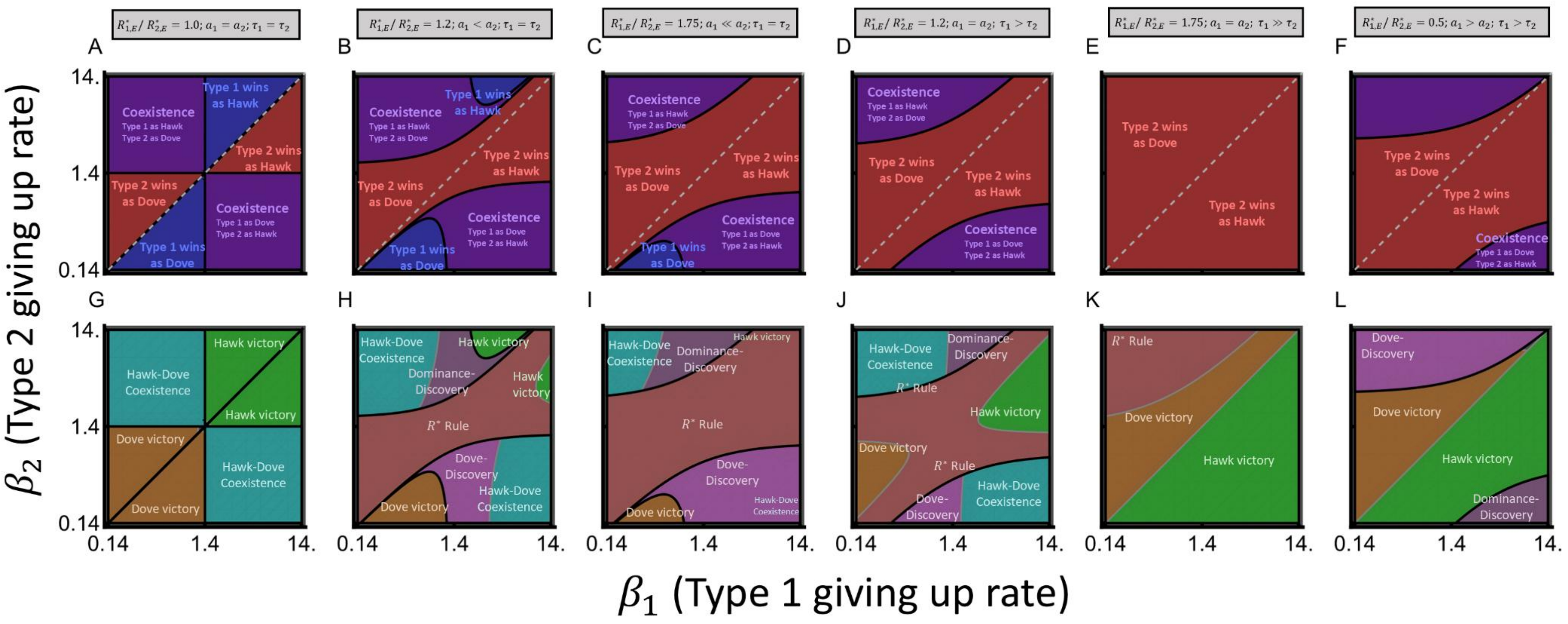


**Fig. I5:** The same as Fig. 4 from the main text, except $r = 0.052$ instead of $r = 0.26$.

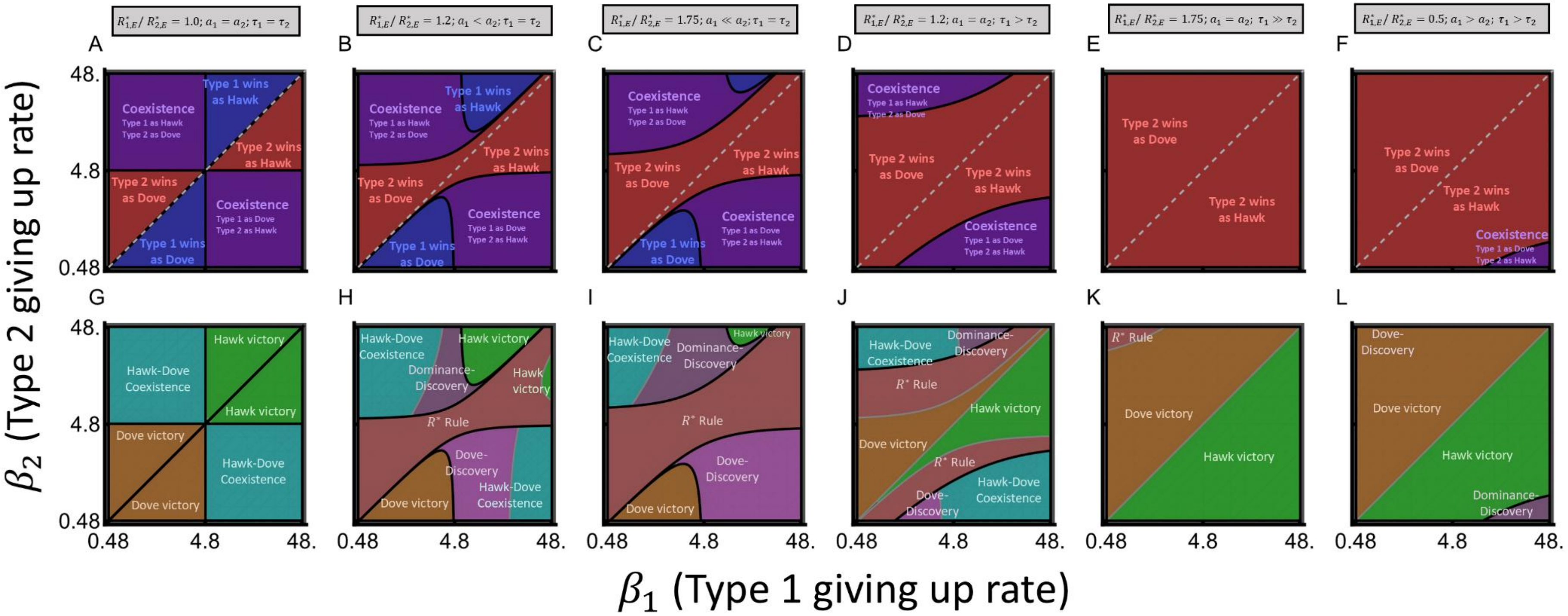


**Fig. I6:** The same as Fig. 4 from the main text, except $r = 1.3$ instead of $r = 0.26$.

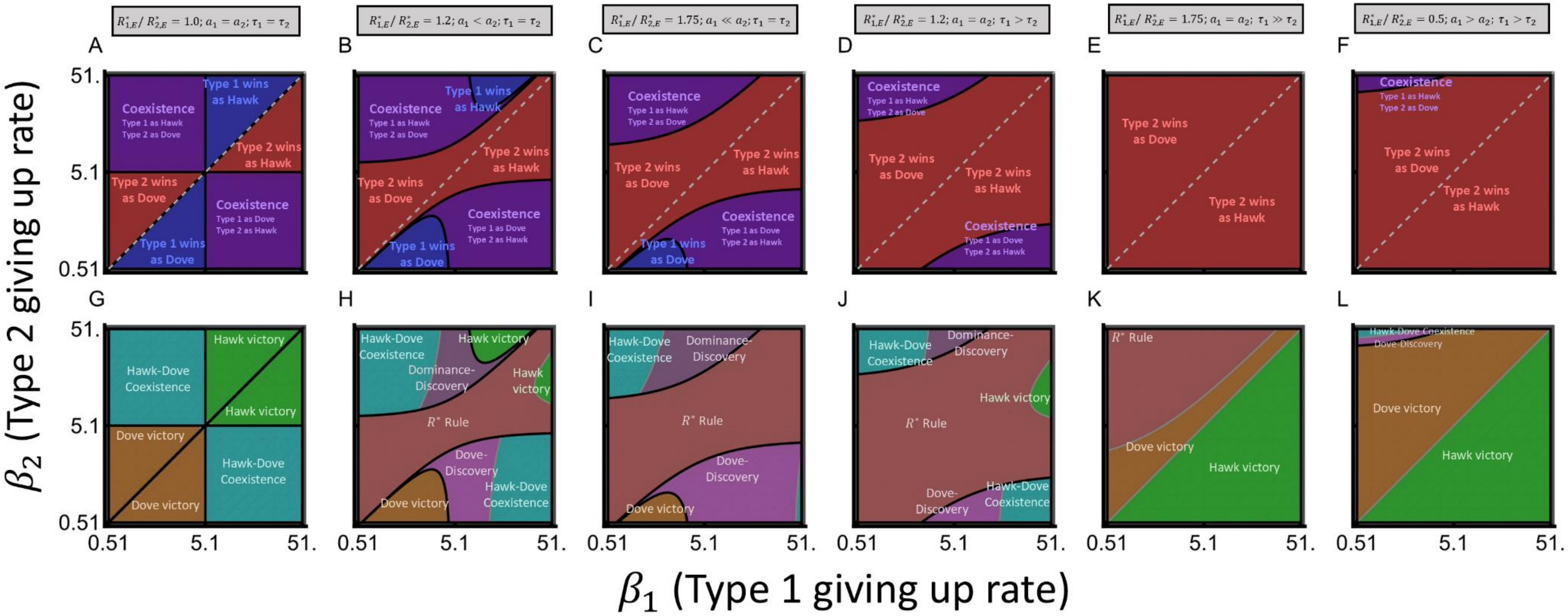


**Fig. I7:** The same as Fig. 4 from the main text, except $m = 0.025$ instead of $m = 0.015$.

\

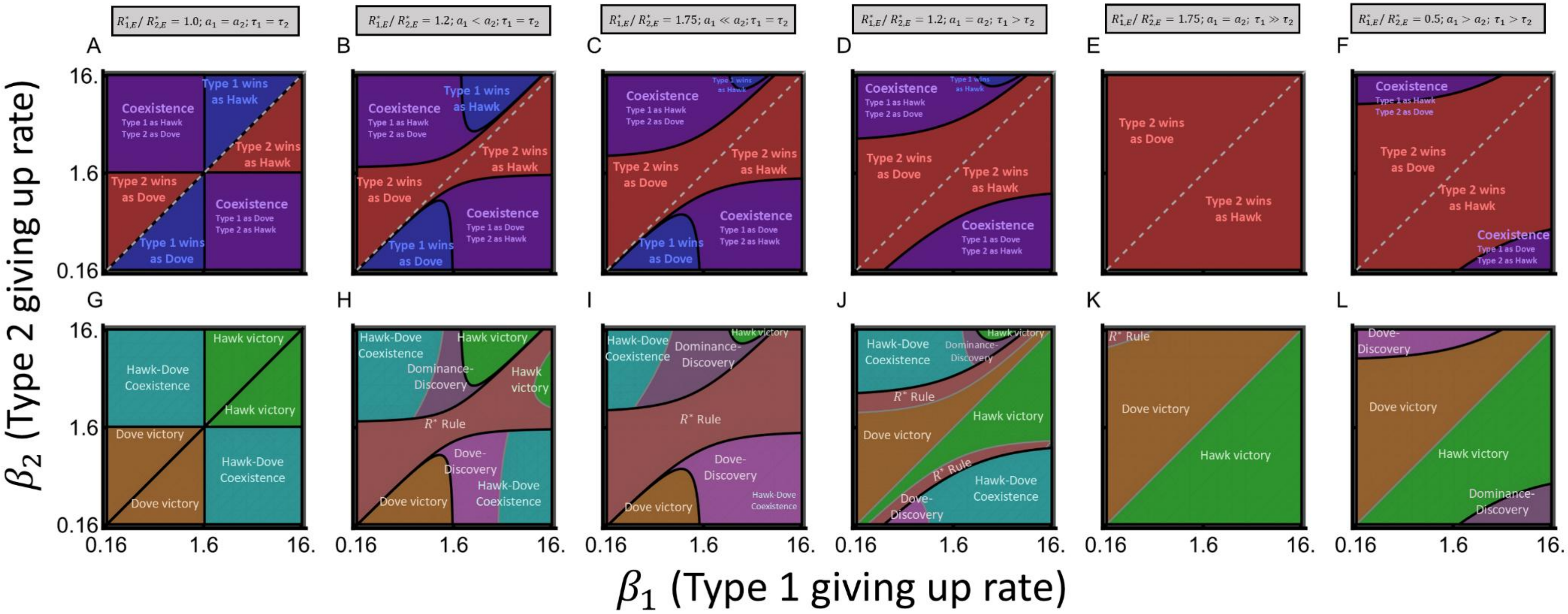


**Fig. I8:** The same as Fig. 4 from the main text, except $m = 0.009$ instead of $m = 0.015$.

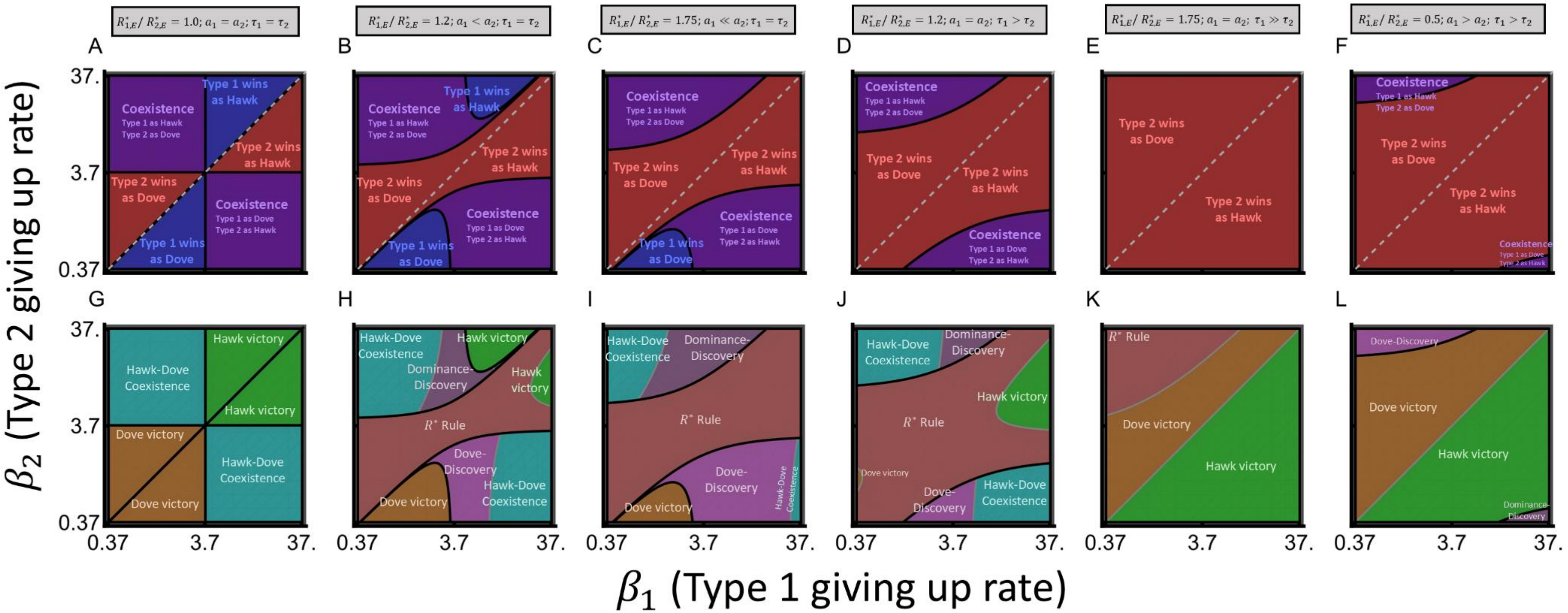


**Fig. I9:** The same as Fig. 4 from the main text, except $e = 0.025$ instead of $e = 0.035$.

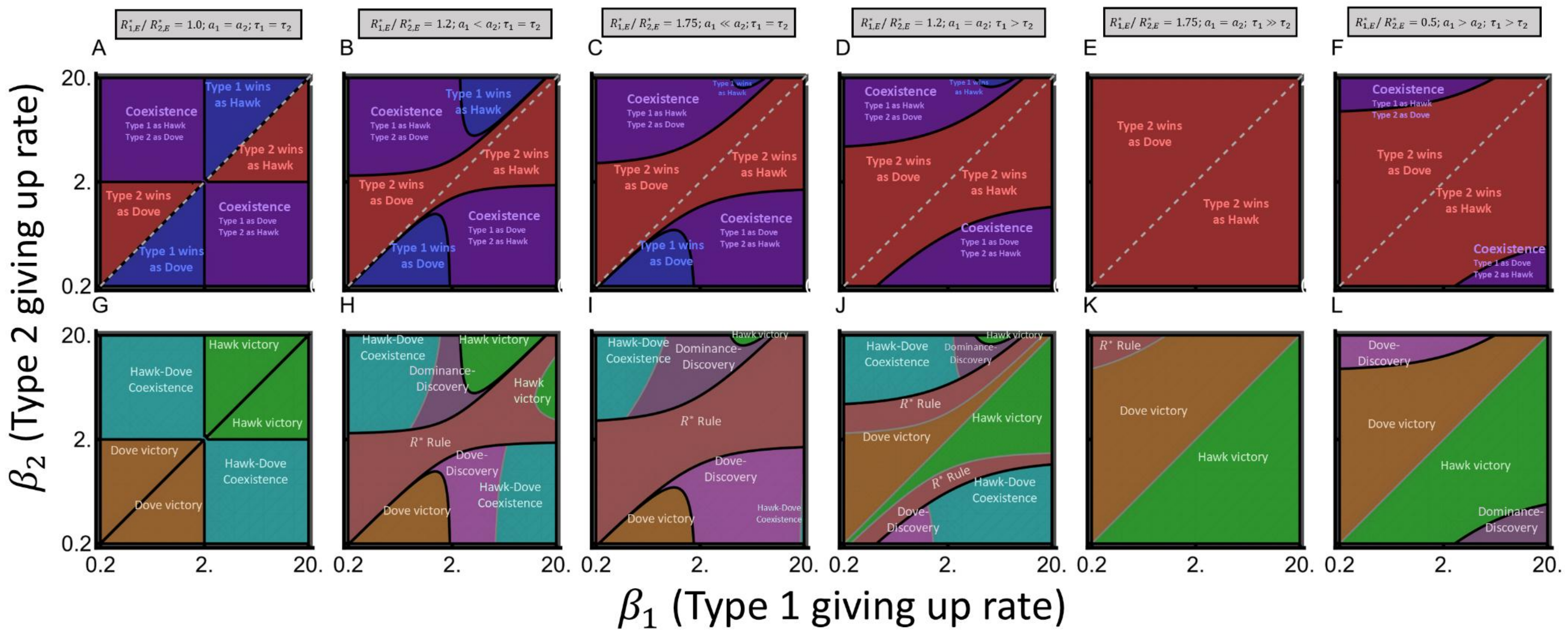


**Fig. I10:** The same as Fig. 4 from the main text, except $e = 0.045$ instead of $e = 0.035$.
